# LolA and LolB are conserved in Bacteroidetes and are crucial for gliding motility and Type IX secretion

**DOI:** 10.1101/2024.09.13.612696

**Authors:** Tom De Smet, Elisabeth Baland, Fabio Giovannercole, Julien Mignon, Laura Lizen, Rémy Dugauquier, Frédéric Lauber, Marc Dieu, Gipsi Lima-Mendez, Catherine Michaux, Damien Devos, Francesco Renzi

## Abstract

In Gram-negative bacteria, lipoproteins are major components of the outer membrane (OM) where they play a variety of roles, from the involvement in membrane biogenesis to virulence. Bacteroidetes, a widespread phylum of Gram-negative bacteria, including free-living organisms, commensals and pathogens, encode an exceptionally high number of outer membrane lipoproteins. These proteins are crucial in this phylum mainly because they are key components of SUS-like nutrient acquisition systems as well as of the Type 9 secretion (T9SS) and gliding motility machineries. The transport of lipoproteins to the OM has mainly been studied in *E. coli* and relies on the Lol system, composed of the inner membrane extraction machinery LolCDE, the periplasmic carrier LolA and the OM lipoprotein LolB. While most Lol proteins are essential and conserved across Gram-negative bacteria, to date, no LolB homologs have been identified outside of γ- and β-proteobacteria. How lipoproteins reach and are inserted in the OM of Bacteroidetes is not known. Here we identified LolB homologs in Bacteroidetes and disclosed the co-existence of several LolA and LolB in several species. We provide evidence that one LolA (LolA1) and one LolB (LolB1) of *F. johnsoniae* are devoted to targeting gliding and T9SS lipoproteins to the OM. A proteomic analysis of the OM composition of the *lolA1* and *lolB1* mutants supports this evidence. Furthermore, we show that, while LolB1 and LolA1 have conserved functions in Bacteroidetes, they are functionally different from their *E. coli* counterparts. We also show that surface lipoprotein transport is LolA and LolB independent. Finally, the finding that, in the absence of LolA and LolB homologs, lipoproteins still localize to the OM, suggests the presence in Bacteroidetes of yet unidentified LolAB-alternative lipoprotein transport pathways. In conclusion, Bacteroidetes have evolved different and more complex lipoprotein transport pathways than other Gram-negative bacteria and further research is required to uncover their complexity.

**Significance:** In Gram-negative bacteria, lipoproteins are key components of the outer membrane (OM), essential for functions like membrane biogenesis and virulence. Bacteroidetes, a widespread phylum, encode a high number of OM lipoproteins crucial for nutrient acquisition, Type IX secretion, and gliding motility. While lipoprotein transport in *E. coli* depends on the Lol system, LolB homologs were previously unidentified outside γ- and β-proteobacteria. Here we identify LolB homologs in Bacteroidetes, revealing the co-existence of multiple LolA and LolB proteins in various species. In *F. johnsoniae*, LolA1 and LolB1 specifically target gliding and Type 9 secretion system lipoproteins to the OM. Despite this, lipoproteins still localize to the OM without LolA and LolB, suggesting alternative transport pathways. These findings indicate that Bacteroidetes have evolved more complex lipoprotein transport mechanisms than other Gram-negative bacteria, requiring further research to fully understand them.

## Introduction

The cell envelope of Gram-negative bacteria is composed of two membranes, the inner membrane (IM) delimiting the cytoplasm, and the outer membrane (OM) delimiting the periplasm. Residing in between these two membranes is an additional thin layer made of peptidoglycan. Unlike the IM, composed exclusively of phospholipids, the OM is an asymmetric bilayer, with phospholipids on the inner and lipopolysaccharide (LPS) on the outer leaflet, forming a dense network impermeable to most compounds (1). Import of nutrients through the OM thus relies on integral outer membrane proteins called porins, water-filled channels that allow substrates to cross the OM. In addition, lipoproteins, proteins with an acylated N-terminus allowing their anchorage in the membrane, are found in both membranes and participate in a multitude of functions including nutrient acquisition, stress sensing, cell morphology, transport etc (1). In *E. coli*, the most abundant lipoprotein is Lpp, inserted into the inner leaflet of the OM, which anchors the peptidoglycan to the OM thus playing a crucial role in the envelope architecture (2). OM lipoproteins are also part of membrane protein complexes crucial for envelope maturation and cell viability such as the BAM complex, responsible for β-barrel protein insertion and folding into the OM (3) and the Lpt complex involved in LPS transport across the OM (4). While in most studied bacteria OM lipoproteins are mainly anchored facing the periplasm, it has recently been shown that surface-exposed lipoproteins are more widespread than initially thought especially in bacteria of the phylum Bacteroidetes where they are crucial for nutrient acquisition (5–7). Furthermore, OM lipoproteins are key elements of the gliding and Type 9 secretion (T9SS) systems, two intertwined machineries which are a hallmark and unique to Bacteroidetes (8). Despite different localizations, lipoproteins share a common maturation pathway. They are synthesized in the cytoplasm and after translocation to the periplasm, mainly through the Sec machinery, their N-terminal cysteine is acylated by the action of three IM enzymes. Once lipidated, lipoproteins remain either in the IM or are targeted to the OM (9). The periplasm is a hydrophilic aqueous environment, yet, lipoproteins being hydrophobic, as most OM components, they require a specific transport system (1, 10).

In *E. coli*, lipoproteins are transported to the OM thanks to the Lol system, composed of five proteins: the LolCDE *ATP*-binding cassette (ABC) transporter, responsible for lipoprotein extraction from the IM; the periplasmic chaperone LolA, that transports lipoproteins through the periplasm and delivers them to the OM lipoprotein LolB that inserts them in the OM (11). Strikingly, while the components of the Lol machinery are essential for cell viability in *E. coli*, they are only partially conserved throughout Gram-negative bacteria. Indeed, LolB seems to be present only in β- and γ-Proteobacteria (12), while a bifunctional LolA, capable of both chaperoning and inserting lipoproteins in the OM, has recently been identified in α- Proteobacteria and Spirochetes (13, 14). In addition, bacteria such as Spirochetes and Bacteroidetes abundantly expose lipoproteins on their surface (5, 9, 15).

In Bacteroidetes, lipoproteins are targeted to the surface by an N-terminal lipoprotein export signal (LES), however, how they cross the OM is still unknown (16, 17). In Spirochetes, such an export signal could not be identified but lipoprotein surface localization is dependent on an Lpt-like pathway where a homolog of the LPS transporter LptD is responsible lipoprotein surface translocation (15). It remains therefore unknown how Bacteroidetes insert lipoproteins into their OM and how they flip some of them to the surface.

Here, by *in silico* distant homolog prediction we identify LolB homologs in bacteria of the phylum Bacteroidetes and demonstrate that multiple LolA and LolB proteins co-exist within the same Bacteroidetes species.

We show that in the Bacteroidetes *Flavobacterium johnsoniae* that possesses three LolA and two LolB homologs, one LolA and one LolB are involved in lipoprotein trafficking and their deletion determines lack of gliding and T9 protein secretion as well as severe membrane alterations. Strikingly, despite the presence of several LolA and LolB homologs, we additionally provide evidence that surface lipoprotein transport can work independently of these proteins in *F. johnsoniae*. Finally, the finding that *F. johnsoniae* can survive in the absence of all LolA and LolB proteins strongly supports the hypothesis that another pathway is responsible for or can overtake OM lipoproteins transport in the absence of LolA and LolB in Bacteroidetes.

## Results

### *Flavobacterium johnsoniae* possess several LolA and LolB

While the presence of LolA homologs has been reported in Bacteroidetes, no LolB homologs has ever been identified in bacteria of this Phylum by sequence similarity searches. We started from the hypothesis that Bacteroidetes might encode remote LolB homologs with low sequence identity but that would share a similar structure. To identify such homologs, we performed an *in silico* prediction analysis searching for remote homologs of *E. coli* LolB in *F. johnsoniae* (see methods for more details).

Surprisingly, this analysis identified two LolB homologs in *F. johnsoniae*: Fjoh_1066 (LolB1) and Fjoh_1084 (LolB2). Both candidates are predicted to encode a SPII signal sequence, in accordance with LolB being a lipoprotein. We performed the same analysis to search for *E. coli* LolA homologs and surprisingly three proteins were identified: the two LolA already found by sequence similarity and reported in the literature (Fjoh_2111, LolA1, and Fjoh_1085, LolA2) (18, 19) and a third one, Fjoh_0605, LolA3. LolA is a periplasmic carrier and thus harbors a SPI signal allowing it to cross the IM via the Sec pathway and reach the periplasm (20). All three LolA homologs carry such a signal sequence and thus likely localize in the periplasm.

Interestingly, while genes *Fjoh_2111* (*lolA1*), *Fjoh_0605* (*lolA3*) and *Fjoh_1066* (*lolB1*) are encoded in different genomic loci, genes *Fjoh_1084* (*lolB2*) and *Fjoh_1085* (*lolA2*) are part of an operon involved in flexirubin synthesis and transport, thus suggesting their implication in biosynthesis of this pigment (19).

Next, we modelled the *F. johnsoniae* LolA and LolB protein structures and compared their overall secondary and tertiary structures with respect to crystal structures of their *E. coli* homologs (**Figure 1**). *F. johnsoniae* LolA and LolB variants retain the main features of the *E. coli* unclosed β-barrel presenting a convex and a concave side, composed of 11 antiparallel β-strands in which helices (α- and/or 3_10_-helices) are embedded (**Figure 1**), and enclosed by an N-terminal α-helix. Among the LolA proteins, *E. coli* <u>LolA</u> (LolA*_Ec_*) and LolA1 have the same mainly disordered C-terminal region possessing one short 3_10_-helix as well as one extra parallel β-strand, while LolA2 and LolA3 have a shorter and longer C-terminal end, respectively. In addition, the *E. coli* LolB (LolB*_Ec_*) specific feature, consisting in a protruding loop comprising an exposed hydrophobic amino acid, is conserved in LolB1 and LolB2 (**Figure 1**). Interestingly, in LolB*_Ec_* and LolB1 this is a leucine, while in LolB2 a phenylalanine. LolB2 is structurally closer to LolB*_Ec_* than LolB1 which has a long N- and C-terminal disordered tail, as well as a very long loop extension compacted beneath the barrel.

**Figure 1.**
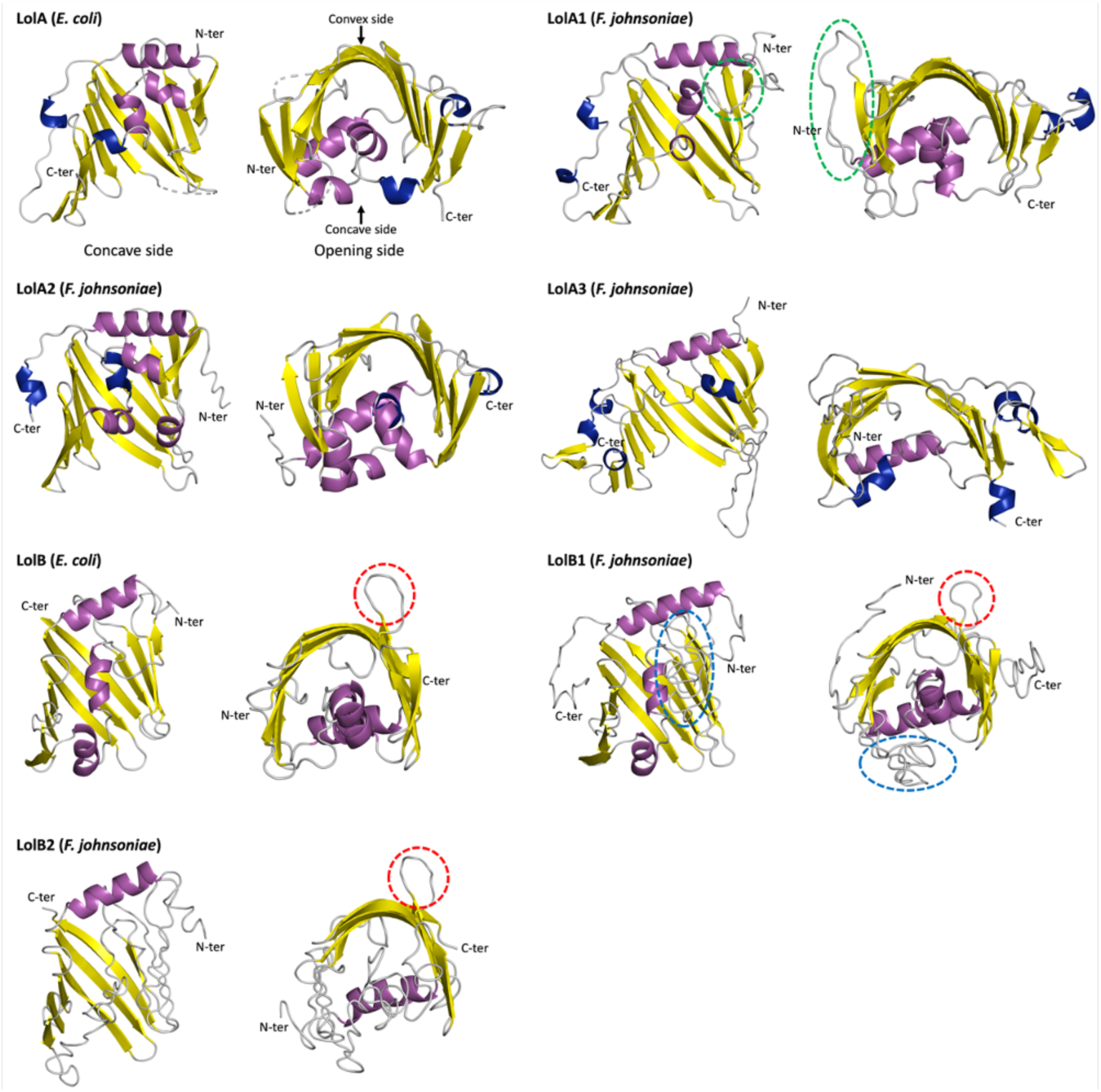
Structural comparison between *E. coli* and *F. johnsoniae* LolA and LolB homologs. Comparison of crystallized *E. coli* LolA (PDB entry: 1UA8) and LolB (PDB entry: 1IWM) structures with models of LolA1, LolA2, LolA3, LolB1, and LolB2 homologs from *F. johnsoniae*. Each protein is shown as a cartoon representation and secondary structure elements are colored as follows: α-helix (purple), 3_10_-helix (blue), β-sheet (yellow), and coil (grey). On each structure, the N- and C-terminal positions are indicated. In LolA1, the loop extension between α-helix 3 and β-strand 7 is highlighted with a dashed green ellipse. In LolB, LolB1, and LolB2, the upward conserved loop is highlighted with a dashed red circle. In LolB1, the compacted loop extension between α-helix 3 and β-strand 7 is highlighted with a dashed blue ellipse.

The mechanism of transfer of lipoproteins from LolA*_Ec_* to LolB*_Ec_* occurs via a complex formation through electrostatic potential complementarity, forming a contiguous hydrophobic surface, *i.e.* a tunnel-like structure, composed of both LolA and LolB concave sides. The lipoprotein is transferred due to a higher affinity of its lipid moiety for LolB than LolA due to a hydrophobic gradient between the cavities of the two partners (21). LolA*_Ec_* concave side, especially along the edge of the β-barrel opening, is enriched in negatively charged amino acids, interacting with the LolB*_Ec_* convex side mainly positively charged (21). With this mechanism in mind, we aimed at comparing the hydrophobicity (**Supplementary Figure S1**) and charge state (**Supplementary Figure S2**) of *F. johnsoniae* LolA and LolB homologs to highlight differences and, potentially predict LolA-LolB interactions.

An interesting tendency is the distinct enrichment in highly hydrophobic (leucine and isoleucine) amino acids highlighted for all the *F. johnsoniae* LolA and LolB homologs (**Supplementary Table S1**). Consequently, these are overall more hydrophobic than LolA*_Ec_* and LolB*_Ec,_* and LolA2 stands out as the most hydrophobic one. Furthermore, LolB1 contains a higher proportion of hydrophobic residues than LolA1, suggesting a favorable hydrophobic gradient for the lipoprotein transfer between LolA1 and LolB1. Although LolB2 has less hydrophobic residues than LolA2, its higher content in leucine and their local concentration within the concave side of the β-barrel could nonetheless allow the lipoprotein transfer toward LolB2 (**Supplementary Figure S1**). The same rationale can be applied to the putative LolA1-LolB1 couple. In the case of LolA3, the side chains pointing toward the inside of the concavity are substituted in tyrosine and isoleucine, increasing the overall polarity of the binding site.

Regarding the electrostatic driven LolAB paired complexes formation, a slight decrease in arginine compensated by a large increase in the lysine content is observed for all the *F. johnsoniae* proteins (**Supplementary Table S1**), especially for LolA1 and LolA2, changing the concave side charge from negative to mostly positive (**Supplementary Figure S2**). LolA3 seems closer to LolA*_Ec_*, with a predominance of aspartate and glutamate residues in the concave side, giving it a global negative charge. Most remarkably, its unique long and disordered C-terminal region bears a high positive charge, due to a dense population of lysine. Furthermore, while the positive character of the LolB convex side is mainly conserved in its *F. johnsoniae* homologs, a more evocative negative charge is exhibited by the concave side, particularly at the level of the loop extension of LolB1, absent in LolB*_Ec_* and LolB2 (**Supplementary Figure S2**). Therefore, the mechanism and preferential orientation by which LolA-LolB complexes are formed in *F. johnsoniae* most likely differ from that of *E. coli*, and pairs selectivity will also arise from subtle modifications in hydrophobicity and charge distribution.

### Deletion of *lolA1* and *lolB1* affects gliding motility and Type IX secretion

Interestingly, in a transposon mutagenesis screening aiming at identifying genes involved in gliding motility, a *F. johnsoniae* non-gliding mutant with a transposon insertion in *Fjoh_2111* (*lolA1*) was isolated (18). Since LolA allows lipoproteins to cross the periplasm and localize to the OM, the lack of gliding of the Tn mutant could be due to mislocalization and consequent depletion of lipoproteins in the OM. To confirm the absence of motility reported for a *lolA1* mutant and to determine whether deletion of any of the other *F. johnsoniae* LolA and LolB proteins could have a similar effect, we assessed the gliding motility of the *lolA1*, *lolA2*, *lolA3*, *lolB1* and *lolB2* mutants on plates. As expected, *lolA1* deletion resulted in non-spreading colonies and, interestingly, a lack of motility comparable to that of a *gldJ* non-gliding mutant was also observed when *lolB1* was deleted (23) (**Figure 2A**). GldJ is an OM lipoprotein required for both gliding and T9 secretion (23, 24). Deletion of *lolA2*, *lolA3* and *lolB2* did not affect gliding. Co-deletion of both *lolA1* and *lolB1* resulted in the same phenotype as the single *lolA1* mutant. Plasmid-borne complementation of *lolA1* and *lolB1* deletion with *lolA1* and *lolB1* fully restored gliding motility (**Figure 2A**).

**Figure 2.**
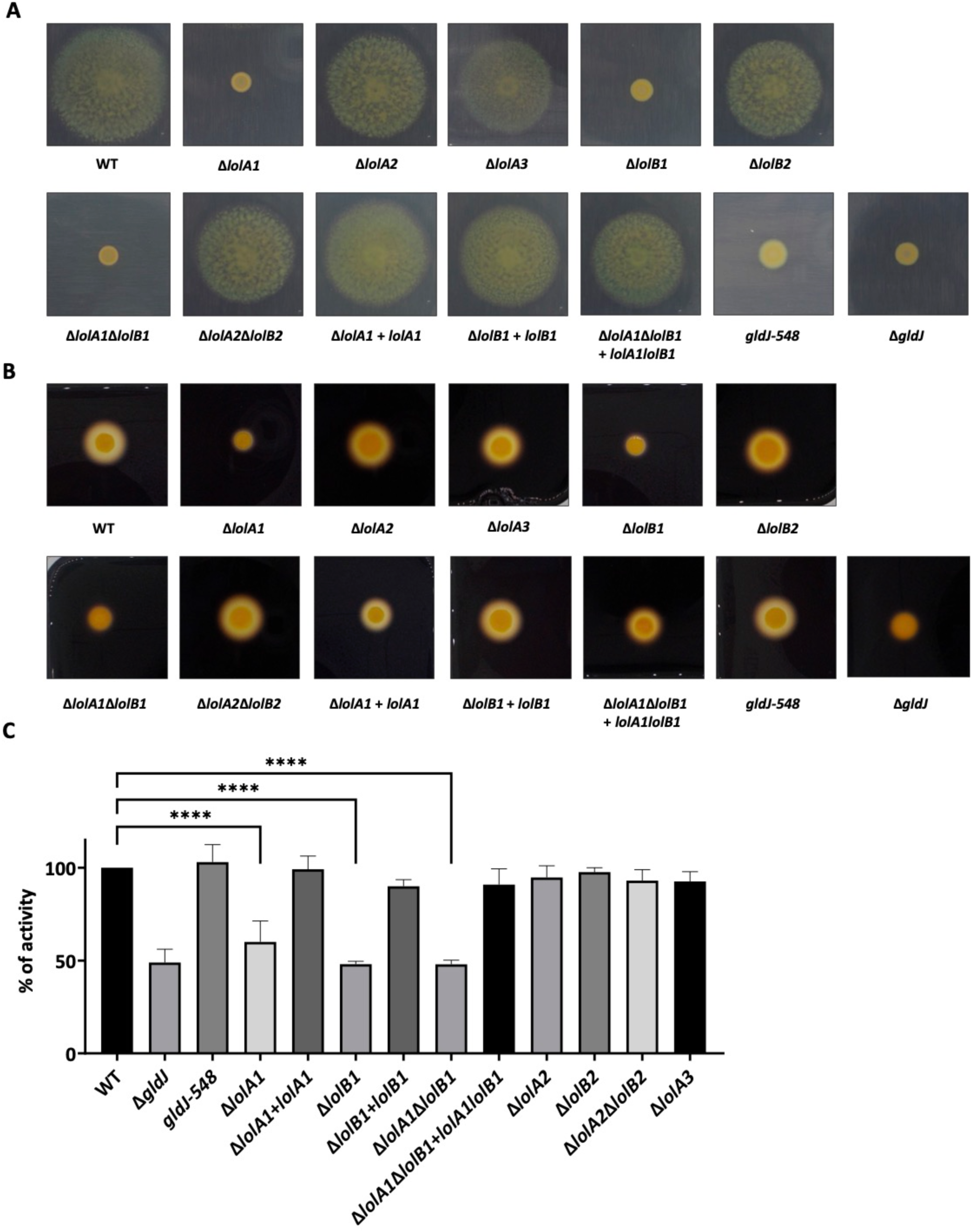
Deletion of *lolA1* and *lolB1* affects gliding motility and Type IX secretion. Gliding motility on MM agar plates after 48h (**A**). Starch degradation by T9-secreted amylases on LB starch plates after 24h (**B**) and amylase activity of cell culture supernatants (**C**). Data is represented as means of 3 biological replicates ± standard deviation, and statistical analysis was a one-way ANOVA followed by Dunnett’s multiple comparison test (F-value = 47.47; DF = 22). Asterisks indicate the following: (****): p<0.0001.

Since gliding motility and T9 secretion are highly interlinked and both require several OM lipoproteins for their activity (8, 22), we next determined the impact of *lolA* and *lolB* homologs deletion on T9SS activity. This secretion system is unique to Bacteroidetes and is involved in the secretion of various proteins mainly involved in adhesion and nutrient acquisition, such as α-amylases that allow *F. johnsoniae* to hydrolyze and feed on starch (24). To monitor Type 9 secretion, we measured the starch degradation activity of secreted amylases on starch containing plates and on cell culture supernatants. Starch degradation was significantly reduced in the *lolA1*, *lolB1* and *lolA1lolB1* mutant strains compared to the WT and like that of the *gldJ* non-gliding and non-secreting mutant (23) (**Figure 2B** and **2C**). Deletion of *lolA2*, *lolA3*and *lolB2* did not affect amylase secretion (**Figure 2B** and **2C**). Plasmid-borne complementation of *lolA1* and *lolB1* strains fully restored amylase secretion (**Figure 2B** and **2C**). The reduced amylase activity observed for the *lolA1* and *lolB1* mutants could not be ascribed to the lack of motility of these strains as a *gldJ-548* mutant, which is non-motile but has an active T9SS (8), degraded starch to the same extent as the WT strain (**Figure 2B** and **2C**).

Overall, these data indicate that LolA1 and LolB1 are crucial for both gliding motility and T9 secretion and reinforce our initial hypothesis that these two proteins might belong to a same pathway, namely the transport of lipoproteins to the OM. Given their pivotal role in *F. johnsoniae*, we focused on further characterizing the function of these two proteins.

### Deletion of *lolA1* and *lolB1* affects outer membrane proteome composition

Absence of gliding and T9 secretion could be due to mislocalization of OM lipoproteins crucial for these pathways in the absence of LolA1 and LolB1. To test this hypothesis, we determined the impact of *lolA1* and *lolB1* deletion on the OM protein composition of bacteria grown in CYE rich medium. We purified the OM of WT, *lolA1*, and *lolB1* strains and identified proteins by mass spectrometry. Overall, the absence of *lolA1* or *lolB1* respectively affected 610 and 172 proteins in a significant manner (Fold Change ≥ 1.5; Significance ≥ 20) (**Figure 3A** and **Supplementary Table S2**). We sorted the proteins according to their localization: cytoplasmic (no SP), integral OM and soluble periplasmic proteins (SPI), periplasmic-facing lipoproteins (SPII), and surface-exposed lipoproteins (SPII-LES). The OM proteome of the *lolB1* mutant was enriched in membrane proteins and lipoproteins while, to our surprise, that of the *lolA1* mutant contained many cytoplasmic proteins, accounting for more than 60 % of significant entries (**Figure 3A**). We clustered proteins harboring an SP by predicted function and mapped those involved in polysaccharide utilization to their respective(s) PUL(s) (25).

**Figure 3.**
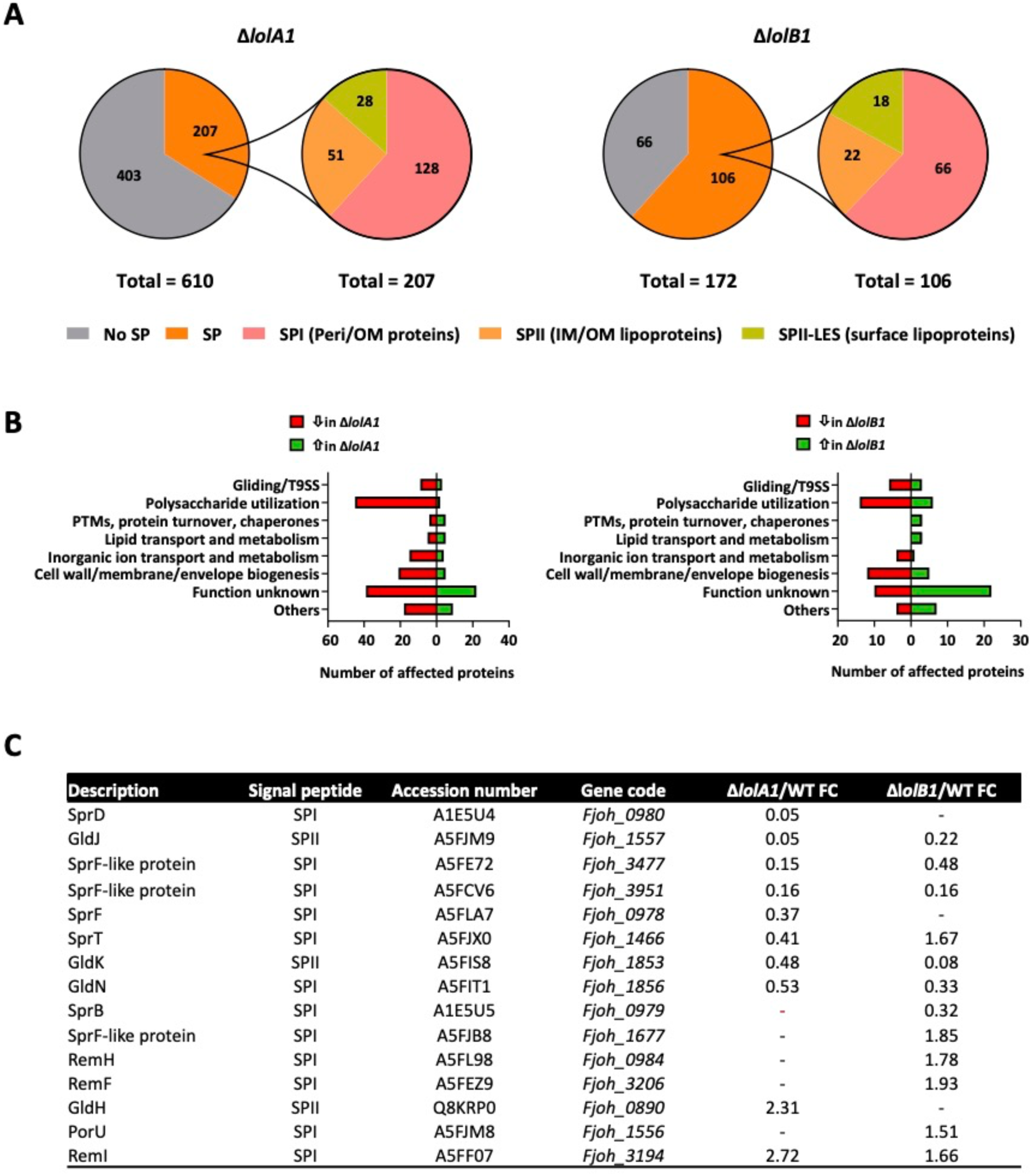
The OM protein composition of *lolA1* and *lolB1* mutants is altered compared to the WT. Proteins identified by mass spectrometry whose abundance significantly differs (FC ≥ 1.5, Significance ≥ 20) in the OM fraction of the *lolA1* and *lolB1* deleted strains compared to the WT clustered by predicted localization (SignalP server). SPI: signal peptide I (OMP or periplasmic), SPII: periplasmic-facing lipoproteins, SPII-LES: surface-exposed lipoproteins (**A**). Proteins with a SP whose abundance is affected in the OM fraction of the *lolA1* and *lolB1* mutants compared to the WT clustered by functional class (EggNOG) (**B**). Gliding and T9SS specific proteins whose abundance is altered in *lolA1* and/or *lolB1* OM compared to the WT (**C**).

In both mutants, the most affected known cellular functions were polysaccharide utilization (2 proteins increased (+) and 45 decreased (-) in *lolA1*, +6/-14 in *lolB1*) and cell wall/membrane/envelope biogenesis (+5/-21 in *lolA1*, +5/-12 in *lolB1*). In addition, several gliding/T9 secretion-specific proteins were affected too (+3/-9 in *lolA1*, +3/-6 in *lolB1*). Overall, 207 SPI/SPII proteins were impacted by the deletion of *lolA1* and 106 by the deletion of *lolB1* (**Figure 3B**). Among these, the abundance of 59 proteins was decreased in both mutants and among them several gliding/T9 secretion-related proteins (**Figure 3C**). GldK and GldJ, two lipoproteins essential for gliding motility and T9 secretion (26) were much less abundant in the OM of the *lolA1* and *lolB1* mutants than in the WT strain (GldK FC in *lolA1* = 0.48 and 0.08 in *lolB1*; GldJ FC in *lolA1* = 0.05 and 0.22 in *lolB1*). GldN, another critical component of the gliding and T9 machineries and the physical partner of GldK (27, 28) was less abundant as well (FC = 0.53 in *lolA1* and 0.33 in *lolB1*). In addition, one SprF-like protein (Fjoh_3951), implicated in T9 secretion (26), was also less abundant in the OM of both mutants (FC = 0.16 in *lolA1* and in *lolB1*). We also noticed a few proteins potentially involved in envelope biogenesis whose amount in the OM was altered in the two mutants, namely a peptidoglycan-binding LysM protein (FC = 3.12 in *lolA1* and 1.86 in *lolB1*), a BamB-like protein (FC = 0.44 in *lolA1* and 0.57 in *lolB1*), a polysaccharide export protein Wza (FC = 0.27 in *lolA1* and 0.23 in *lolB1*), and a peptidoglycan hydrolase (FC = 0.18 in *lolA1* and 0.28 in *lolB1*). Among the 148 proteins specifically affected by the deletion of *lolA1*, we found other components of the gliding/T9 secretion machineries (**Figure 3C**). Among them, SprT (FC = 0.41), SprF (FC = 0.37), and SprD (FC = 0.05) are all required for secretion of SprB to the surface of *F. johnsoniae* and, thus, for gliding motility (29, 30). Absence of *lolA1* resulted in the decrease of numerous OM proteins dedicated to polysaccharide utilization and inorganic ion transport (**Figure 3B**) of which most were SusC-like proteins or TonB-dependent receptors, *i.e.* proteins with a β-barrel domain.

Within the 47 proteins specifically affected by the deletion of *lolB1*, we also found gliding/T9 secretion-related proteins (**Figure 3C**). Interestingly, LolA1 amount was also increased in the *lolB1* mutant (FC = 2.04). In contrast with the data specific to *lolA1*, we did not observe an obvious pattern in the data specific to *lolB1*. The loss of *lolB1*, as for *lolA1*, decreased the number of PUL-encoded proteins in the OM but to a much lower extent (**Figure 3B**).

We did not observe a general mislocalization of OM lipoproteins upon deletion of *lolA1* or *lolB1*, as one could have expected. Nevertheless, the proteomics data confirm and provide an explanation for the loss of gliding motility and T9 secretion that we observed for both mutants. Although the deletion of *lolA1* or *lolB1* alters the OM protein composition in different ways, bacteria lacking either of these two proteins fail to properly localize a subset of proteins and lipoproteins, including core components of gliding motility/T9 secretion such as GldN, GldK and GldJ. While the deletion of *lolA1* was more consequential for the OM of *F. johnsoniae* than the deletion of *lolB1*, the most impacted cellular functions (polysaccharide utilization and cell wall/membrane/envelope biogenesis) were in common. Considering that the abundance of several OM lipoproteins was altered in both mutants, our data suggest that *lolA1* and *lolB1* participate in the same pathway, *i.e.* the transport of lipoproteins to the OM and in particular lipoproteins involved in gliding motility and T9 secretion. Yet, *lolA1* deletion clearly determined a bigger perturbation of the OM proteome.

### Deletion of *lolA1* and of *lolB1* affects growth and cell morphology

Deletion of *lolA1* and *lolB1* determines lack of gliding and T9 secretion as well as alterations in the OM proteome composition. We thus wondered whether lack of LolA1 and LolB1 would also affect bacterial fitness. To this aim we monitored the growth of these mutants in two media, casitone yeast extract (CYE), the rich medium normally used to grow *F. johnsoniae* and motility medium (MM) a poor medium that stimulates gliding motility (31).

As shown in **Figure 4A**, all mutants displayed a growth like the WT in CYE. In contrast, in MM, the deletion of *lolA1* severely impacted growth while deletion of *lolB1* did not (**Figure 4A**). In the absence of both LolA1 and LolB1 growth was comparable to that of the single *lolA1* mutant. Plasmid-borne complementation of *lolA1* and *lolA1lolB1* mutants with *lolA1* and *lolA1* and *lolB1* fully restored growth to the WT levels (**Figure 4A**). Deletion of any of the other LolA and LolB proteins did not affect growth in any condition tested (**Figure 4A**).

**Figure 4.**
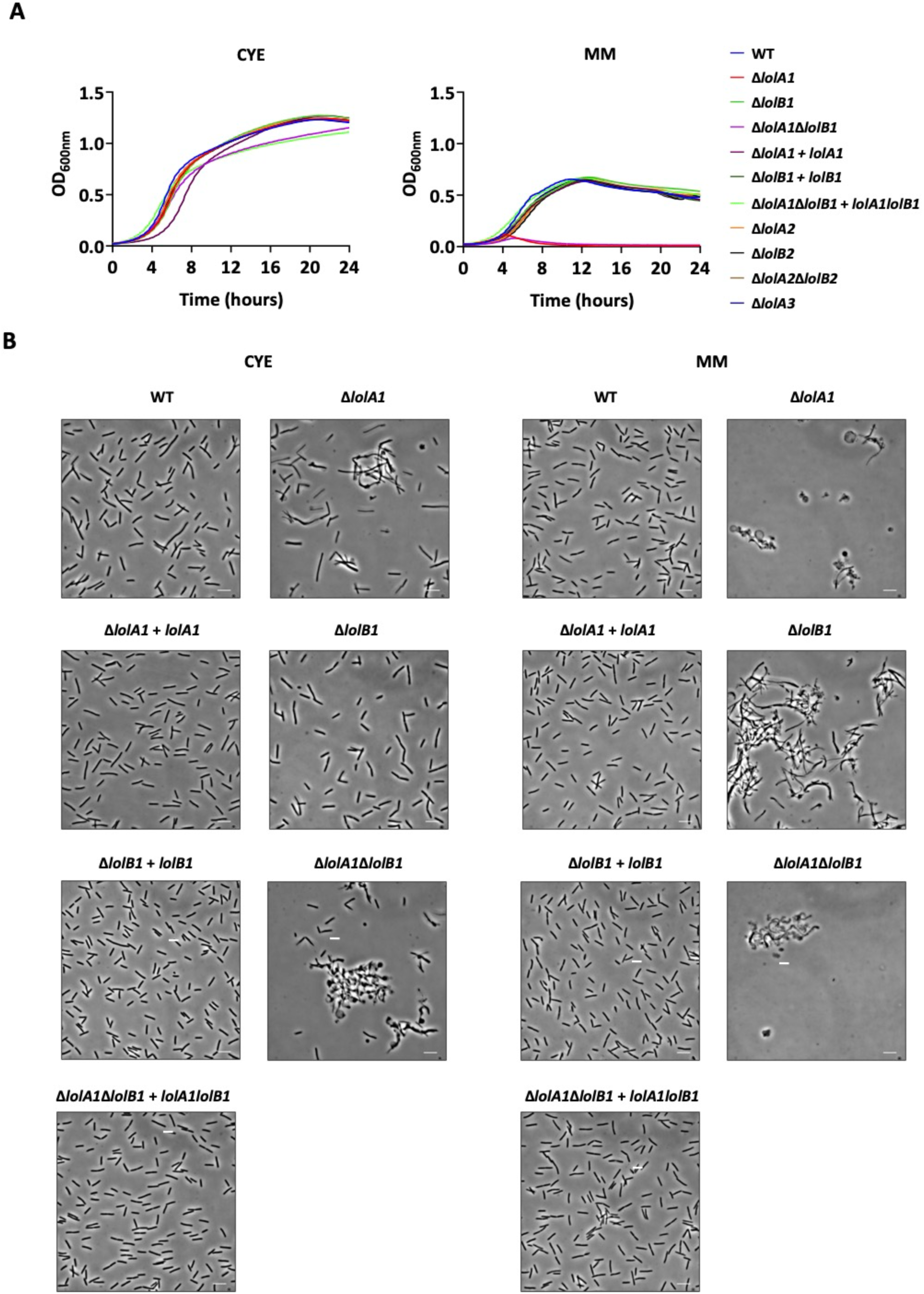
Deletion of *lolA1* and of *lolB1* affects growth and cell morphology. Growth curves of WT and mutant strains in CYE and MM liquid media (**A**). Bright field microscopy images of bacteria grown for 16 hours in CYE and MM media (bar = 5µm) (**B**).

Next, we assessed whether the mutations had any morphological effect on bacteria. We observed by optic and transmission electron microscopy cells grown for 16 hours in CYE medium and found that *lolA1* bacteria presented some aberrant shapes, some resembling the shape of a lollipop, while others being completely round (**Figure 4B** and **Supplementary Figure S3**). The same abnormal morphologies, along with more cell aggregation, were also observed in the *lolA1lolB1* double mutant, while *lolB1* mutants resembled the WT strain. The aberrant cell morphology phenotype of the *lolA1* and *lolA1lolB1* strains was enhanced when bacteria were grown in MM and, interestingly, the formation of abnormal cells was also observed for the *lolB1* mutant in this medium (**Figure 4B**). Deletion of any of the other LolA and LolB proteins did not affect cell morphology in either CYE or MM (**Supplementary Figure S4**). Complementation of *lolA1, lolB1* and *lolA1lolB1* with plasmid-borne *lolA1, lolB1*, and *lolA1* and *lolB1* fully restored the cell morphology to WT levels (**Figure 4B**).

Overall, while growth in MM medium was affected only by the deletion of *lolA1,* deletion of *lolA1* and of *lolB1* affected cell morphology, in particular in MM for this latter mutant. Co-deletion of *lolA1* and *lolB1* determined an enhancement in abnormal cell morphology in both CYE and MM **(Figure 4B** and **Supplementary Figure S3)**.

### Deletion of *lolA1* and to a minor extent of *lolB1* affects envelope integrity

The observed growth and morphological phenotypes of the *lolA1* mutant when grown in MM hinted to a possible membrane instability. A main difference in composition between these two media is the absence of magnesium in the form of MgSO_4_ in MM. It is known and has been largely reported that divalent cations and in particular magnesium stabilize LPS–LPS interactions (32). In bacteria with OM alterations due to protein and/or lipid shifts, magnesium has been shown to compensate for these phenotypes. We thus tested whether the growth defect and morphology changes observed for the *lolA1* and *lolB1* mutants could be rescued by magnesium supplementation in MM. As shown in **Figure 5**, addition of 8 mM MgSO_4_ (as in CYE) to MM restored growth (**Figure 5A**) of the *lolA1* and *lolA1lolB1* mutants, thus suggesting an alteration of the OM of these mutants. Concerning cell morphology, addition of magnesium alleviated abnormal cell morphologies of the mutants but did not totally restore them to WT levels (**Figure 5B**).

**Figure 5.**
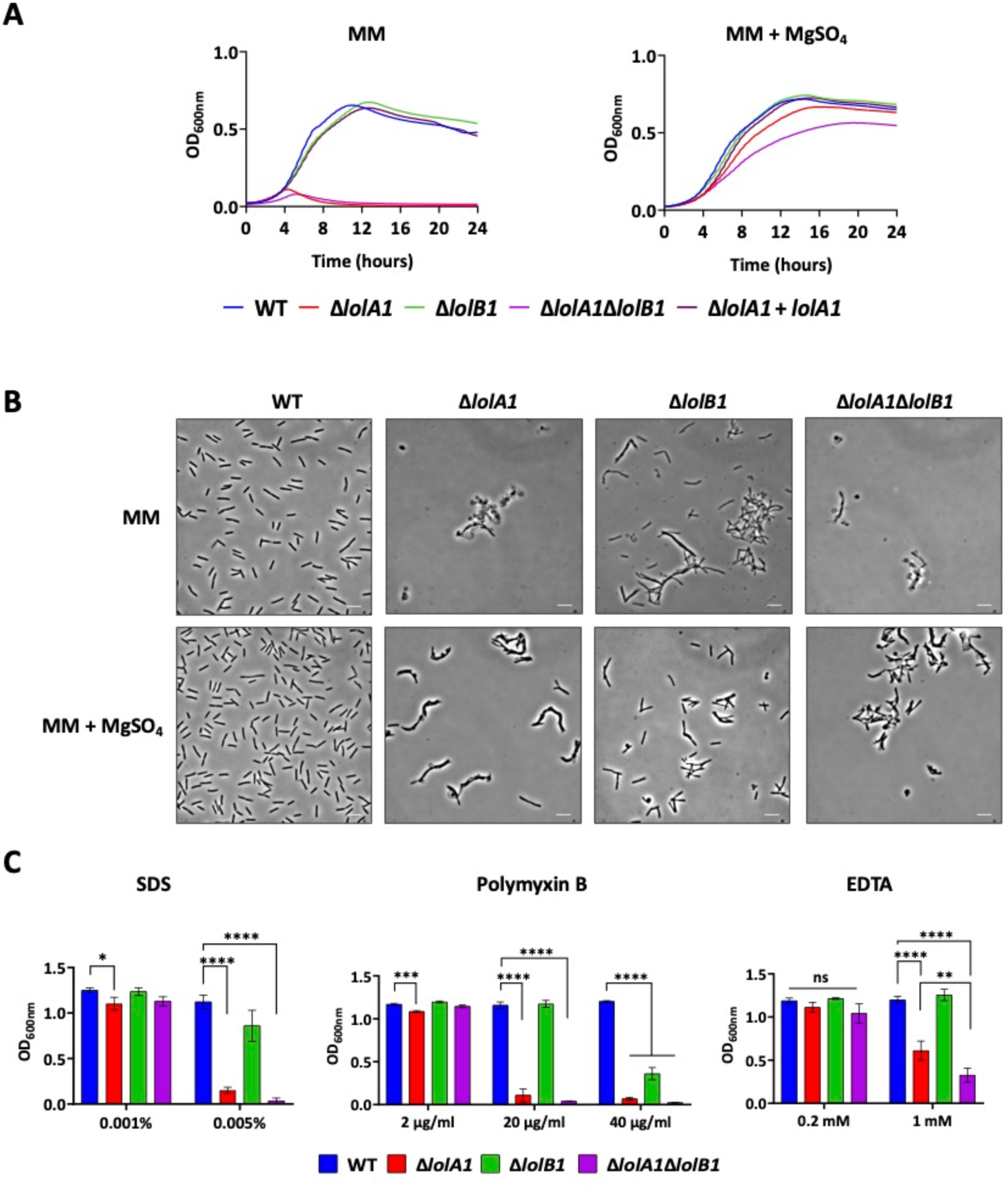
Impact of deletion of *lolA1* and of *lolB1* on OM integrity and stress sensitivity. Growth curves of WT and mutant strains in MM liquid medium without and with 8 mM MgSO_4_ (**A**). Bright-field microscopy images of bacteria grown in liquid MM medium without and with 8 mM MgSO_4_. (bar = 5µm) (**B**). Growth after 24 hours in CYE liquid medium in the presence of different OM perturbing agents (SDS, Polymyxin and EDTA) (**C**). Data is represented as means of 3 biological replicates ± standard deviation, and statistical analysis was a one-way ANOVA followed by Tukey’s multiple comparison test (SDS 0.001% F-value = 6.602; SDS 0.005% F-value = 440.9; Polymyxin B 2 µg/ml F-value = 43.21; Polymyxin B 20 µg/ml F-value = 540.5; Polymyxin B 40 µg/ml F-value = 669.9; EDTA 0.2 mM F-value = 4.37; EDTA 1 mM F-value = 99.03; DF = 3). Asterisks indicate the following: (*): p<0.0332, (**): p<0.0021, (***): p<0.0002, (****): p<0.0001.

We also tested whether *lolA1* and/or *lolB1* deletion might affect the sensitivity of the mutants to several OM perturbing agents, namely SDS, Polymyxin B and EDTA. We observed that while deletion of *lolA1* and/or *lolB1* did not affect growth at low SDS concentration, the *lolA1* mutant showed significant growth impairment at a higher concentration (**Figure 5C**). The *lolB1* strain also showed reduced growth even though to a lesser extent than the *lolA1* mutant (**Figure 5C**). The *lolA1lolB1* double mutant showed a phenotype like the *lolA1* mutant. Polymyxin B affected the growth of the three mutants in a similar concentration-dependent fashion (**Figure 5C**). In contrast, EDTA affected the growth of the *lolA1* and *lolA1lolB1* mutants, but not of the *lolB1* mutant at the tested concentrations (**Figure 5C**). Interestingly, at an EDTA concentration of 1 mM, the *lolA1lolB1* double mutant showed an increased sensitivity compared to that of the *lolA1* mutant, meaning that the deletion of *lolB1* in a *lolA1* mutant background increases the sensitivity of bacteria to this compound.

Taken together, these results suggest that deletion of *lolA1* affects OM stability, as the *lolA1* and *lolA1lolB1* mutants showed acute sensitivity to all tested stresses. On the other hand, *lolB1* single deletion seems to have a lower impact on OM stability, as this mutant was less sensitive than the *lolA1* one to SDS and Polymyxin B and resisted to EDTA. The increased sensitivity of the *lolA1lolB1* double mutant to EDTA compared to the *lolA1* single mutant support our observations of cell morphology (**Figure 4B**) as well as the idea that the deletion of both *lolA1* and *lolB1* is more detrimental to *F. johnsoniae* than their single deletion.

### The protruding loop and the C-terminal region of LolB1 are crucial for its function

In *E. coli*, the hydrophobic protruding loop between β-strands 3 and 4 is critical for lipoprotein insertion (33). A highly conserved residue, Leu68, found at the tip of the loop has been shown to be crucial for LolB activity. When this leucine is replaced with a polar residue or deleted, LolB can no longer efficiently insert lipoproteins into the OM (13, 33). A loop is also present in LolB1 with a leucine (L74) at its tip (**Figure 6A**). To see whether the loop and this residue are crucial for LoLB1 function, we tested whether expression of LolB1 where Leu74 is replaced by the polar glutamate (L74E) or where the loop is deleted (residues 73-76) could complement the gliding and T9SS phenotypes of the *lolB1* mutant. As shown in **Figure 6B**, the LolB1_L74E_ variant only partially restored gliding motility while amylase secretion was like WT levels. In contrast, deletion of the loop completely abolished gliding and secretion.

**Figure 6.**
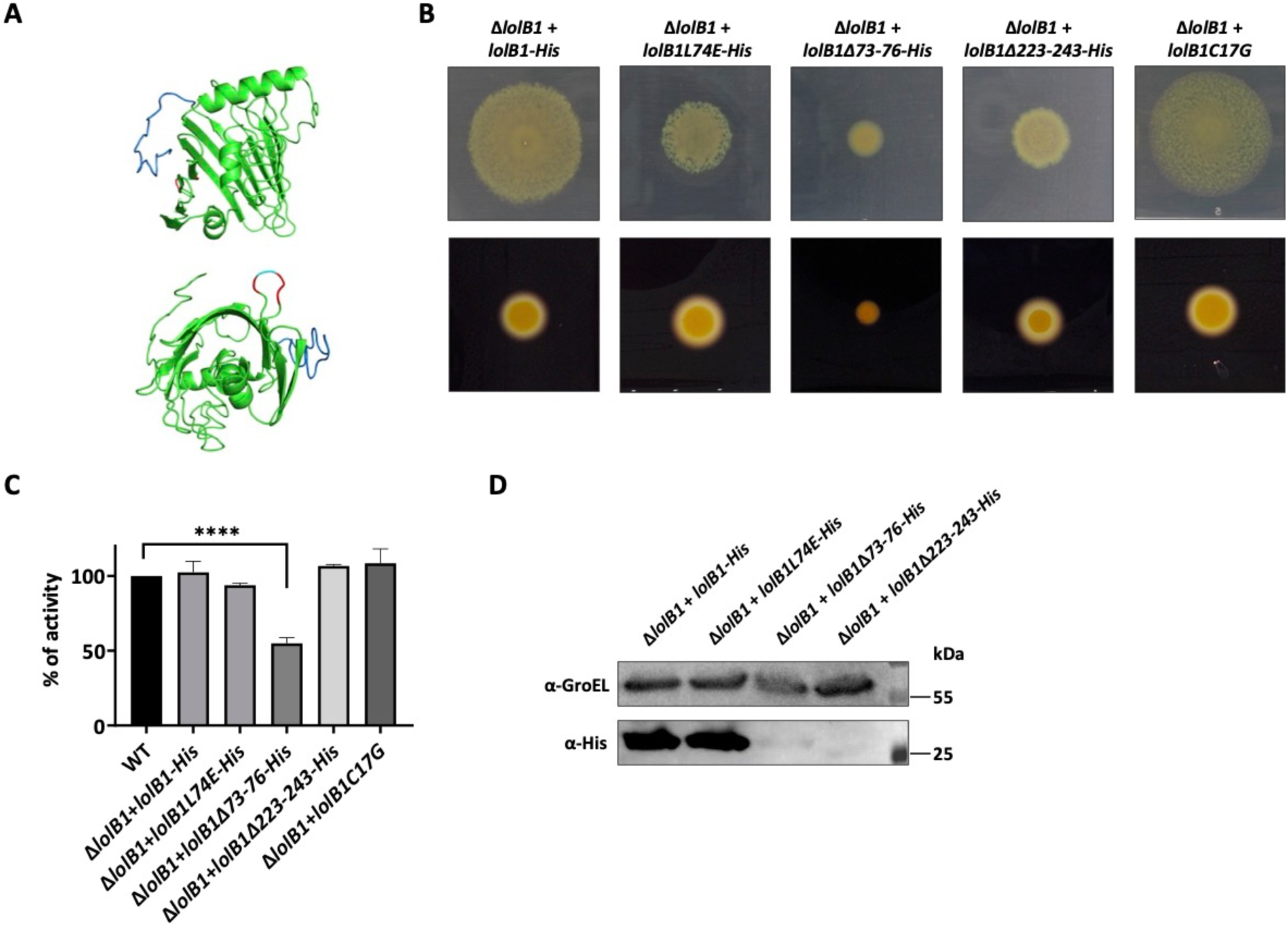
The protruding loop and the C-terminal domain of LolB1 are crucial for its function. I-TASSER model of LolB1 with the protruding loop (in red) with the conserved L74 (in cyan) and the C-terminal domain (in blue) (**A**). Gliding on MM agar plates after 48h and starch degradation on LB starch plates after 24h of *lolB1* mutants expressing: WT LolB1, LolB1 _L74E_, LolB1_Δ73-76_, LolB1_Δ223-243_ and LolB1_C17G_ (**B**). Amylase activity of the supernatant of WT and *lolB1* mutants expressing: WT LolB1, LolB1 _L74E_, LolB1_Δ73-76_, LolB1_Δ223-243_ and LolB1_C17G_ (**C**). Data is represented as means of 3 biological replicates ± standard deviation, and statistical analysis was a one-way ANOVA followed by Dunnett’s multiple comparison test (F-value = 47.47; DF = 22). Asterisks indicate the following: (****): p<0.0001. Detection by Western blot of C-terminally His-tagged: WT LolB1, LolB1 _L74E_, LolB1_Δ73-76_ and LolB1_Δ223-243._ GroEL protein levels are shown and used as loading control (**D**).

These data suggest that the loop and the conserved Leu are important for LolB1 function, as in *E. coli* LolB. Partial complementation of the LolB1^L74E^ protein could not be ascribed to a different protein expression level of this mutation since protein amount of WT LolB1 and LolB1_L74E_ variants was comparable (**Figure 6C**). In contrast, deletion of residues 73-76 of LolB1 (LolB1_Δ73-76_) resulted in significant reduction of protein amount, thus suggesting that this mutation affects protein stability or induces degradation of this non-functional LolB variant (**Figure 6C**).

A main difference between *E. coli* LolB and LolB1 is the presence of a C-terminal domain which could not be folded by modelling softwares (**Figure 6A**). We wondered whether this domain could be important for LolB1 function and thus deleted it (*lolB1Δ223-243*). This mutant LolB1 could only partially complement gliding while amylase secretion was like WT levels (**Figure 6B**). As for LolB1_Δ73-76_, deletion of the C-terminal domain determined a significant reduction in the protein levels, thus suggesting that this domain of LolB1 is crucial for protein function and/or stability.

In *E. coli*, LolB does not require its lipid anchor to correctly insert lipoproteins in the OM (34). To see if this was also the case for LolB1, we generated a periplasmic soluble form of LolB1, by introducing a C17G mutation in the signal peptide of LolB1 (LolB1_C17G_), thus replacing the lipidated cysteine and generating an SPI. *In trans* expression of LolB1 _C17G_ fully complemented *lolB1* deletion in *F. johnsoniae* as shown by the recovered gliding motility and amylase secretion (**Figure 6B**). As for *E. coli* LolB, lipidation does not seem to be crucial for *F. johnsoniae* LolB1 function.

Overall, these data support the evidence that LolB1 shares common features with *E. coli* LolB, as the loop, while having some compositional differences, and the C-terminal domain, both crucial for its function.

### *E. coli* and *F. johnsoniae* LolA and LolB are not interchangeable

*In silico* structure comparison and protein characterization show high structural similarity between LolA1 and LolB1 and *E. coli* LolA and LolB (**Figure 1**) while highlighting some differences regarding hydrophobicity and charge distribution (**Supplementary Figures S1** and **S2 and Supplementary Table S1**). We thus wondered whether LolA and LolB from *E. coli* might complement the deletion of *lolA1* and *lolB1* in *F. johnsoniae*. Surprisingly, in *trans* expression of LolA*_Ec_* and LolB*_Ec_* did not restore gliding nor T9 secretion of the *lolA1* and *lolB1* mutants (**Figure 7A** and **7B**). Similarly, co-expression of LolA*_Ec_* and LolB*_Ec_* did not restore gliding nor secretion of the *lolA1lolB1* double mutant (**Figure 7A** and **7B**). However, expression of LolA*_Ec_* in *lolA1* bacteria partially complemented growth and cell shape in MM medium (**Figure 7C** and **7D**). The same could be observed for the double *lolA1lolB1* mutant expressing LolA*_Ec_* and LolB*_Ec_*. Cell shape was also enhanced in *lolB1* bacteria expressing LolB*_Ec_* (**Figure 7C** and **7D**). Next, we tested whether *F. johnsoniae* LolA1 and LolB1 could complement the deletion of *lolA* and *lolB* in *E. coli*. Since *lolA* and *lolB* are essential genes in *E. coli,* we first expressed the *F. johnsoniae* proteins LolA1 and LolB1 individually or together (LolA1-LolB1) in the *E. coli* MG1655 WT strain and then attempted to delete *lolA* or *lolB*. While deletion of *lolA* and *lolB* could be achieved when LolA*_Ec_* and LolB*_Ec_* were expressed *in trans* (**Supplementary Figure S5**), we could not obtain any mutant when the *F. johnsoniae* proteins were expressed. Similar results were obtained when LolA2 and/or LolB2 proteins were expressed in *E. coli*. In conclusion, despite structural and physico-chemical similarities, *E. coli* and *F. johnsoniae* LolA and LolB proteins are not interchangeable. Interestingly, the evidence that *E. coli* LolA and LolB do not restore gliding and T9 secretion in *lolA1* and *lolB1* bacteria might reflect differences among the clients of these proteins, namely OM lipoproteins involved in gliding and T9 secretion which are unique to Bacteroidetes and absent in *E. coli*.

**Figure 7.**
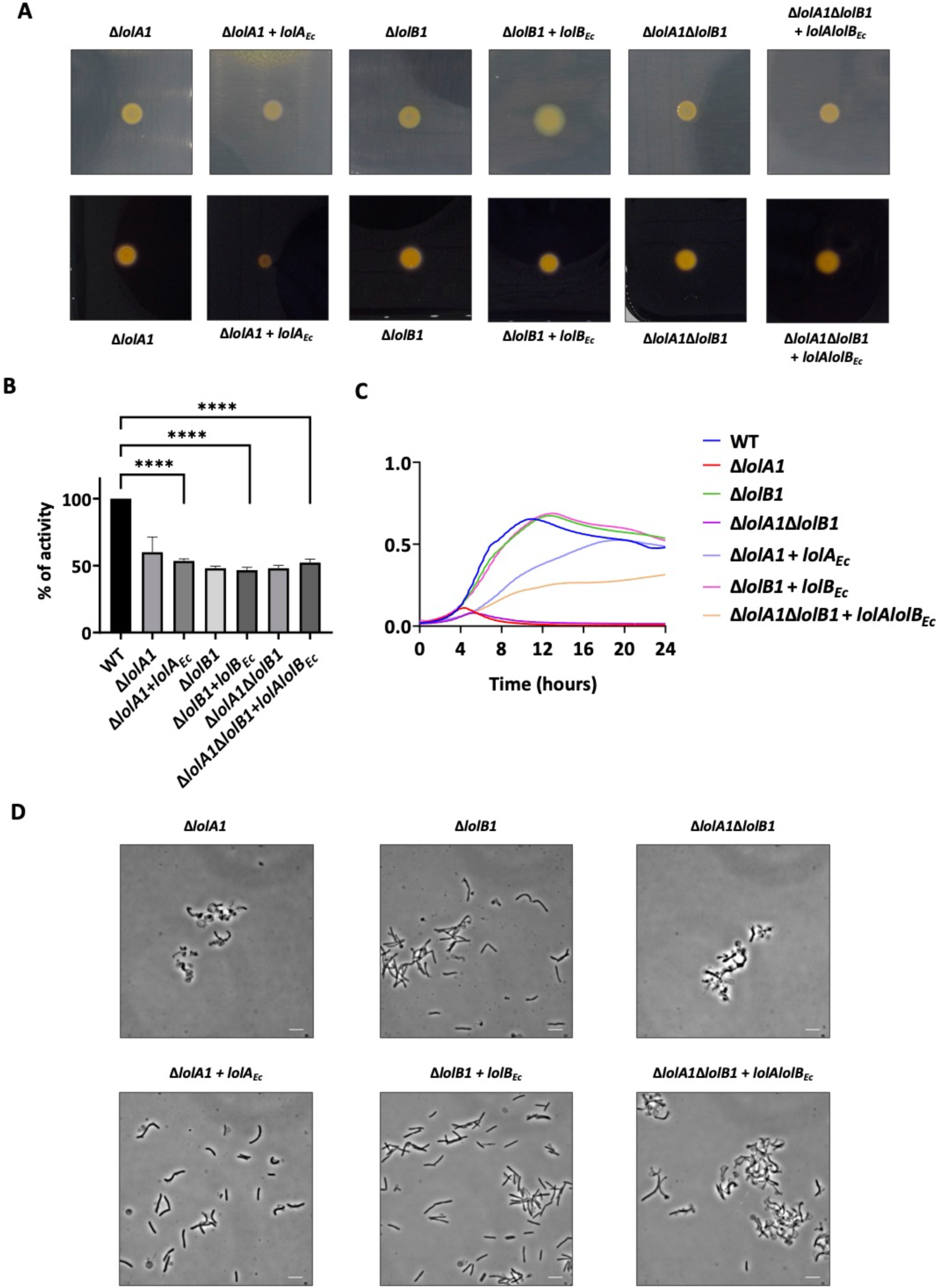
*E. coli* LolA and LolB partially complement *lolA1* and *lolB1* deletion in *F. johnsoniae*. Gliding motility on MM agar plates after 48h and starch degradation on LB starch plates after 24h (**A**). Amylase activity of cell culture supernatants. Data is represented as means of 3 biological replicates ± standard deviation, and statistical analysis was a one-way ANOVA followed by Dunnett’s multiple comparison test (F-value = 47.47; DF = 22). Asterisks indicate the following: (****): p<0.0001 (**B**). Growth curves in MM liquid medium (**C**). Bright field microscopy images of bacteria grown in liquid MM medium for 16 hours (bar = 5 µm) (**D**).

### LolA and LolB are conserved among Bacteroidetes

We next explored if the *F. johnsoniae* LolA and LolB proteins were conserved in bacteria of the phylum Bacteroidetes. To this aim, we performed an *in silico* sequence similarity search for homologs of the five *F. johnsoniae* LolA and LolB proteins in several Bacteroidetes species. Interestingly, this analysis identified LolA and LolB homologs in all the analyzed species (**Figure 8** and **Supplementary Table S4**). In addition, while we found LolA1 and LolB1 homologs in all species, LolA2, LolB2 and LolA3 homologs seem to be more restricted. Among the species carrying only LolA1 and LolB1 homologs, there is *C. canimorsus* which belongs to the family Flavobacteriaceae and is a normal oral commensal of dogs and cats which can cause severe infections in humans upon contact with these animals (35). In *C. canimorsus*, LolA and LolB homologs: Ccan_16490 (LolA) and Ccan_17050 (LolB) are essential since attempts to delete their encoding genes failed. Regarding *lolA2* and *lolB2*, both genes were always found to be encoded in *loci* dedicated to synthesis and transport of a flexirubin-like pigments (19), suggesting that LolA2 and LolB2 are specific to these processes.

**Figure 8.**
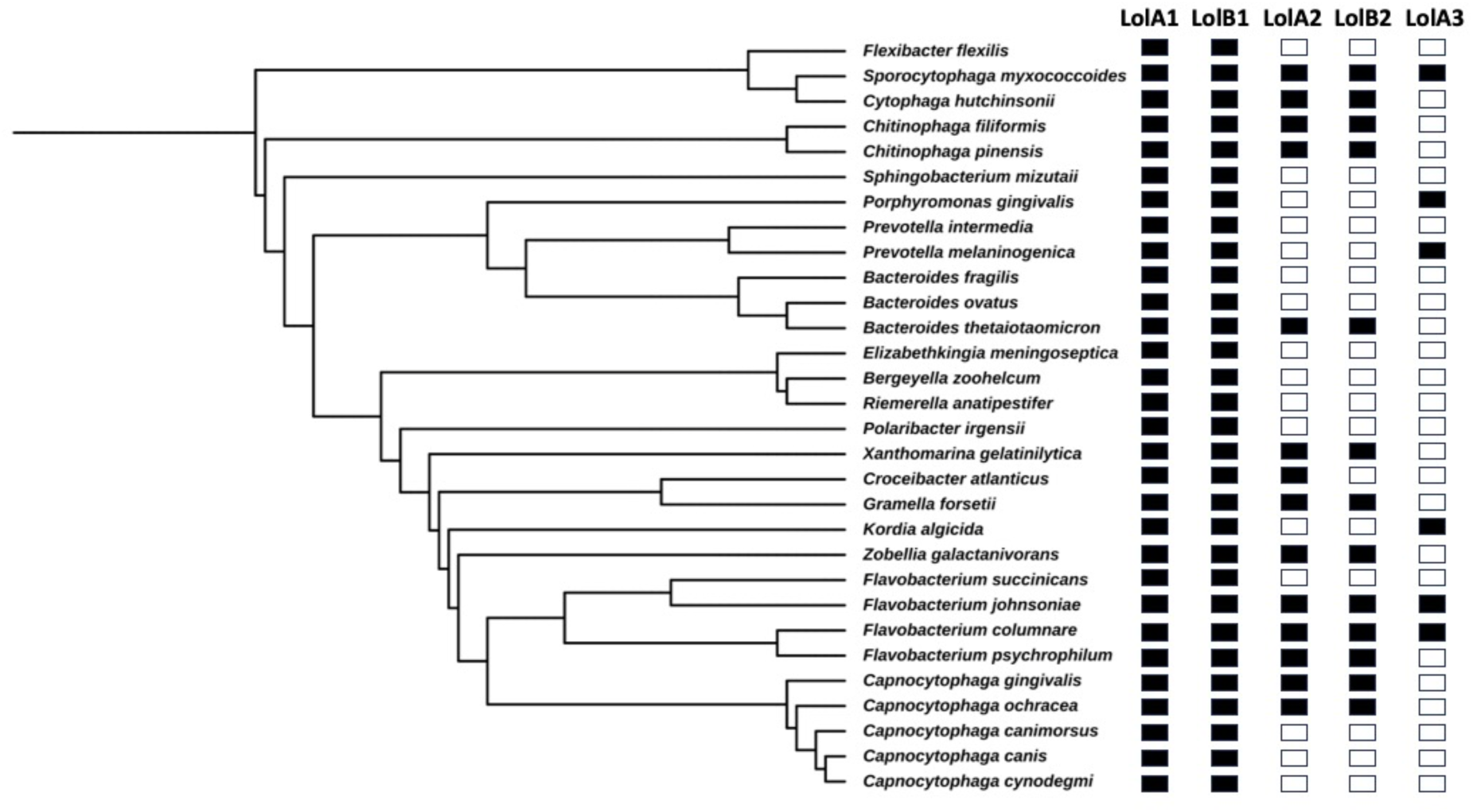
Conservation of LolA and LolB in Bacteroidetes. Homologs of *F. johnsoniae* LolA and LolB proteins in several Bacteroidetes species identified by DELTA Blast (36). Phylogenetic tree based on NCBI taxonomy generated via phyloT (37).

To see whether the *F. johnsoniae* LolA1 and LolB1 and *C. canimorsus* LolA and LolB homologs share the same function in these bacteria we first tested whether expression of *C. canimorsus* LolA (Ccan_16490) and LolB (Ccan_17050) could complement the deletion of *lolA1* and *lolB1* in *F. johnsoniae* respectively. Complementation with *C. canimorsus* LolA or LolB fully restored the ability to glide and secrete via the T9SS of the *F. johnsoniae lolA1* and *lolB1* mutants (**Figure 9A** and **9B**) as well as the growth and cell morphology phenotypes in MM (**Figure 9C** and **9D**).

**Figure 9.**
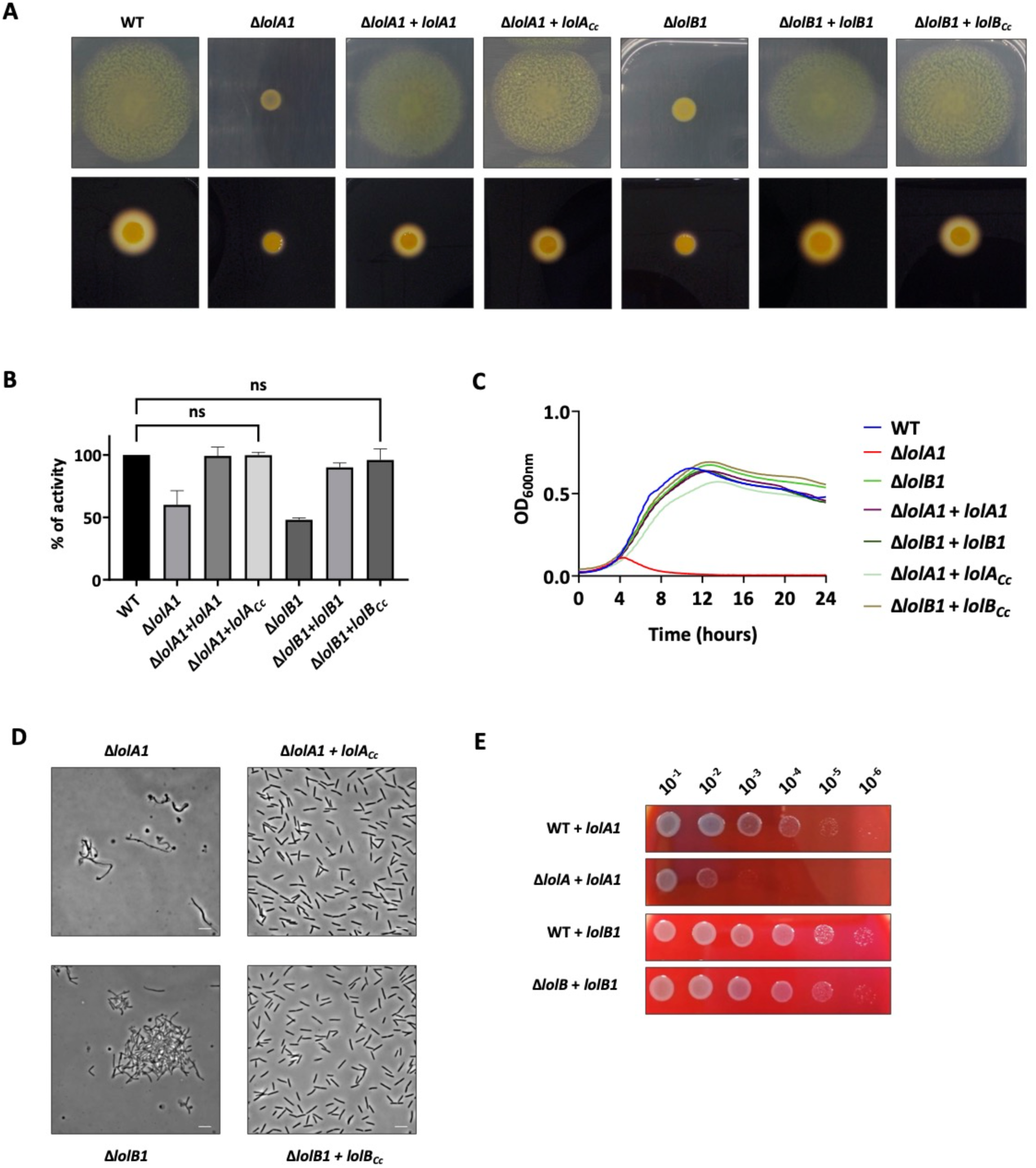
Heterologous complementation of LolA and LolB mutants of *F. johnsoniae* and *C. canimorsus* with LolA1 and LolB1 homologs. Gliding motility on MM agar plates after 48h (top), starch degradation by T9-secreted amylases on LB starch plates after 24h (down) (**A**) and amylase activity of cell culture supernatants (**B**) of Δ*lolA1* and Δ*lolB1 F. johnsoniae* strains complemented with *C. canimorsus* LolA (LolA*_Cc_*) and LolB (LolB*_Cc_*). Data is represented as means of 3 biological replicates ± standard deviation, and statistical analysis was a one-way ANOVA followed by Dunnett’s multiple comparison test. Growth in MM liquid medium (**C**) and bright-field microscopy images of bacteria grown in liquid MM medium for 16 hours (**D**) of Δ*lolA1* and Δ*lolB1 F. johnsoniae* strains complemented with *C. canimorsus* LolA (LolA*_Cc_*) and LolB (LolB*_C_c*) (bar = 5 µm). Growth of serial dilution spots of *C. canimorsus lolA* and *lolB* mutants expressing *F. johnsoniae* LolA1 (LolA1) or LolB1 (LolB1) *in trans* on SB plates (**E**).

We next tested whether expression of *F. johnsoniae* LolA1 and LolB1 could bypass the lethality of the deletion of *lolA* and *lolB* of *C. canimorsus.* To test complementation with LolA1, we performed a plasmid exchange in the *C. canimorsus lolA (Ccan_16490)* mutant strain expressing plasmid-borne *C. canimorsus lolA* with the vector encoding *lolA1* (see methods for additional details). To test LolB1 complementation, we first expressed LolB1 *in trans* in the *C. canimorsus* WT strain and then deleted *lolB* (*Ccan_17050*). Interestingly, both *C. canimorsus lolA* and *lolB* deletion mutants were viable when LolA1 or LolB1 were expressed thus confirming that the *F. johnsoniae* and *C. canimorsus* proteins share similar functions (**Figure 9E**). However, while complementation of *lolB* deletion in *C. canimorsus* with *lolB1* did not result in any growth defect on rich medium, expression of *lolA1* only partially complemented *lolA* deletion (**Figure 9E**). Overall, these results suggest that homologs of *F. johnsoniae* LolA1 and LolB1 share the same function in *C. canimorsus* and that the same could be true in Bacteroidetes in general.

### Simultaneous deletion of all *F. johnsoniae* LolA and LolB homologs is not lethal and does not affect surface lipoprotein localization

Next, to see whether *F. johnsoniae* could tolerate the lack of all its LolA and LolB homologs, we tried to generate a *lolA1 lolA2 lolA3 lolB1 lolB2* quintuple mutant by sequentially deleting all genes. Surprisingly, we succeeded in obtaining this strain, indicating that the bacterium can perform its vital functions even in the absence of any LolA and LolB.

Finally, we tested whether surface-exposed lipoproteins are correctly localized in the absence of all LolA and LolB homologs in *F. johnsoniae*. To this end, we monitored surface lipoprotein exposure using a reporter lipoprotein: the sialidase (SiaC) from *C. canimorsus*. SiaC is an OM lipoprotein facing the periplasm where it is responsible for the cleavage of sialic acid from eukaryotic glycoproteins (38). Addition of the lipoprotein export signal (LES) to SiaC (LES-SiaC) was shown by our group to be sufficient to localize it at the surface of *C. canimorsus* (16, 17). We thus expressed SiaC and LES-SiaC in the *F. johnsoniae* WT and in the *lolA1 lolA2 lolA3 lolB1 lolB2* quintuple mutant strain and verified their localization by immunofluorescence microscopy using anti-SiaC antibodies and secondary fluorescently labeled antibodies (**Figure 10**). As expected, SiaC WT was not exposed at the bacterial surface of both strains while we found LES-SiaC surface exposed in both WT and LolAB lacking strains (**Figure 10**) thus suggesting that surface-exposed lipoproteins correctly localize at the cell surface in the absence of any LolA and LolB proteins and thus that the pathway responsible for surface-lipoproteins localization is independent of LolA and LolB homologs of *F. johnsoniae*.

**Figure 10.**
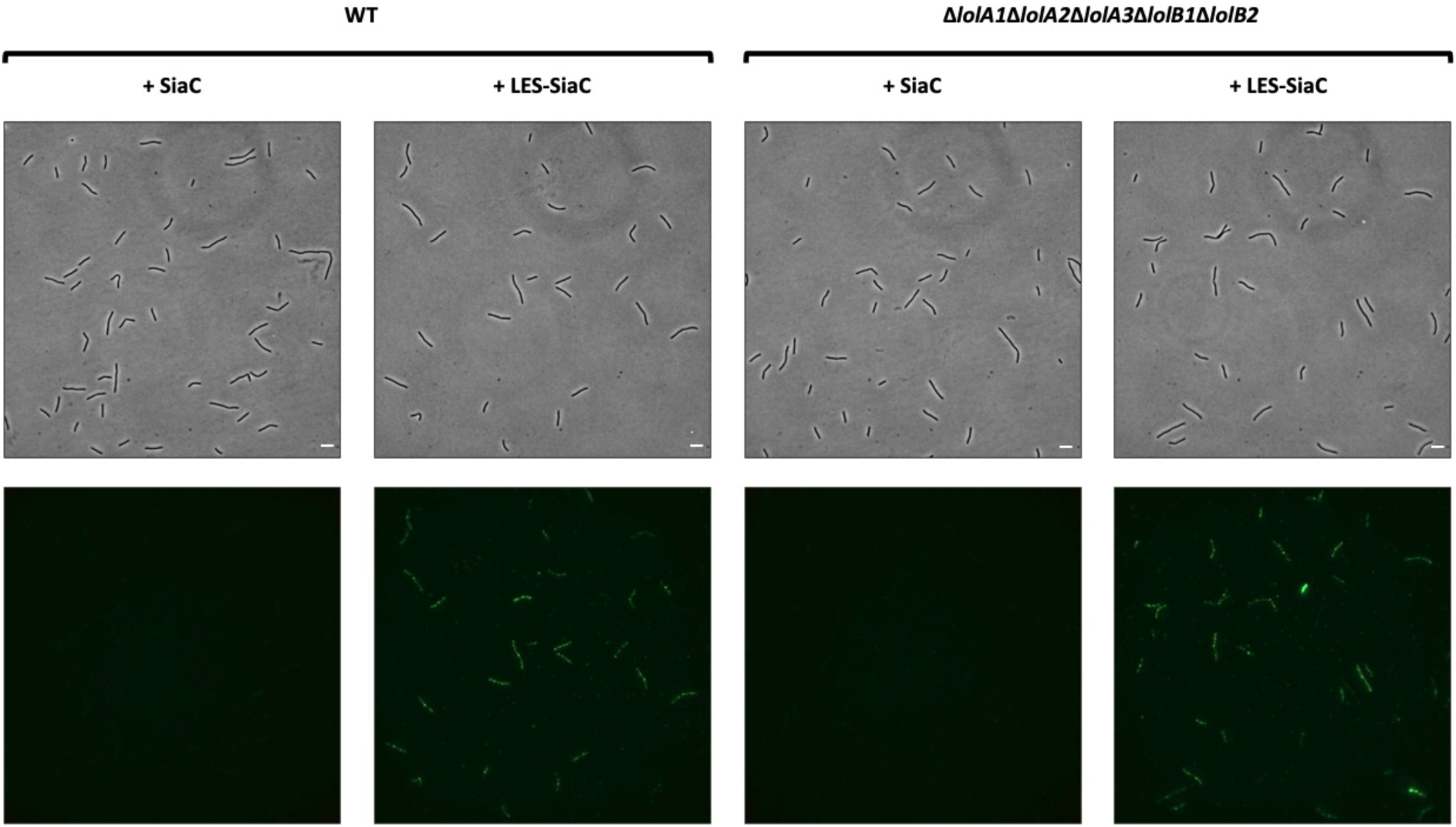
Deletion of all LolA and LolB homologs of *F. johnsoniae* does not affect surface lipoproteins localization. Immunofluorescence microscopy images of WT and *lolA1 lolA2 lolA3 lolB1 lolB2* mutant bacteria expressing sialidase (SiaC) or LES-sialidase (LES-SiaC) labeled with anti-SiaC serum (scale bar = 5 µm).

## Discussion

In conclusion, we show for the first time that Bacteroidetes possess LolB homologs, indicating that, in contrast to what was thought, the Lol pathway is conserved. In addition, we found that several LolA and LolB can coexist in some Bacteroidetes species such as *F. johnsoniae*, which encodes 3 LolA and 2 LolB homologs. We show that deletion of *lolA1* and/or *lolB1* results in the loss of gliding motility and T9 secretion. Proteomic data show that these two phenotypes can be explained by the significant reduction of key proteins and lipoproteins involved in gliding/T9 secretion in the OM of both *lolA1* and *lolB1* mutants. Indeed, absence of OM lipoproteins GldJ and/or GldK is sufficient to completely abolish gliding and T9 secretion (8, 27). We believe that these lipoproteins might be mislocalized because they require LolA1 and then LolB1 to reach the OM. Two other proteins essential for gliding motility, SprB and SprD, were decreased in *lolA1* and *lolB1*, though the significance did not meet our statistical parameters for both mutants **(Table 1** and **Supplementary Table S3**). Interestingly, the OM of the *lolA1* mutant was enriched in GldH (FC = 2.31), a lipoprotein necessary for the stability of GldJ and/or GldK (8, 23). This might be an attempt by *F. johnsoniae* to restore gliding motility and T9 secretion by stabilizing GldJ and GldK within the machinery. In *X. campestris* and *P. aeruginosa*, it is known that *lolA* deletion and depletion, respectively, impact cell envelope stability, with an increased sensitivity of the mutants to several OM stresses (39, 40). In *E. coli*, removal of *lolA* or *lolB* determines mislocalization of Lpp and RcsF that leads to cell death (41). Here, we show that deletion of *lolA1* and, to a lesser extent, *lolB1* destabilizes the OM of *F. johnsoniae*. Hence, we observe substantial cell shape abnormalities for both mutants in poor medium, and even in rich medium for the *lolA1* single and *lolA1lolB1* double mutants. We attempted to rationalize the weaker fitness of bacteria lacking LolA1 with our proteomic data. In total, the amount of four essential proteins of *F. johnsoniae* (Fjoh_0540, Fjoh_3469, Fjoh_5008, Fjoh_0105) was reduced in the OM of the *lolA1* mutant while not in the *lolB1* mutant (**Supplementary Table S2**). Besides, one of them is a lipoprotein homologous to *E. coli* BamD (Fjoh_3469, FC = 0.45), which is required for the activity of the BAM complex (42). Malfunction of the BAM complex would be a sensible explanation for the great decrease in β-barrel proteins that we observed in the *lolA1* OM. Interestingly, 13 SusC-like proteins (out of the 21 whose amount was reduced) were decreased with their SusD-like partner upon deletion of *lolA1*. Such a diminution of β-barrel proteins is a modification of the OM proteome that might destabilize the OM, to the point of making it more permeable to environmental stresses (**Figure 5C**). A possible explanation for the difference of fitness between *lolA1* and *lolB1* mutants could be that the IM essential LolCDE complex (41) becomes stalled in absence of LolA1 to pick up lipoproteins, which would obstruct or at least hinder the lipoprotein transport pathway. On the other hand, deletion of *lolB1* likely has no effect over the functioning of LolCDE, given their respective localization in the outer and inner membranes. Additionally, the evidence that, upon TEM samples observation, *lolA1* and *lolA1lolB1* bacteria with a detaching OM were often found in pairs (**Supplementary Figure S6**) and that two peptidoglycan degrading proteins were less abundant in the OM of *lolA1* (Fjoh_1328, FC = 0.31; Fjoh_1913, FC = 0.18) might hint at cell division defects. More data is needed to confirm this phenotype. The growth complementation and the partial restoration of cell shape aberrations in MM medium by MgSO_4_, a known stabilizer of LPS-LPS interactions, further support the idea that deletion of *lolA1* or *lolB1* disrupts OM stability.

In this work, we explored the structure of LolB1, based on our knowledge of *E. coli* LolB. The importance of the conserved protruding loop and the leucine 74 residue for LolB1 function (**Figure 6B**) strongly suggests that *E. coli* LolB and *F. johnsoniae* LolB1 work in a similar fashion. This hypothesis is supported by the fact that *E. coli* LolB and LolA partially complement the *lolB1* and *lolA1* mutants of *F. johnsoniae*, suggesting that LolB and LolA can transport a subset of *F. johnsoniae* lipoproteins which does not include gliding/T9 secretion lipoproteins. The partial complementation could also be due to differences in the structures and physico-chemical properties between *F. johnsoniae* LolA1 and LolB1 and *E. coli* LolA and LolB (**Figure 1** and **Supplementary Table S1, Figure S1** and **S2**). It could be interesting to compare the OM proteomes of *F. johnsoniae lolA1* + *lolA_Ec_* and *lolB1* + *lolB_Ec_* with those of *lolA1* and *lolB1* strains in order to identify the subset of lipoproteins which can be transported by LolA*_Ec_* and LolB*_Ec_*. Our data also show that proper activity of LolB1 necessitates a disordered C-terminal domain, which appears to be a unique feature of Bacteroidetes LolB1 proteins and that is absent from *E. coli* LolB.

Lastly, we found that LolA2 and LolB2 are encoded in the locus of flexirubin synthesis and transport of *F. johnsoniae* (19), and that these two proteins are conserved in Bacteroidetes which synthesize flexirubin or flexirubin-like pigments (**Figure 8**). Considering the ability of LolA and LolB-like proteins to bind hydrophobic moieties, this might suggest that LolA2 and LolB2 participate in the transport of either flexirubin/flexirubin precursors, or lipoproteins encoded in the locus. To our knowledge, this would be the first example of such a specialization of LolA and LolB-like proteins to a specific pathway and we aim to rule this out in the next future. Concerning LolA3, we could not observe any phenotype upon its deletion but its poor conservation and low prevalence in Bacteroidetes might suggest a specific role of this protein restricted to some species. Whether this protein can interact with LolB1 and/or LolB2 and participate to different pathways requires further investigations. As previously shown in *E. coli*, we were able to generate a viable mutant strain devoid of any LolA or LolB protein. Essential OM lipoproteins like those part of the BAM and Lpt complexes are likely correctly localized in *F. johnsoniae* despite the absence of LolA and LoB homologs, strongly suggesting that an alternative LolA-LolB independent lipoprotein sorting pathway exists in Bacteroidetes as well (41). In addition, we show that lipoproteins are still localized at the bacterial surface in the absence of LolA and LolB proteins thus suggesting that this specific class of lipoproteins does not require these elements of the Lol pathway for its localization. Overall, this study unveils the hidden diversity of LolA and LolB proteins which likely share a common function, *i.e.* the transport of lipidated cargos across the periplasm, but specialized over time to suit different pathways and organisms. Gene duplication events may have allowed some Bacteroidetes species like *F. johnsoniae* to repurpose LolA and LolB proteins to the sole transport of lipoproteins involved in gliding motility and T9 secretion or to the synthesis/transport of outer membrane lipids such as flexirubin, for example. This diversification of lipoprotein transport routes fits with the extensive use of OM lipoproteins by Bacteroidetes. We believe that these findings will contribute to a better understanding of lipoprotein trafficking in Gram-negative bacteria, a field of great importance that remains understudied outside of Proteobacteria.

## Methods

### Bacterial strains and growth conditions

Bacterial strains used in this work are listed in **Table S5**. Unless otherwise stated, *F. johnsoniae ATCC 17061* UW101 strains were routinely grown in Casitone Yeast Extract (CYE) medium (10 g/l casitone, 5 g/l yeast extract, 8 mM MgSO_4_ and 10 mM Tris-HCl pH 7.5) or Motility Medium (MM) (3.3 g/l casitone, 1.7 g/l yeast extract, 3.3 mM Tris-HCl pH 7.5) at 30°C. When required, the following concentrations of antibiotics were used: Gentamicin (Gm) at 20 μg/ml, Tetracycline (Tet) at 20 μg/ml, and Erythromycin (Em) at 100 μg/ml. *E. coli* strains were grown in lysogeny broth (LB) at 37°C and the following concentrations of antibiotics were used accordingly: Ampicillin (Amp) at 100 μg/ml, Kanamycin (Kan) at 50 μg/ml, Chloramphenicol (Cm) at 30 μg/ml and Tetracycline (Tet) at 12.5 μg/ml. *Capnocytophaga canimorsus* 5 strains were routinely grown at 37°C in the presence of 5% CO_2_ on heart infusion agar (HIA) supplemented with 5% sheep blood (SB) plates with, when required, the following concentrations of antibiotics: Gentamicin (Gm) at 20 μg/ml, Cefoxitin (Cfx) at 10 μg/ml, Erythromycin (Em) at 10 μg/ml and Tetracycline (Tet) at 10 μg/ml.

### *In silico* search for LolA and LolB homologs

To search for LolA and LolB remote homologs, whole proteome structure predictions were realized for *Flavobacterium johnsoniae* UW101 strain using the HHSearch suite (43) with default parameters. The structures of *E. coli* LolA and LolB were predicted using the same program and parameters. An in-house Perl script was used to compare the hit list for each of the *F. johnsoniae* proteins with the hit list for *E. coli* LolA and LolB, with cut-off value of 10^-3^. Proteins with identical templates were selected as potential candidates.

### *In silico* modelling and characterization of LolA and LolB homologs

From the sequences of identified full-length *F. johnsoniae* LolA1 (Fjoh_2111), LolA2 (Fjoh_1085), LolB1 (Fjoh_1066), and LolB2 (Fjoh_1084), proteins were modeled using the Iterative Threading ASSEmbly Refinement (I-TASSER) protein structure predictor without specifying secondary structure preferences and additional distance restraints or templates (44); models displaying the highest probability score were selected. Giving the poor quality of the LolA3 (Fjoh_0605) I-TASSER model, the best ranking AlphaFold 3 model was chosen instead for representing LolA3 structure (45). The presence of signal peptides was determined using the SignalP 6.0 server (46). As such, the N-terminal cleaved peptides were not considered. Electrostatic potentials and hydrophobicity scores were 3D mapped on the refined crystallographic structures of *E. coli* LolA (PDB entry: 1UA8) and LolB (PDB entry: 1IWL), as well as on the selected I-TASSER and AlphaFold 3 models of LolA and LolB variants from *F. johnsoniae*. For each protein, the corresponding PDB file was prepared for pKa and continuum solvation calculations with PROPKA 3.2 and PDB2PQR software in the AMBER force field, respectively. Equations of continuum electrostatics were solved using the Adaptive Poisson-Boltzmann Solver (APBS) software suite in the following conditions: pH 7.0, 298.15 K, 150 mM NaCl, 1.1 and 1.7 Å ionic radii for Na+ and Cl-, respectively. All calculation steps were carried out via the APBS/PDB2PQR website (47). Resulting Poisson-Boltzmann electrostatic potentials were mapped on the van der Waals surface of LolA and LolB proteins, using a potential scale ranging from −3 k_B_T/e to +3 k_B_T/e. Regarding hydrophobicity computation, residues were colored according to the Eisenberg’s normalized consensus hydrophobicity scale using the Color h script in the PyMOL software (48). Scores were mapped on the van der Waals surface of LolA and LolB proteins. All the structures were visualized using the PyMOL Molecular Graphics System, Version 1.2r3pre, Schrödinger, LLC.

### Construction of *F. johnsoniae* deletion strains

Suicide plasmids (**Supplementary Table S6**) for deletions in *F. johnsoniae* were constructed by amplifying the 2 kb chromosomal regions upstream and downstream of the target gene with specific primers (**Supplementary Table S7**) and cloned sequentially (or by Gibson assembly (49) (NEB)) into the pYT354 suicide plasmid. Suicide plasmids were introduced into the appropriate *F. johnsoniae* background strain by triparental mating using *E. coli* Top10 as donor and *E. coli* MT607 as helper strain. Erythromycin resistance was used to select cells with chromosomally integrated plasmid. One of the resulting clones was grown overnight in CYE without antibiotics to allow for loss of the plasmid backbone and then plated on CYE agar containing 5 % sucrose. Sucrose-resistant colonies were screened by PCR for the presence of the desired chromosomal modification.

### Construction of plasmids for complementation in *F. johnsoniae* and *C. canimorsus*

Genes of interest were amplified from genomic DNA using specific primers (**Supplementary Table S7**) with Q5 DNA Polymerase. The expression plasmids pCP23-P*ermF* or pMM47.A and the amplified genes were digested with appropriate restriction enzymes (New England Biolabs) and ligated with T4 ligase (New England Biolabs) O/N at 16°C. The ligation products were transformed by heat-shock or electroporation into *E. coli* Top10 and plated on LB Amp plates. Resistant clones were screened by PCR for the presence of the correct plasmid and constructs were checked by sequencing. Plasmids (**Supplementary Table S6**) were transferred to *F. johnsoniae* by triparental mating and tetracycline resistance was used to select for cells that received the plasmid. Plasmids were transferred to *C. canimorsus* by electroporation and cefoxitin resistance was used to select for cells that received the plasmid.

### Motility assays

To assess gliding motility of *F. johnsoniae,* strains were grown O/N in CYE at 30°C. A volume of the cultures corresponding to an OD_600_ of 0.1 was collected and centrifuged for 3 minutes at 5,000 × g. The resulting pellets were resuspended in 1 ml of PBS. 3 μl of the resuspended cells were spotted on MM plates and incubated 48h at 30°C. Pictures were taken with a Nikon CoolPix L29 camera.

### Amylase activity assays

*F. johnsoniae* strains were grown O/N in CYE at 30°C. For plate amylase activity assays, a volume of the cultures corresponding to an OD^600^ of 0.1 was collected and centrifuged for 3 minutes at 5,000 × g. The resulting pellets were resuspended in 1 ml of PBS. 3 μl of the resuspended cells were spotted on LB (0.2% starch (Merck)) and were incubated 24h at 30°C. A solution of 1% KI and 1% iodine was poured on the agar for starch staining and pictures were taken with a Nikon CoolPix L29 camera. For quantitative amylase activity assays, bacterial cultures were diluted to an OD^600^ of 0.05 in CYE and grown in a microplate reader in the same conditions as for the growth assays. All the strains were grown for 12 hours and, after checking that they had all reached the same OD_600_, cultures were retrieved and centrifuged for 3 minutes at 5,000 × g. Supernatant was collected and filtered with 0.2 µm membranes to remove bacteria. Filtered supernatant was mixed with a 1% starch solution (0.2% final) and incubated for 4 hours at 30°C, 160 rpm. Finally, 143 µl of the solution was mixed with 107 µl of DNSA 96 mM, boiled at 99°C for 20 minutes, and absorbance was read at 540 nm with a SpectraMax iD3 microplate reader (Molecular Devices).

### Membrane purification

Membranes were purified as previously described with some modifications (22) *F. johnsoniae* liquid cultures (50 ml) of the WT, *lolA1* mutant, and *lolB1* mutant strains were grown O/N in CYE at 30°C, 160 rpm. The equivalent of an OD_600_ of 40 was collected through centrifugation (10 minutes at 7,000 × g, 4°C), resuspended in 4 ml of a buffer (HEPES 10 mM pH 7.4, 1 mM EDTA, EDTA-free protease inhibitor cocktail from a tablet (Roche)) and lysed by 3 passages through a cell disrupter at 2.4 kbar. The lysates were then centrifuged 10 minutes at 2,500 × g, 4°C, to pellet unbroken cells and debris. The inner and outer membranes were precipitated through a first ultracentrifugation step (107 minutes at 55,000 rpm with a Beckman TLA-100.3 fixed-angle rotor, 4°C). To solubilize the inner membrane, the pellets were resuspended in a solution of HEPES 10 mM pH 7.4 and N-lauroylsarcosine sodium salt 2% (sarcosyl) (Sigma-Aldrich) and were incubated on a roller shaker for 60 minutes at RT. To separate the membranes, a second ultracentrifugation step (identical settings as the first) was performed, and the outer membrane pellet was resuspended in 250 µl of HEPES 10 mM pH 7.4.

### Verification of the purity of the membrane fractions

To verify for inner membrane contamination in the outer membrane fraction, a SDH activity assay was carried out (50). During the OM purification, 200 µl were sampled right after cell lysis (total lysate fraction), 200 µl of supernatant right after the first ultracentrifugation step (periplasm + cytoplasm fraction), and 200 µl after full resuspension of the inner and outer membranes in HEPES (inner and outer membrane fraction). Inner membrane and outer membrane fractions were stored at the end of the purification. After membrane extraction, the protein concentration of the different fractions was quantified with a Bradford protein assay (Bio-rad). The fractions were diluted to 0.24 µg/µl of protein. A 96-well plate was filled with either 100 μl of sample or water. Then, 60 μl of the following reaction mix (final concentrations) was added to each well: 50 mM Tris-HCl (pH 8.0), 4 mM KCN, and 40 mM disodium succinate. After 5 minutes of incubation at RT, 20 μl of 4 mM DCIP and 20 µl of 2 mM PMS were subsequently added. Absorbance at 600 nm was measured using a SpectraMax iD3 microplate reader (Molecular Devices) every 36 seconds during 1h at 25°C. As an additional verification of outer membrane purity, 2 μg of protein were loaded on a 12% SDS-PAGE gel and visualized by silver staining. OM fractions from five independent biological replicates with low SDH activity and characteristic silver staining OM band patterns were analysed by mass spectrometry.

### Protein digestion prior MS

The samples were treated using Filter-Aided Sample Preparation (FASP) using the following protocol:

To first wash the filter, 100 µl of formic acid 1% were placed in each Millipore Microcon 30 MRCFOR030 Ultracel PL-30 before centrifugation at 14,500 rpm (Eppendorf 5424 centrifuge) for 15 minutes. For each sample, 15 µg of proteins adjusted in 150 µl of urea buffer 8 M (urea 8 M in buffer Tris 0.1 M at pH 8.5) were placed individually in a column and centrifuged at 14,500 rpm for 15 minutes. The filtrate was discarded, and the columns were washed three times by adding 200 µl of urea buffer followed by a centrifugation at 14,500 rpm for 15 minutes. For the reduction step, 100 µl of dithiothreitol (DTT) were added and mixed for 1 minute at 400 rpm with a thermomixer before an incubation of 15 minutes at 24 °C. Samples were then centrifuged at 14,500 rpm for 15 minutes, the filtrate was discarded, and the filter was washed by adding 100 µl of urea buffer before another centrifugation at 14,500 rpm for 15 minutes. An alkylation step was performed by adding 100 µl of iodoacetamide ((IAA), in urea buffer) in the column and mixing at 400 rpm for 1 minute in the dark before an incubation of 20 minutes in the dark and a centrifugation at 14,500 rpm for 15 minutes. To remove the excess of IAA, 100 µl of urea buffer were added and the samples were centrifuged at 14,500 rpm for 15 minutes. To quench the rest of IAA, 100 µl of DTT were placed on the column, mixed for 1 minute at 400 rpm and incubated for 15 minutes at 24 °C before centrifugation at 14,500 rpm for 15 minutes. To remove the excess of DTT, 100 µl of urea buffer were placed on the column and centrifuged at 14,500 rpm for 15 minutes. The filtrate was discarded, and the column was washed three times by adding 100 µl of sodium bicarbonate buffer 50 mM ((ABC), in ultrapure water) followed by a centrifugation at 14,500 rpm for 15 minutes. The last 100 µl were kept at the bottom of the column to avoid evaporation. The digestion process was performed by adding 80 µl of mass spectrometry-grade trypsin (1/50 in ABC buffer) in the column and mixed at 300 rpm overnight at 24°C, in the dark. The Microcon columns were placed on LoBind tubes of 1.5 ml and centrifuged at 14,500 rpm for 15 minutes. 40 µl of ABC buffer were placed on the column before centrifugation at 14,500 rpm for 15 minutes. 2 µl of trifluoroacetic acid (TFA) 10 % in ultrapure water were added to the filtrate (0.2 % TFA final). The samples were dried in a SpeedVac, resuspended in injection solvent (acetonitrile (ACN) 2%, formic acid 0.1%) for a final concentration of 250 ng/µl, and transferred to an injection vial.

### Mass Spectrometry

The digest was analysed using nano-LC-ESI-MS/MS tims TOF Pro (Bruker, Billerica, MA, USA) coupled with an UHPLC nanoElute2 (Bruker). The different samples were analysed with a gradient of 60 min. Peptides were separated by nanoUHPLC (nanoElute2, Bruker) on a 75 μm ID, 25 cm C18 column with integrated CaptiveSpray insert (Aurora, ionopticks, Melbourne) at a flow rate of 200 nl/min, at 50°C. LC mobile phases A was water with 0.1% formic acid (v/v) and B ACN with formic acid 0.1% (v/v). Samples were loaded directly on the analytical column at a constant pressure of 800 bar. The digest (1 µl) was injected, and the organic content of the mobile phase was increased linearly from 2% B to 15% in 22 min, from 15% B to 35% in 38 min, from 35% B to 85% in 3 min. Data acquisition on the tims TOF Pro was performed using Hystar 6.1 and timsControl 2.0. tims TOF Pro data were acquired using 160 ms TIMS accumulation time, mobility (1/K0) range from 0.75 to 1.42 Vs/cm². Mass-spectrometric analysis was carried out using the parallel accumulation serial fragmentation (PASEF) (51) acquisition method. One MS spectra followed by six PASEF MSMS spectra per total cycle of 1.16 s.

### PEAKS proteomic analysis

Data analysis was performed using PEAKS Studio 11 with ion mobility module and Q module for label-free quantification (Bioinformatics Solutions Inc., Waterloo, ON). Protein identifications were conducted using PEAKS search engine with 20 ppm as parent mass error tolerance and 0.05 Da as fragment mass error tolerance. Carbamidomethylation was allowed as fixed modification, oxidation of methionine and acetylation (N-term) as variable modification. Enzyme specificity was set to trypsin, and the maximum number of missed cleavages per peptide was set to two. The peak lists were searched against the *Flavobacterium johnsoniae* UW101 proteome from Uniprot (220718-5,021 entries) and a contaminants database. Peptide spectrum matches and protein identifications were normalized to less than 1.0% false discovery rate.

Label-free quantitation (LFQ) method is based on expectation - maximization algorithm on the extracted ion chromatograms of the three most abundant unique peptides of a protein to calculate the area under the curve (52). For the quantitation, mass error and ion mobility tolerance were set respectively to 20 ppm and 0.08 1/k0. For the label-free quantitation results, peptide quality score was set to be ≥ 20 and protein significance score threshold was set to 20. The significance score is calculated as the −10log10 of the significance testing p-value (0.01). ANOVA was used as the significance testing method. Modified peptides were excluded and only proteins with at least two peptides were used for the quantitation. Total ion current was used to calculate the normalization factors.

### In *silico* characterization of the proteins identified by label-free mass spectrometry

The presence of signal peptide I or II was predicted using the SignalP 6.0 server (46). Proteins devoid of any signal peptide were excluded from further analyses. Proteins with a SPII (lipoproteins) were manually checked for the presence of the LES (surface lipoproteins) (16, 17). Localization of the proteins was manually checked, according to the signal peptide, the predicted 3D structure, Interpro domains, and prediction by PSORTb 3.0.3 (53). Proteins belonging to SUS-like systems were annotated with the CAZy database (54). Essentiality of the proteins was determined thanks to a transposon sequencing (Tn_seq) analysis previously realized in our group. Proteins were functionally annotated using the MicroScope platform (55), which employs data from EggNOG 5.0 (56) and EggNOG-mapper 2.1 (57).

### Growth and OM stress tolerance assays

*F. johnsoniae* strains were grown O/N in CYE at 30°C, 160 rpm. Cultures were diluted to an OD_600_ of 0.05 and washed once with PBS. For growth assays, cells were resuspended in fresh CYE or MM. For stress tolerance assays, cells were resuspended in fresh CYE in the absence or presence of the following compounds: Polymyxin B (2, 20 and 40 μg/ml), EDTA (0.2, 0.5 and 1 mM) and SDS (0.001 and 0.005%). Optical density at 600 nm was measured every 10 minutes during 24h at 30°C with continuous linear shaking (567 cpm) using an EPOCH2 Microplate Reader (BioTek).

### Phase-contrast microscopy

For live imaging, cells were grown O/N in regular CYE. Cultures were then diluted to an OD_600_ of 0.05 in 10 ml of CYE low magnesium (4 mM MgSO_4_) or MM, in flasks. Cells were incubated for 16 hours at 30°C, 160 rpm, and directly spotted on a microscope glass slide. Pictures were acquired with an Axio Observer (Zeiss) microscope equipped with an Orca-Flash 4.0 camera (Hamamatsu) and the Zen Pro 3.9 software (Zeiss).

### Transmission Electron Microscopy

Bacterial samples for electron microscopy were prepared as previously described with some modifications (58). *F. johnsoniae* was grown O/N in regular CYE. Cultures were then diluted to an OD_600_ of 0.05 in 10 ml of CYE low magnesium (4 mM MgSO_4_) in flasks and incubated for 16 hours at 30°C, 160 rpm. The fixation procedure was performed as follows: 2 ml of bacterial culture were centrifuged for 3 minutes at 5,000 × g and the pellet was resuspended in 400 µl of a solution of glutaraldehyde 2.5% and cacodylate 0.1 % (pH 7.4). After 2h30 of incubation at 4°C, bacteria were washed three times with cacodylate 0.2%, resuspended in a solution of osmic acid 2% and cacodylate 0.1%, and incubated for one additional hour at 4°C. Then, bacteria were washed two more times with cacodylate 0.2%. Samples were dehydrated by incubation with increasing concentrations of ethanol (30%, 50%, 70%, 85%, and 100%) at RT. For the embedding in resin, samples were first washed four times with propylene oxide and then incubated in three propylene oxide/resin solutions: 15 minutes in 75% propylene oxide/25% resin, 20 minutes in 50% propylene oxide/50% resin, and 20 minutes in 25% propylene oxide/75% resin. Finally, pellets were embedded in pure epoxy resin. Resin was dried by incubation at 37°C O/N, 24h of incubation at 45°C, and three days at 60°C. Ultrathin sections were cut with a DiATOME ultra 45° diamond knife and visualized with a FEI Tecnai T10 electron microscope at 60-80 kV. Images were acquired by the software Item of Olympus with an Olympus Megaview 1024x1024 pixels camera.

### Complementation of *lolB1* mutant with mutated variants of LolB1

*lolB1 (Fjoh_1066)* was amplified with oligonucleotides containing the desired mutation (L74E, Δ73-76, C17G) (**Supplementary Table S7**). For the deletion of the C-terminal domain (223-243), Phe222 was substituted with a stop codon. The mutated genes were then cloned into pCP23-P*ermF*, and the resulting plasmids transferred to the *F. johnsoniae lolB1* mutant by triparental mating. To add a 6xHis tag at the C-terminus of the WT and mutated LolB1 variants, the genes were amplified with a forward oligonucleotide and a reverse one devoid of the stop codon (**Supplementary Table S7**) and cloned into pCP23-P*ermF* in frame with the 6XHis tag encoded in the plasmid.

### Detection of His-tagged LolB variants and SiaC by Western-Blot

For Western Blot analyses, bacteria were grown in CYE supplemented with tetracycline till stationary phase and a volume of bacterial culture (grown O/N in CYE) equivalent to an OD_600_ of 1 was centrifuged for 3 minutes at 5,000 × g. The pellet was resuspended in 80 µl of sterile water and 20 µl of SDS-PAGE buffer 5x. After heating the samples for 5 minutes at 98°C, 10 μl were loaded on a 12% SDS-PAGE gel. After migration, the proteins were transferred onto a nitrocellulose membrane with a Trans-Blot Turbo Transfer System (Bio-Rad). Proteins were detected with rabbit anti-His (1/5,000) (PM032, MBL), rabbit anti-GroEL (1/1,600) (G6535, Sigma-Aldrich), or rabbit anti-SiaC (1/5,000) primary antibodies and polyclonal swine-HRP anti-rabbit (1/5,000) (P0217, Dako) secondary antibodies. Membranes were revealed using a KPL LumiGLO Reserve Chemiluminescent Substrate Kit (SeraCare) and images were captured using a GE Healthcare Amersham Imager 600.

### Deletion of *lolA* and *lolB* and heterologous complementation in *E. coli*

For the deletion of *lolA* and *lolB* in *E. coli MG1655*, forward and reverse primers were designed respectively with the start and end of the gene of interest sequence flanked with regions that anneal on the pKD4 plasmid containing the Kan^R^ gene to be amplified: oligonucleotides 8675/8679 and 8677/8680 for *lolA* and *lolB* deletion respectively **(Supplementary Table S7**) (59). The *E. coli* MG1655 mini-λ-Tet strain was transformed with the pBAD33-Ara inducible plasmid expressing the following genes: *lolA, lolB, lolA1*, *lolB1*, *lolA1* and *lolB1*, *lolA2*, *lolB2*, *lolA2* and *lolB2*. The plasmid-borne complemented strains were grown O/N at 30°C in LB Tet Cm and 1 ml of the O/N liquid culture was transferred to 100 ml of LB Tet Cm.

The λ red system was then induced by placing the cultures in a 42°C water bath for 15 minutes at 220 rpm. Cells were then made electro-competent and transformed with PCR fragments for the deletion of either *lolA* or *lolB* chromosomal *loci* by electroporation, then resuspended in 1 ml of LB 0.2% arabinose (Ara) and incubated for 1h at 37°C. 100 μl of the cells were then plated on LB Kan Cm 0.2% Ara plates. When present, colonies grown after 24h and/or 48h were checked for deletion of either *lolA* or *lolB* by PCR.

### Conservation of LolA and LolB in Bacteroidetes

A Delta-blast (36) sequence similarity search was conducted on the 30 Bacteroidetes species: *Bacteroides fragilis, Bacteroides ovatus, Bacteroides thetaioathaomicron, Bergeyella zoohelcum, Capnocytophaga canimorsus, Capnocytophaga canis, Capnocytophaga cynodegmi, Capnocytophaga gingivalis, Capnocytophaga ochracea, Chitinophaga filiformis, Chitinophaga pinensis, Christiangramia forsetii, Croceibacter atlanticus, Cytophaga hutchinsonii, Flavobacterium columnare, Flavobacterium johnsoniae, Flavobacterium meningosepticum, Flavobacterium psychrophilum, Flavobacterium succinicans, Flexibacter flexilis, Kordia algicida, Polaribacter irgensii, Porphyromonas gingivalis, Prevotella intermedia, Prevotella melaninogenica, Riemerella anatipestifer, Sphingobacterium mizutaii, Sporocytophaga myxococcoides, Xanthomarina gelatinilytica, Zobellia galactanivorans*, using the five LolA and LolB protein sequences of *F. johnsoniae ATCC 17061 UW101* as queries: LolA1 (WP_081432686.1), LolA2 (WP_012023170.1), LolA3 (WP_012022695.1), LolB1 (WP_012023151.1) and LolB2 (WP_012023169.1). The genomic neighbourhood of all hits with a Delta-blast E value ≤ 0.001 was inspected using MaGe (60). The structure of the corresponding proteins was predicted using AlphaFold server (with default parameters) (45) and compared with the corresponding structures of the of *F. johnsoniae* initial queries by manual inspection. This resulted in a list of 93 putative homologs. The phylogenetic tree based on NCBI taxonomy was generated via phyloT (61, 62). Conservation of the C-terminal domain of LolB1 homologs was performed by manual inspection on AlphaFold models.

### Deletion of *lolA* and *lolB* and heterologous complementation in *C. canimorsus*

To obtain the *C. canimorsus lolA* (*Ccan_16490*) mutant strain expressing *F. johnsoniae lolA1*, we transformed the pMM47-*lolA1* (Cfx^R^) plasmid into *C. canimorsus lolA* harboring pFL63 plasmid (Tet^R^) which expresses WT *C. canimorsus lolA*). We selected cefoxitin-resistant colonies and subcultured them 3 times on cefoxitin-containing SB plates. Colonies were checked for tetracycline sensitivity and loss of the pFL63 plasmid was confirmed by PCR. To obtain the *C. canimorsus lolB* (*Ccan_17050*) mutant strain expressing *F. johnsoniae lolB1* we first transformed plasmid pMM47-lolB1 into WT *C. canimorsus* and then deleted gene *Ccan_17050.* Deletion of *Ccan_17050* in the *C. canimorsus lolB1*-expressing strain was performed by amplification and cloning into pYT354 suicide plasmid of the 500 bases of the chromosomal regions upstream and downstream of *Ccan_17050* (**Supplementary Table S6** and **S7**). Suicide plasmid was introduced into the *C. canimorsus* pMM47-*lolB1* strain by triparental mating using *E. coli* Top10 as donor and *E. coli* MT607 as helper strain. Erythromycin resistance was used to select cells with chromosomally integrated plasmid. One of the resulting clones was grown overnight on a SB plate without antibiotics to allow for loss of the plasmid backbone and then plated on a SB plate containing 3 % sucrose. Sucrose-resistant colonies were screened by PCR for the presence of the *Ccan_17050* deletion.

### Immunofluorescence microscopy of sialidase

*F. johnsoniae* WT and *lolA1 lolA2 lolA3 lolB1 lolB2* strains harbouring plasmid pCP23-P*ermF*-*siaC* or pCP23-P*ermF*-LES-*siaC* were grown O/N in CYE Tet at 30°C. A volume equivalent to an OD_600_ of 0.1 was centrifuged 3 minutes at 5,000 × g and resuspended in 1 ml of PBS. Cells were collected by a centrifugation of 5 minutes at 4,000 × g and resuspended in 200 μl of a PBS-BSA 1% solution and incubated for 30 minutes at RT. Cells were centrifuged 5 minutes at 4,000 × g and resuspended in 200 μl of PBS 1/2,000 diluted rabbit anti-SiaC serum and incubated for 30 minutes at RT. Cells were washed 3 times with PBS and resuspended in 200 μl of PBS 1/500 diluted donkey anti-rabbit AlexaFluor-488TM (Thermofisher) and incubated 30 minutes at RT in the dark. After another 3 washes with PBS, cells were fixed with 200 μl of PFA 4% for 15 minutes in the dark. Cells were washed once with PBS and stored at 4°C prior analysis. Labelled bacteria were spotted on 1% agarose PBS pads and pictures were taken using an Axio Observer (Zeiss) microscope equipped with an Orca-Flash 4.0 camera (Hamamatsu) and the Zen Pro 3.9 software (Zeiss).

### Statistical analyses

Results are presented as means with standard deviations. OM stress tolerance assay results (**Figure 5**) were analyzed using a one-way ANOVA followed by Tukey’s multiple comparison test with a p-value ≤ 0.05 considered significant in GraphPad Prism version 8.0.1 (https://www.graphpad.com). For quantitative amylase activity assays, a one-way ANOVA followed by Dunnett’s multiple comparison test was used.

## Accession codes

Label free quantitative mass spectrometry data have been deposited in the PRIDE PRoteomics IDEntification Database.

## Acknowledgements

We are grateful to Isabelle Hamer (UNamur) for providing help with ultracentrifuges and to Paul Guiraud (UNamur) for his help with optical microscopy. We thank Gwennaëlle Louis for her help on amylase activity assays set up and Catherine Demazy (UNamur) for help on samples preparation for proteomics. We are grateful to the Electron microscopy facility of UNamur. This research has been funded by the Incentive Grant for Scientific Research (MIS F.4533.20F) from the *Fonds de la Recherche Scientifique-Fonds National de la Recherche Scientifique* (FRS-FNRS, http://www.fnrs.be) to F. Renzi. T. De Smet is founded by a PhD fellowship (Aspirant) from the FRS-FNRS. F. Renzi is a research associate of the FRS-FNRS.

## Author Contributions

T. D. S., E. B., D. D. and F. R. designed research; T. D. S., E. B., F. G., J. M., L. L., R. D.,F. L., M. D., G. L-M., C. M., D. D. and F. R. performed research; T. D. S., E. B., J. M., M. D., G. L-M., C. M., D. D. and F. R. analyzed data; and T. D. S., J. M., C. M. and F. R. wrote the paper.

## References

1. Silhavy TJ, Kahne D, Walker S. The bacterial cell envelope. Cold Spring Harb Perspect Biol. 2010;2(5):a000414.

2. Braun V. Covalent lipoprotein from the outer membrane of Escherichia coli. Biochim Biophys Acta. 1975;415(3):335–77.

3. Knowles TJ, Scott-Tucker A, Overduin M, Henderson IR. Membrane protein architects: the role of the BAM complex in outer membrane protein assembly. Nat Rev Microbiol. 2009;7(3):206–14.

4. Sperandeo P, Martorana AM, Polissi A. The lipopolysaccharide transport (Lpt) machinery: A nonconventional transporter for lipopolysaccharide assembly at the outer membrane of Gram-negative bacteria. J Biol Chem. 2017;292(44):17981–90.

5. Manfredi P, Renzi F, Mally M, Sauteur L, Schmaler M, Moes S, et al. The genome and surface proteome of Capnocytophaga canimorsus reveal a key role of glycan foraging systems in host glycoproteins deglycosylation. Mol Microbiol. 2011;81(4):1050–60.

6. Wilson MM, Anderson DE, Bernstein HD. Analysis of the outer membrane proteome and secretome of Bacteroides fragilis reveals a multiplicity of secretion mechanisms. PLoS One. 2015;10(2):e0117732.

7. Wilson MM, Bernstein HD. Surface-Exposed Lipoproteins: An Emerging Secretion Phenomenon in Gram-Negative Bacteria. Trends Microbiol. 2016;24(3):198–208.

8. Johnston JJ, Shrivastava A, McBride MJ. Untangling Flavobacterium johnsoniae Gliding Motility and Protein Secretion. J Bacteriol. 2018;200(2).

9. Zuckert WR. Secretion of bacterial lipoproteins: through the cytoplasmic membrane, the periplasm and beyond. Biochim Biophys Acta. 2014;1843(8):1509–16.

10. Bos MP, Robert V, Tommassen J. Biogenesis of the gram-negative bacterial outer membrane. Annu Rev Microbiol. 2007;61:191–214.

11. Okuda S, Tokuda H. Lipoprotein sorting in bacteria. Annu Rev Microbiol. 2011;65:239–59.

12. Sutcliffe IC, Harrington DJ, Hutchings MI. A phylum level analysis reveals lipoprotein biosynthesis to be a fundamental property of bacteria. Protein Cell. 2012;3(3):163–70.

13. Smith HC, May KL, Grabowicz M. Teasing apart the evolution of lipoprotein trafficking in gram-negative bacteria reveals a bifunctional LolA. Proc Natl Acad Sci U S A. 2023;120(6):e2218473120.

14. Murphy BT, Wiepen JJ, Graham DE, Swanson SK, Kashipathy MM, Cooper A, et al. Borrelia burgdorferi BB0346 is an Essential, Structurally Variant LolA Homolog that is Primarily Required for Homeostatic Localization of Periplasmic Lipoproteins. bioRxiv. 2024.

15. He H, Pramanik AS, Swanson SK, Johnson DK, Florens L, Zuckert WR. A Borrelia burgdorferi LptD homolog is required for flipping of surface lipoproteins through the spirochetal outer membrane. Mol Microbiol. 2023;119(6):752–67.

16. Lauber F, Cornelis GR, Renzi F. Erratum for Lauber, et al., Identification of a New Lipoprotein Export Signal in Gram-Negative Bacteria. mBio. 2016;7(6).

17. Lauber F, Cornelis GR, Renzi F. Identification of a New Lipoprotein Export Signal in Gram-Negative Bacteria. mBio. 2016;7(5).

18. Rhodes RG, Samarasam MN, Van Groll EJ, McBride MJ. Mutations in Flavobacterium johnsoniae sprE result in defects in gliding motility and protein secretion. J Bacteriol. 2011;193(19):5322–7.

19. Schoner TA, Fuchs SW, Schonau C, Bode HB. Initiation of the flexirubin biosynthesis in Chitinophaga pinensis. Microb Biotechnol. 2014;7(3):232–41.

20. Pugsley AP. The complete general secretory pathway in gram-negative bacteria. Microbiol Rev. 1993;57(1):50–108.

21. Nakada S, Sakakura M, Takahashi H, Okuda S, Tokuda H, Shimada I. Structural investigation of the interaction between LolA and LolB using NMR. J Biol Chem. 2009;284(36):24634–43.

22. Hunnicutt DW, McBride MJ. Cloning and characterization of the Flavobacterium johnsoniae gliding-motility genes gldB and gldC. J Bacteriol. 2000;182(4):911–8.

23. Braun TF, McBride MJ. Flavobacterium johnsoniae GldJ is a lipoprotein that is required for gliding motility. J Bacteriol. 2005;187(8):2628–37.

24. Kharade SS, McBride MJ. Flavobacterium johnsoniae PorV is required for secretion of a subset of proteins targeted to the type IX secretion system. J Bacteriol. 2015;197(1):147–58.

25. Grondin JM, Tamura K, Dejean G, Abbott DW, Brumer H. Polysaccharide Utilization Loci: Fueling Microbial Communities. J Bacteriol. 2017;199(15).

26. Rhodes RG, Samarasam MN, Shrivastava A, van Baaren JM, Pochiraju S, Bollampalli S, et al. Flavobacterium johnsoniae gldN and gldO are partially redundant genes required for gliding motility and surface localization of SprB. J Bacteriol. 2010;192(5):1201–11.

27. Shrivastava A, Johnston JJ, van Baaren JM, McBride MJ. Flavobacterium johnsoniae GldK, GldL, GldM, and SprA are required for secretion of the cell surface gliding motility adhesins SprB and RemA. J Bacteriol. 2013;195(14):3201–12.

28. Gorasia DG, Veith PD, Hanssen EG, Glew MD, Sato K, Yukitake H, et al. Structural Insights into the PorK and PorN Components of the Porphyromonas gingivalis Type IX Secretion System. PLoS Pathog. 2016;12(8):e1005820.

29. Sato K, Naito M, Yukitake H, Hirakawa H, Shoji M, McBride MJ, et al. A protein secretion system linked to bacteroidete gliding motility and pathogenesis. Proc Natl Acad Sci U S A. 2010;107(1):276–81.

30. Rhodes RG, Nelson SS, Pochiraju S, McBride MJ. Flavobacterium johnsoniae sprB is part of an operon spanning the additional gliding motility genes sprC, sprD, and sprF. J Bacteriol. 2011;193(3):599–610.

31. Liu J, McBride MJ, Subramaniam S. Cell surface filaments of the gliding bacterium Flavobacterium johnsoniae revealed by cryo-electron tomography. J Bacteriol. 2007;189(20):7503–6.

32. Clifton LA, Holt SA, Hughes AV, Daulton EL, Arunmanee W, Heinrich F, et al. An Accurate In Vitro Model of the E. coli Envelope. Angew Chem Weinheim Bergstr Ger. 2015;127(41):12120–3.

33. Hayashi Y, Tsurumizu R, Tsukahara J, Takeda K, Narita SI, Mori M, et al. Roles of the protruding loop of factor B essential for the localization of lipoproteins (LolB) in the anchoring of bacterial triacylated proteins to the outer membrane. J Biol Chem. 2014;289(15):10530–9.

34. Tsukahara J, Mukaiyama K, Okuda S, Narita S, Tokuda H. Dissection of LolB function--lipoprotein binding, membrane targeting and incorporation of lipoproteins into lipid bilayers. FEBS J. 2009;276(16):4496–504.

35. Butler T. Capnocytophaga canimorsus: an emerging cause of sepsis, meningitis, and post-splenectomy infection after dog bites. Eur J Clin Microbiol Infect Dis. 2015;34(7):1271–80.

36. Boratyn GM, Schaffer AA, Agarwala R, Altschul SF, Lipman DJ, Madden TL. Domain enhanced lookup time accelerated BLAST. Biol Direct. 2012;7:12.

37. Letunic I, Doerks T, Bork P. SMART 7: recent updates to the protein domain annotation resource. Nucleic Acids Res. 2012;40(Database issue):D302–5.

38. Mally M, Shin H, Paroz C, Landmann R, Cornelis GR. Capnocytophaga canimorsus: a human pathogen feeding at the surface of epithelial cells and phagocytes. PLoS Pathog. 2008;4(9):e1000164.

39. Liao CT, Chiang YC, Hsiao YM. Functional characterization and proteomic analysis of lolA in Xanthomonas campestris pv. campestris. BMC Microbiol. 2019;19(1):20.

40. Fernandez-Pinar R, Lo Sciuto A, Rossi A, Ranucci S, Bragonzi A, Imperi F. In vitro and in vivo screening for novel essential cell-envelope proteins in Pseudomonas aeruginosa. Sci Rep. 2015;5:17593.

41. Grabowicz M, Silhavy TJ. Redefining the essential trafficking pathway for outer membrane lipoproteins. Proc Natl Acad Sci U S A. 2017;114(18):4769–74.

42. Malinverni JC, Werner J, Kim S, Sklar JG, Kahne D, Misra R, et al. YfiO stabilizes the YaeT complex and is essential for outer membrane protein assembly in Escherichia coli. Mol Microbiol. 2006;61(1):151–64.

43. Steinegger M, Meier M, Mirdita M, Vohringer H, Haunsberger SJ, Soding J. HH-suite3 for fast remote homology detection and deep protein annotation. BMC Bioinformatics. 2019;20(1):473.

44. Zhou X, Zheng W, Li Y, Pearce R, Zhang C, Bell EW, et al. I-TASSER-MTD: a deep-learning-based platform for multi-domain protein structure and function prediction. Nat Protoc. 2022;17(10):2326–53.

45. Abramson J, Adler J, Dunger J, Evans R, Green T, Pritzel A, et al. Accurate structure prediction of biomolecular interactions with AlphaFold 3. Nature. 2024;630(8016):493–500.

46. Teufel F, Almagro Armenteros JJ, Johansen AR, Gislason MH, Pihl SI, Tsirigos KD, et al. SignalP 6.0 predicts all five types of signal peptides using protein language models. Nat Biotechnol. 2022;40(7):1023–5.

47. Jurrus E, Engel D, Star K, Monson K, Brandi J, Felberg LE, et al. Improvements to the APBS biomolecular solvation software suite. Protein Sci. 2018;27(1):112–28.

48. Eisenberg D, Schwarz E, Komaromy M, Wall R. Analysis of membrane and surface protein sequences with the hydrophobic moment plot. J Mol Biol. 1984;179(1):125–42.

49. Gibson DG, Young L, Chuang RY, Venter JC, Hutchison CA, 3rd, Smith HO. Enzymatic assembly of DNA molecules up to several hundred kilobases. Nat Methods. 2009;6(5):343–5.

50. Kasahara M, Anraku Y. Succinate Dehydrogenase of *Escherichia coli* Membrane Vesicles Activation and Properties of the Enzyme. The Journal of Biochemistry. 1974;76(5):959–66.

51. Meier F, Brunner AD, Koch S, Koch H, Lubeck M, Krause M, et al. Online Parallel Accumulation-Serial Fragmentation (PASEF) with a Novel Trapped Ion Mobility Mass Spectrometer. Mol Cell Proteomics. 2018;17(12):2534–45.

52. Lin H, He L, Ma B. A combinatorial approach to the peptide feature matching problem for label-free quantification. Bioinformatics. 2013;29(14):1768–75.

53. Yu NY, Wagner JR, Laird MR, Melli G, Rey S, Lo R, et al. PSORTb 3.0: improved protein subcellular localization prediction with refined localization subcategories and predictive capabilities for all prokaryotes. Bioinformatics. 2010;26(13):1608–15.

54. Drula E, Garron ML, Dogan S, Lombard V, Henrissat B, Terrapon N. The carbohydrate-active enzyme database: functions and literature. Nucleic Acids Res. 2022;50(D1):D571–D7.

55. Vallenet D, Calteau A, Dubois M, Amours P, Bazin A, Beuvin M, et al. MicroScope: an integrated platform for the annotation and exploration of microbial gene functions through genomic, pangenomic and metabolic comparative analysis. Nucleic Acids Res. 2020;48(D1):D579–D89.

56. Huerta-Cepas J, Szklarczyk D, Heller D, Hernandez-Plaza A, Forslund SK, Cook H, et al. eggNOG 5.0: a hierarchical, functionally and phylogenetically annotated orthology resource based on 5090 organisms and 2502 viruses. Nucleic Acids Res. 2019;47(D1):D309–D14.

57. Cantalapiedra CP, Hernandez-Plaza A, Letunic I, Bork P, Huerta-Cepas J. eggNOG-mapper v2: Functional Annotation, Orthology Assignments, and Domain Prediction at the Metagenomic Scale. Mol Biol Evol. 2021;38(12):5825–9.

58. Wenzel M, Dekker MP, Wang B, Burggraaf MJ, Bitter W, van Weering JRT, et al. A flat embedding method for transmission electron microscopy reveals an unknown mechanism of tetracycline. Commun Biol. 2021;4(1):306.

59. Court DL, Swaminathan S, Yu D, Wilson H, Baker T, Bubunenko M, et al. Mini-lambda: a tractable system for chromosome and BAC engineering. Gene. 2003;315:63–9.

60. Vallenet D, Labarre L, Rouy Z, Barbe V, Bocs S, Cruveiller S, et al. MaGe: a microbial genome annotation system supported by synteny results. Nucleic Acids Res. 2006;34(1):53–65.

61. Letunic I, Bork P. Interactive Tree Of Life (iTOL): an online tool for phylogenetic tree display and annotation. Bioinformatics. 2007;23(1):127–8.

62. Letunic I, Bork P. Interactive Tree Of Life v2: online annotation and display of phylogenetic trees made easy. Nucleic Acids Res. 2011;39(Web Server issue):W475–8.

