## Supplementary Figures and Tables for "LolA and LolB are conserved in Bacteroidetes and are crucial for gliding motility and Type IX secretion"

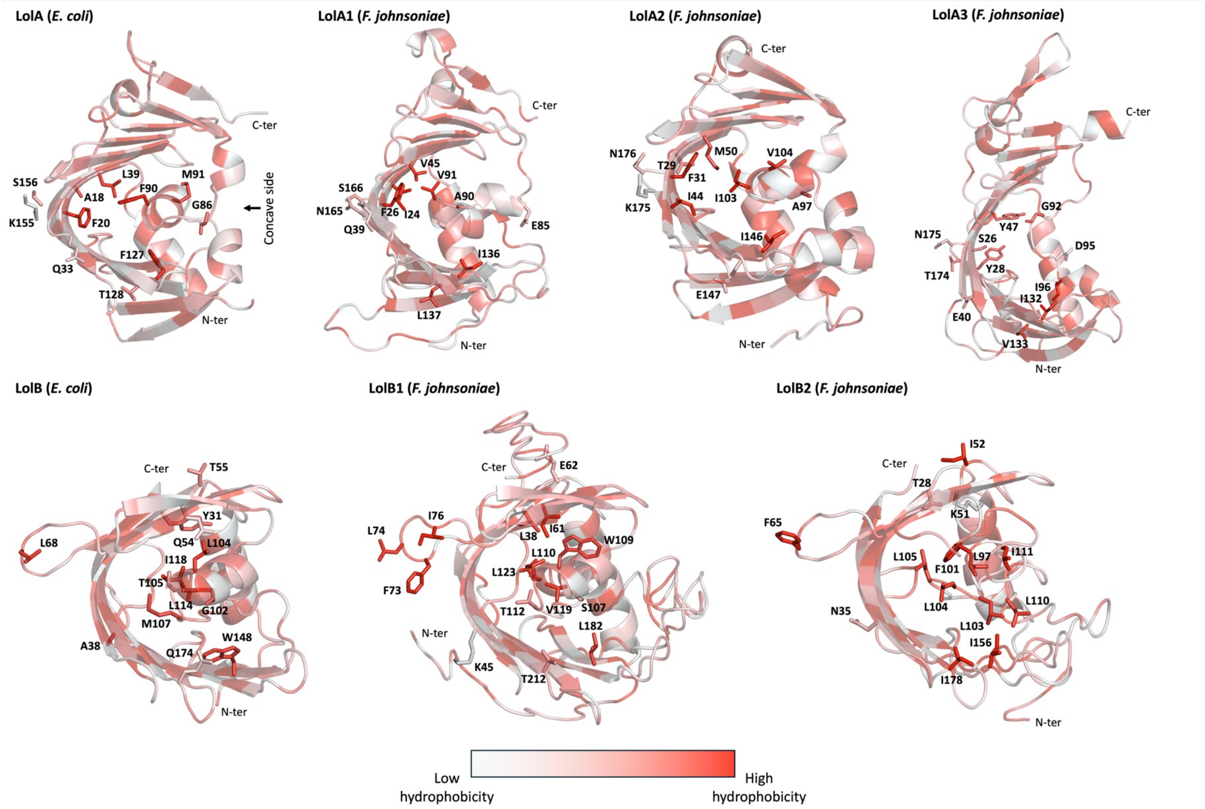

**Supplementary Figure S1**. **Comparison of per-residue hydrophobicity between *E. coli* and *F. johnsoniae* LolA and LolB homologs.** Hydrophobicity scores, ranging from hydrophilic (white) to highly hydrophobic (red), were computed using the Eisenberg's scale and projected on the crystallized LolA (PDB entry: 1UA8) and LolB (PDB entry: 1IWM) structures, as well as LolA1, LolA2, LolA3, LolB1, and LolB2 models represented as cartoon from the barrel opening view. On each structure, the N- and C-terminal positions are indicated. Identified amino acids relevant to lipoprotein binding in *E. coli* and their equivalents in *F. johnsoniae* are shown as stick with their associated labelled position. Residues in the upward loop of LolB proteins are also highlighted as stick with their associated labelled position.

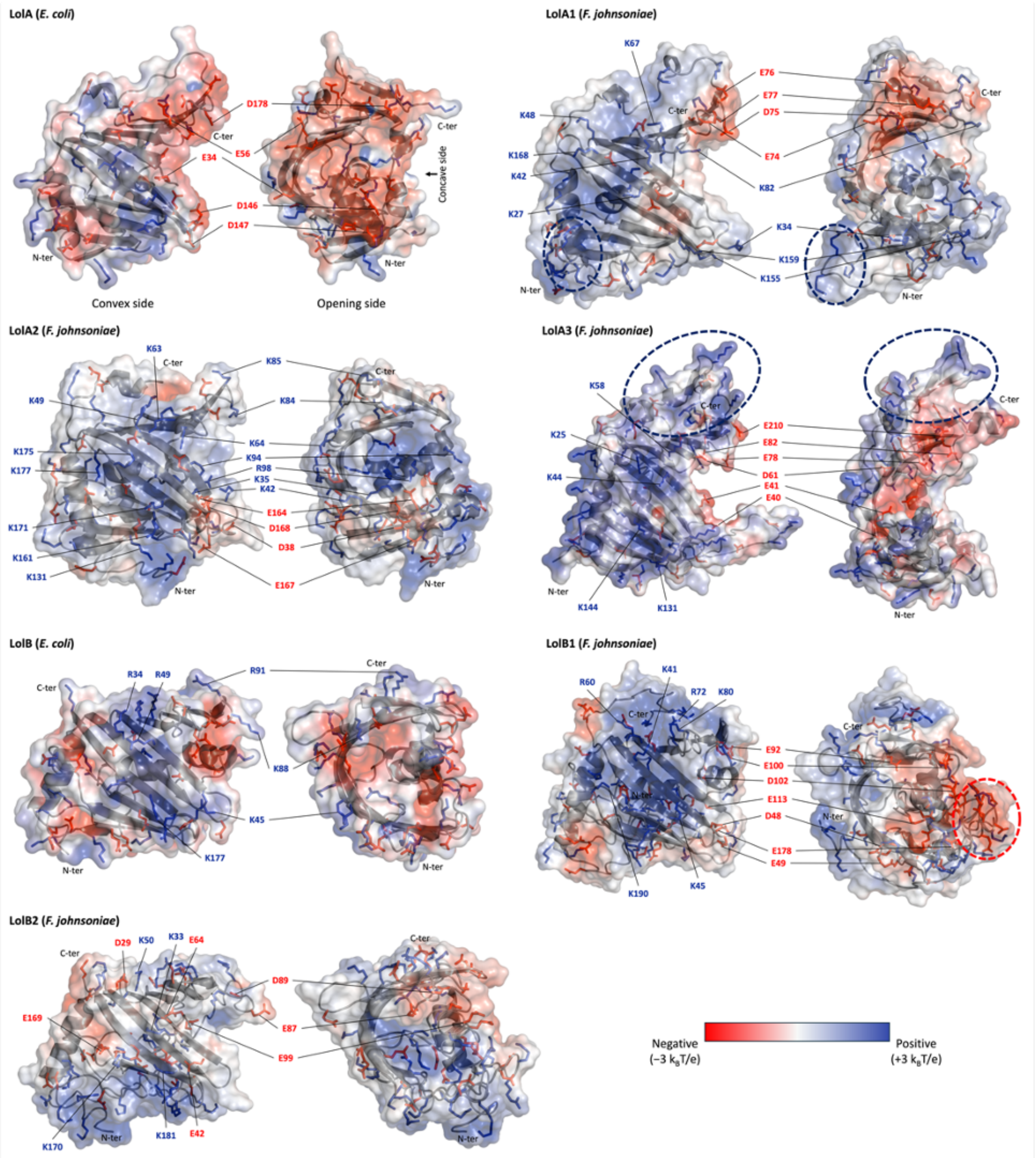

**Supplementary Figure S2**. **Comparison of charge state and distribution between *E. coli* and *F. johnsoniae* LolA and LolB homologs.** Poisson-Boltzmann electrostatic potentials, ranging from −3 $k_BT/e$ (red) to +3 $k_BT/e$ (blue), were mapped on the van der Waals surface of the crystallized LolA (PDB entry: 1UA8) and LolB (PDB entry: 1IWM) structures, as well as LolA1, LolA2, LolA3, LolB1, and LolB2 models represented as grey cartoon. On each structure, the N- and C-terminal positions are indicated. Negatively (aspartate and glutamate) and positively (lysine and arginine) charged residues are shown as red and blue stick, respectively. Relevant charged amino acids with respect to their position on the protein convex side or barrel opening have their position labelled. In LolA1, the positively charged loop extension between α-helix 3 and β-strand 7 is highlighted with a dashed dark blue ellipse. On the LolA3 panel, the positively charged C-terminal region is highlighted with a dashed dark blue ellipse. In LolB1 panel, the negatively charged loop extension between α-helix 3 and β-strand 7 is highlighted with a dashed red ellipse.

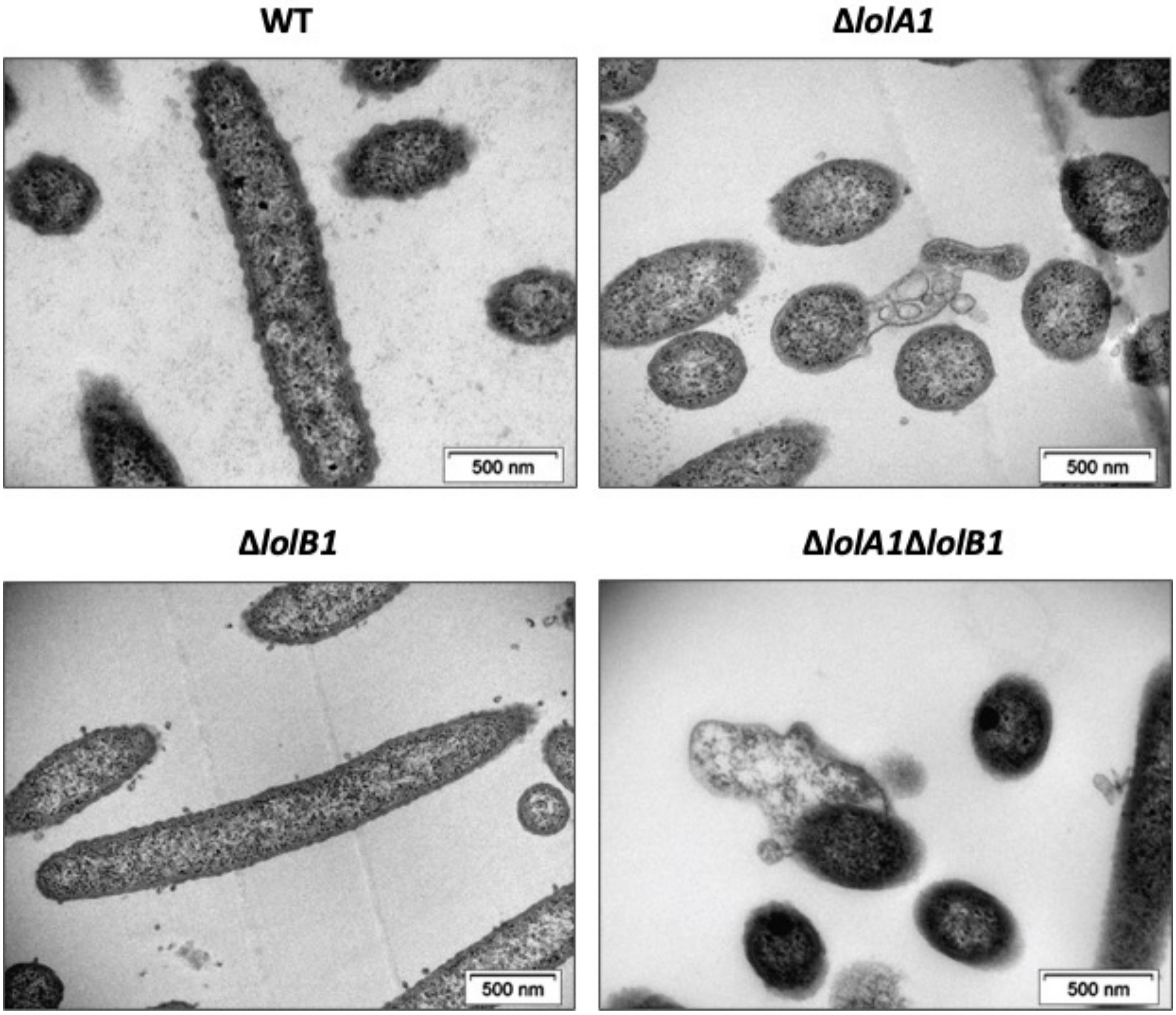

**Supplementary Figure S3.** Transmission Electron Microscopy micrographs of bacteria grown for 16 hours in CYE liquid medium.

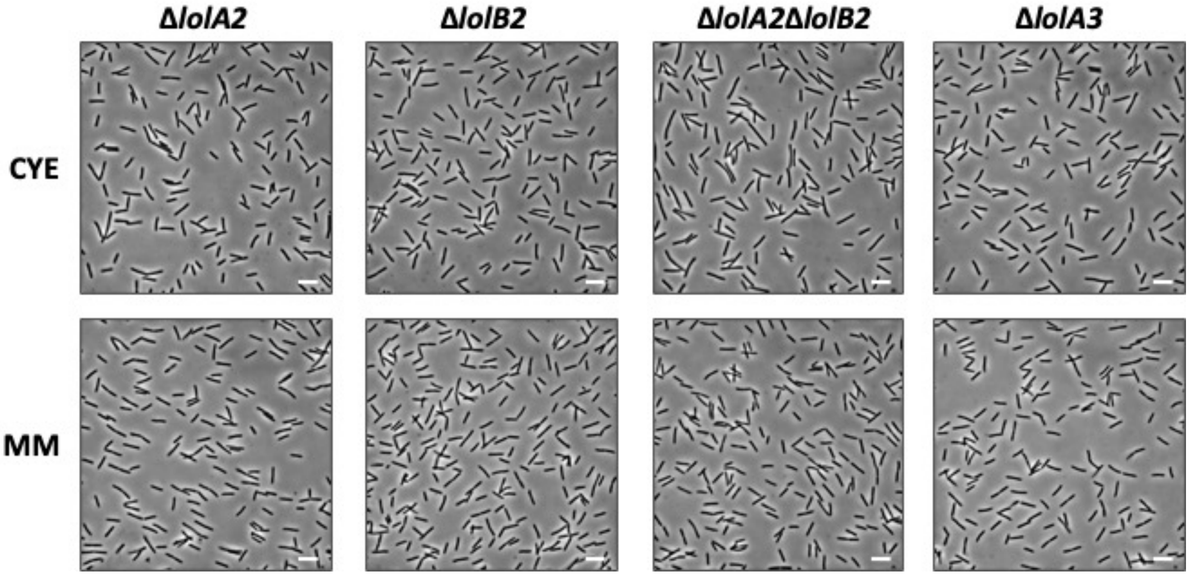

**Supplementary Figure S4.** Bright field microscopy images of bacteria gown for 16 hours in CYE and MM liquid media (bar = 5 µm).

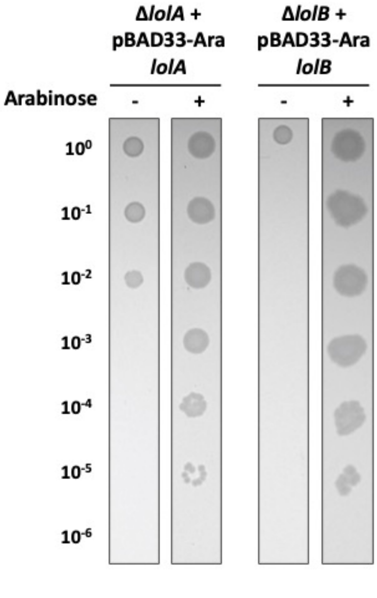

**Supplementary Figure S5 – Deletion of *lolA* or *lolB* in *E. coli MG1655* complemented with *lolA* or *lolB*.** Serial dilution spots on LB agar plates supplemented with (+) or without (-) 0.2% arabinose of the *lolA* and *lolB* mutant strains expressing *E. coli lolA* or *lolB* in trans.

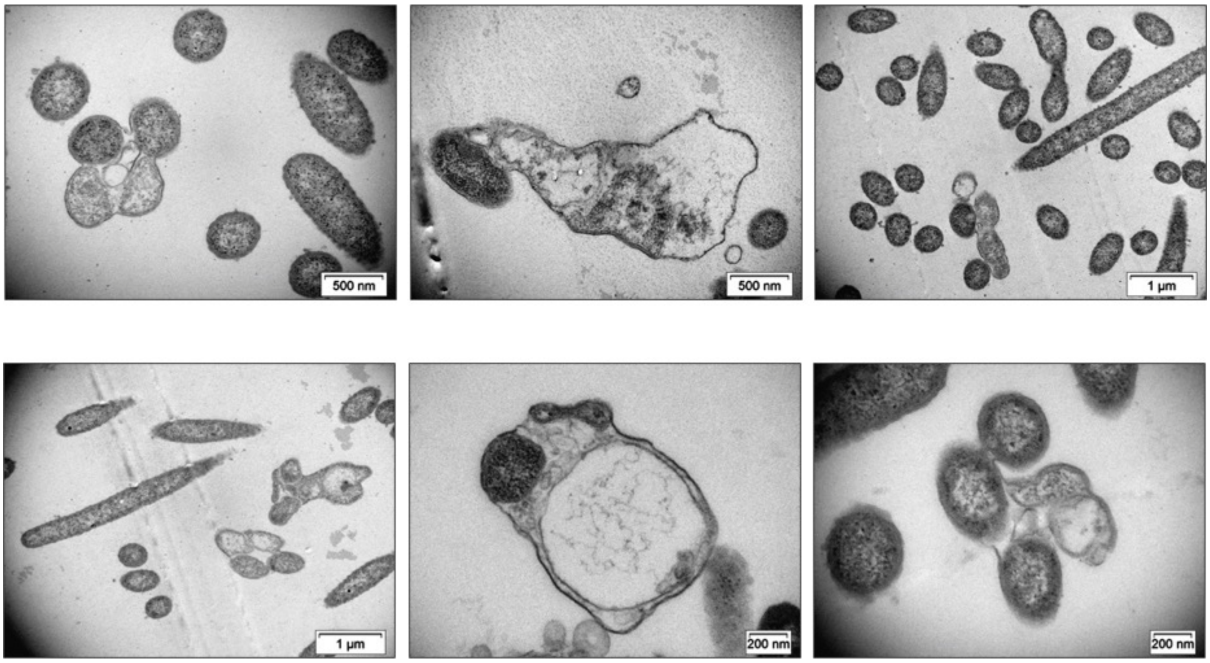

**Supplementary Figure S6.** Transmission Electron Microscopy micrographs of *lolA1* mutant bacteria grown for 16 hours in CYE liquid medium.

**Supplementary Table S1. Composition in hydrophobic, aromatic, negatively and positively charged residues of the Lol protein homologs from *E. coli* (LolA and LolB) and *F. johnsoniae* (LolA1, LolA2, LolA3, LolB1, and LolB2).**

| | Residue percentage (%) | | | | | | | | | |
|---|---|---|---|---|---|---|---|---|---|---|
| | Hydrophobic | Aromatic | Ile/Leu | Val | Asp | Glu | Lys | Arg | − | + |
| **LolA** | 34.6 | 10.4 | 8.8 | 6.6 | 9.3 | 2.2 | 6.6 | 3.3 | 11.5 | 9.9 |
| **LolA1** | 35.6 | 9.8 | 13.4 | 8.2 | 6.7 | 4.1 | 14.4 | 1.0 | 10.8 | 15.4 |
| **LolA2** | 40.5 | 10.5 | 15.8 | 7.4 | 7.4 | 6.3 | 15.8 | 1.0 | 13.7 | 16.8 |
| **LolA3** | 39.7 | 14.2 | 14.6 | 7.3 | 4.6 | 5.9 | 13.2 | 1.8 | 10.5 | 15.0 |
| **LolB** | 34.4 | 9.7 | 12.9 | 4.3 | 7.5 | 3.2 | 6.4 | 5.4 | 10.7 | 11.8 |
| **LolB1** | 39.5 | 10.3 | 16.9 | 5.8 | 4.9 | 8.6 | 11.5 | 2.1 | 13.6 | 13.6 |
| **LolB2** | 37.4 | 12.1 | 18.4 | 4.2 | 6.3 | 8.4 | 14.2 | 2.1 | 14.7 | 16.3 |

**Supplementary Table S2. Proteins with a SPI/SPII/SPII-LES detected in the OM of the *lolA1* and *lolB1* mutants (FC ≥ 1.5, significance ≥ 20).**
Stars (*) indicate proteins for which only a PUL prediction by CAZy was available.
Class descriptions are from the EggNOG database 5.0, except for Polysaccharide utilization (which groups all Sus-like proteins and proteins predicted by EggNOG to be involved in carbohydrate transport and metabolism) and Gliding/T9SS, which are custom classes.

#### Δ*lolA1*

| Accession | Signal peptide | Significance | *lolA1*/WT FC | Description | Gene code | COG Class Description | Localization | CAZy family | PUL (CAZy literature and prediction) | Essentiality |
|---|---|---|---|---|---|---|---|---|---|---|
| A5FIT9 | SPII | 63.91 | 64 | Lipopolysaccharide assembly protein | Fjoh_1847 | Function unknown | OM lipoprotein | | | Yes |
| A5FMM3 | SPII | 101.76 | 64 | Glycine zipper family protein | Fjoh_0505 | Function unknown | OM lipoprotein | | | |
| A5FAM8 | SPI | 41.45 | 64 | SprF-like protein | Fjoh_4749 | Gliding/T9SS | OM integral protein | | | |
| A5FHF7 | SPII | 53.13 | 51.03 | Hypothetical lipoprotein | Fjoh_2342 | Function unknown | OM lipoprotein | | | |
| A5FNR2 | SPI | 20.18 | 23.99 | OMP_b-brl_2 domain-containing protein | Fjoh_0127 | Cell wall/membrane/envelope biogenesis | OM integral protein | | | |
| A5FAI4 | SPII | 69.72 | 20.35 | DUF3300 domain-containing protein | Fjoh_4779 | RNA processing and modification | OM lipoprotein | | | |
| A5FAC1 | SPII-LES | 55.28 | 15.71 | Lipocalin-like domain-containing protein | Fjoh_4843 | Lipid transport and metabolism | Surface-exposed lipoprotein | | | |
| A5FNT5 | SPII | 45.14 | 14.21 | Hypothetical lipoprotein | Fjoh_0097 | Function unknown | OM lipoprotein | | | |
| A5FP10 | SPI | 44.7 | 12.72 | Peptidase, subfamily S1B unassigned peptidases | Fjoh_0019 | Posttranslational modification, protein turnover, chaperones | Periplasmic | | | |
| A5FFN3 | SPI | 58.28 | 11.6 | Uncharacterized protein | Fjoh_2959 | Function unknown | Periplasmic | | | |
| A5FDE8 | SPI | 36.7 | 7.76 | DUF3570 domain-containing protein | Fjoh_3759 | Lipid transport and metabolism | OM integral protein | | | |
| A5FMH4 | SPII-LES | 43.3 | 5.38 | Hypothetical lipoprotein | Fjoh_0561 | Function unknown | Surface-exposed lipoprotein | | | |
| A5FBR6 | SPII | 29.15 | 5.11 | DUF4369 domain-containing protein | Fjoh_4343 | Energy production and conversion | OM lipoprotein | | | |
| A5FHZ5 | SPII | 37.78 | 4.51 | Hypothetical lipoprotein | Fjoh_2151 | Function unknown | OM lipoprotein | | | |
| A5FBX5 | SPII | 50.92 | 4.22 | Efflux transporter, RND family, MFP subunit | Fjoh_4294 | Cell wall/membrane/envelope biogenesis | IM lipoprotein | | | |
| A5FM15 | SPI | 50.7 | 4.18 | BatE-like protein | Fjoh_0723 | Signal transduction mechanisms | Periplasmic | | | Low fitness |
| A5F9V9 | SPII-LES | 50.24 | 4.14 | Cytochrome-c peroxidase | Fjoh_5007 | Energy production and conversion | Surface-exposed lipoprotein | | | Low fitness |
| A5FJP1 | SPII | 42.8 | 3.83 | Heavy metal transport/detoxification protein | Fjoh_1541 | Inorganic ion transport and metabolism | OM lipoprotein | | | Low fitness |
| A5FK44 | SPII-LES | 45.57 | 3.81 | Hypothetical lipoprotein | Fjoh_1393 | Function unknown | Surface-exposed lipoprotein | | | |

| | | | | | | | | |
|---|---|---|---|---|---|---|---|---|
| A5FK25 | SPII | 63.26 | 3.76 | PKD domain containing protein | Fjoh_1410 | Signal transduction mechanisms | OM lipoprotein | |
| A5FM13 | SPI | 34.37 | 3.6 | BatC-like protein | Fjoh_0721 | Coenzyme transport and metabolism | Periplasmic | Yes |
| A5FMJ3 | SPII-LES | 27.98 | 3.56 | Hypothetical lipoprotein | Fjoh_0546 | Gliding/T9SS | Surface-exposed lipoprotein | |
| A5FB40 | SPI | 28.82 | 3.3 | Outer membrane protein beta-barrel domain-containing protein | Fjoh_4578 | Cell wall/membrane/envelope biogenesis | OM integral protein | |
| A5FNB1 | SPII | 41.06 | 3.27 | Peptide-methionine (R)-S-oxide reductase | Fjoh_0270 | Posttranslational modification, protein turnover, chaperones | OM lipoprotein | |
| A5FCD4 | SPII | 26.92 | 3.15 | Efflux transporter, RND family, MFP subunit | Fjoh_4132 | Cell wall/membrane/envelope biogenesis | IM lipoprotein | |
| A5FHB6 | SPI | 23.72 | 3.12 | Peptidoglycan-binding LysM | Fjoh_2379 | Cell wall/membrane/envelope biogenesis | Periplasmic | |
| A5FEP9 | SPII | 32.13 | 3.1 | Glucose sorbosone dehydrogenase | Fjoh_3307 | Carbohydrate transport and metabolism | OM lipoprotein | |
| A5FF07 | SPI | 23.89 | 2.72 | Gliding motility protein RemI | Fjoh_3194 | Gliding/T9SS | Cell surface/extracellular | |
| A5FA29 | SPI | 39.82 | 2.68 | Membrane protein involved in aromatic hydrocarbon degradation | Fjoh_4941 | Lipid transport and metabolism | OM integral protein | |
| A5F9X2 | SPII | 21.3 | 2.64 | DUF6565 domain-containing protein | Fjoh_5000 | Function unknown | OM lipoprotein | |
| A5FIS7 | SPI | 29.78 | 2.38 | Exopolyphosphatase-like protein | Fjoh_1868 | Nucleotide transport and metabolism | Periplasmic | |
| A5FBC4 | SPI | 26.05 | 2.37 | DUF541 domain-containing protein | Fjoh_4501 | Function unknown | Periplasmic | |
| Q8KRP0 | SPII | 39.28 | 2.31 | GldH | Fjoh_0890 | Gliding/T9SS | OM lipoprotein | |
| A5FHK8 | SPI | 20.38 | 2.26 | Peptidase family M1 possible nickel uptake system protein | Fjoh_2281 | Inorganic ion transport and metabolism | Periplasmic | |
| A5FEI9 | SPII | 22.23 | 2.25 | Predicted outer membrane protein | Fjoh_3371 | Function unknown | OM lipoprotein | |
| A5FM48 | SPI | 45.11 | 2.19 | Peptidase family S33 | Fjoh_0689 | Lipid transport and metabolism | Periplasmic | |
| A5FL26 | SPI | 41.91 | 2.06 | Peptidase subfamily M23B-like protein | Fjoh_1067 | Cell cycle control, cell division, chromosome partitioning | Periplasmic | |
| A5FK94 | SPII | 24.36 | 2.03 | Hypothetical lipoprotein | Fjoh_1339 | Function unknown | OM lipoprotein | |
| A5FHZ7 | SPI | 26.93 | 2 | Cytochrome c biogenesis protein, transmembrane region | Fjoh_2139 | Energy production and conversion | Periplasmic | |
| A5FJV6 | SPI | 24.7 | 1.96 | Uncharacterized protein | Fjoh_1486 | Function unknown | Periplasmic | |
| A5FJ14 | SPII | 24.36 | 1.94 | Hypothetical lipoprotein | Fjoh_1777 | Function unknown | OM lipoprotein | |
| A5FHJ6 | SPII | 35.17 | 1.94 | Hypothetical lipoprotein | Fjoh_2300 | Function unknown | OM lipoprotein | |
| A5FHQ3 | SPI | 35.82 | 1.93 | Tetratricopeptide TPR_2 repeat protein | Fjoh_2246 | Function unknown | Periplasmic | |
| A5FLC8 | SPI | 36.74 | 1.92 | PgPepO oligopeptidase peptidase family M13 | Fjoh_0959 | Posttranslational modification, protein turnover, chaperones | Periplasmic | |

| | | | | | | | | | |
|---|---|---|---|---|---|---|---|---|---|
| A5FJF0 | SPI | 22.64 | 1.89 | Sporulation domain protein | Fjoh_1633 | Function unknown | Periplasmic | | |
| A5FMN9 | SPI | 23.48 | 1.84 | TPR repeat-containing protein | Fjoh_0488 | Function unknown | Periplasmic | | |
| A5F9R8 | SPII | 20.61 | 1.83 | Hypothetical lipoprotein | Fjoh_5055 | Function unknown | OM lipoprotein | | |
| A5FHZ0 | SPI | 23.1 | 1.8 | Putative esterase | Fjoh_2146 | Function unknown | Periplasmic | | |
| A5FNV3 | SPII | 27.8 | 1.79 | Periplasmic binding protein | Fjoh_0083 | Inorganic ion transport and metabolism | OM lipoprotein | | |
| A5FHC0 | SPII | 29.65 | 1.78 | Peptidylprolyl isomerase | Fjoh_2368 | Posttranslational modification, protein turnover, chaperones | OM lipoprotein | | |
| A5FGY9 | SPII | 36.08 | 1.77 | Hypothetical lipoprotein | Fjoh_2510 | Function unknown | OM lipoprotein | | |
| A5FKP3 | SPII-LES | 22.64 | 1.71 | Hypothetical lipoprotein | Fjoh_1201 | Function unknown | Surface-exposed lipoprotein | | |
| A5F9Z8 | SPII | 28.45 | 1.67 | Lipocalin-like domain-containing protein | Fjoh_4978 | Lipid transport and metabolism | OM lipoprotein | | |
| A5FMX9 | SPI | 30.21 | 1.65 | Uncharacterized protein | Fjoh_0411 | Function unknown | Periplasmic | | |
| A5FIE5 | SPII-LES | 26.32 | 1.6 | Beta-xylanase | Fjoh_1996 | Carbohydrate transport and metabolism | Surface-exposed lipoprotein | GH10 | |
| A5FNC0 | SPII | 23.08 | 1.51 | DUF4369 domain-containing protein | Fjoh_0264 | Function unknown | OM lipoprotein | | |
| A5FKB8 | SPI | 27.96 | 0.66 | SusC-like TonB-dependent receptor | Fjoh_1314 | Carbohydrate transport and metabolism | OM integral protein | SusC | |
| A5FAG3 | SPI | 20.7 | 0.65 | Uncharacterized conserved protein UCP016719 | Fjoh_4806 | Function unknown | Periplasmic | | |
| A5FFP8 | SPI | 31.86 | 0.63 | Glutathione peroxidase | Fjoh_2944 | Posttranslational modification, protein turnover, chaperones | Periplasmic | | |
| A5FGA4 | SPI | 23.14 | 0.62 | Adhesin domain-containing protein | Fjoh_2750 | Cell wall/membrane/envelope biogenesis | Cell surface/extracellular | | |
| A5FE42 | SPI | 23.85 | 0.61 | Peptidase family M28 | Fjoh_3518 | Function unknown | Periplasmic | | |
| A5FIR7 | SPII-LES | 29.73 | 0.6 | DNA-binding beta-propeller fold protein YncE | Fjoh_1874 | Nucleotide transport and metabolism | Surface-exposed lipoprotein | | |
| A5FNF7 | SPI | 20.66 | 0.59 | Peptidase subfamily M48-like protein | Fjoh_0225 | Cell wall/membrane/envelope biogenesis | Periplasmic | | |
| A5FLE9 | SPI | 39.17 | 0.59 | TonB-dependent receptor, plug | Fjoh_0928 | Inorganic ion transport and metabolism | OM integral protein | | |
| A5FJZ8 | SPI | 24.06 | 0.59 | TonB-dependent receptor, plug | Fjoh_1438 | Inorganic ion transport and metabolism | OM integral protein | | |
| A5FBC3 | SPI | 25.03 | 0.59 | SusC-like TonB-dependent receptor | Fjoh_4500 | Carbohydrate transport and metabolism | OM integral protein | SusC | PUL34* |
| A5FG07 | SPII | 20.56 | 0.59 | Lipocalin-like domain-containing protein | Fjoh_2832 | Lipid transport and metabolism | OM lipoprotein | | |
| A5FCM6 | SPI | 23.82 | 0.58 | AB hydrolase-1 domain-containing protein | Fjoh_4027 | Function unknown | Periplasmic | | |

| | | | | | | | | |
|---|---|---|---|---|---|---|---|---|
| A5FIZ8 | SPI | 44.7 | 0.58 | Outer membrane protein beta-barrel domain-containing protein | Fjoh_1789 | Cell wall/membrane/envelope biogenesis | OM integral protein | | |
| A5FK08 | SPII | 22.79 | 0.58 | DUF4249 domain-containing protein | Fjoh_1435 | Function unknown | OM lipoprotein | | |
| A5FBY8 | SPI | 21.34 | 0.57 | Isochorismatase hydrolase | Fjoh_4273 | Secondary metabolites biosynthesis, transport and catabolism | Periplasmic | | |
| A5FB67 | SPI | 38.98 | 0.55 | SusC-like TonB-dependent receptor | Fjoh_4559 | Carbohydrate transport and metabolism | OM integral protein | SusC | PUL30/34 & PUL35 |
| A5FHU7 | SPII | 33 | 0.55 | Cytochrome C | Fjoh_2192 | Energy production and conversion | OM lipoprotein | | |
| A5FKE6 | SPI | 21.43 | 0.54 | Response regulator receiver protein | Fjoh_1291 | Transcription | Periplasmic | | |
| A5FAE4 | SPII | 29.09 | 0.54 | Candidate beta-glycosidase Glycoside hydrolase family 3 | Fjoh_4819 | Carbohydrate transport and metabolism | OM lipoprotein | Pept_SE/GH 3 | PUL32 & PUL38 |
| A5FIT1 | SPI | 25.61 | 0.53 | GldN | Fjoh_1856 | Gliding/T9SS | Periplasmic | | |
| A5FJJ6 | SPI | 22.99 | 0.53 | TonB-dependent receptor | Fjoh_1588 | Inorganic ion transport and metabolism | OM integral protein | | |
| A5FJI1 | SPII-LES | 27.18 | 0.53 | Hypothetical lipoprotein | Fjoh_1606 | Function unknown | Surface-exposed lipoprotein | | |
| A5FD24 | SPII-LES | 20.07 | 0.53 | RagB/SusD domain protein | Fjoh_3881 | Carbohydrate transport and metabolism | Surface-exposed lipoprotein | SusD | PUL20 & PUL23 |
| A5FBC2 | SPII-LES | 24.38 | 0.53 | RagB/SusD domain protein | Fjoh_4499 | Carbohydrate transport and metabolism | Surface-exposed lipoprotein | SusD | PUL34 |
| A5FFW4 | SPII-LES | 31.84 | 0.52 | DUF4302 domain-containing protein | Fjoh_2891 | Function unknown | Surface-exposed lipoprotein | | |
| A5FFU9 | SPII-LES | 28.33 | 0.52 | RagB/SusD domain protein | Fjoh_2893 | Carbohydrate transport and metabolism | Surface-exposed lipoprotein | SusD | PUL11 & PUL15 |
| A5FJY9 | SPI | 51.64 | 0.5 | Integral membrane sensor signal transduction histidine kinase | Fjoh_1442 | Signal transduction mechanisms | IM lipoprotein | | |
| A5FEB7 | SPI | 24.91 | 0.5 | Outer membrane protein beta-barrel comain-containing protein | Fjoh_3442 | Cell wall/membrane/envelope biogenesis | OM integral protein | | |
| A5FAF7 | SPI | 25.13 | 0.5 | Uncharacterized conserved protein UCP016719 | Fjoh_4816 | Function unknown | Periplasmic | | |
| A5FCW3 | SPII-LES | 31.7 | 0.5 | RagB/SusD domain protein | Fjoh_3944 | Carbohydrate transport and metabolism | Surface-exposed lipoprotein | SusD | PUL21 & PUL24 |
| A5FA28 | SPII | 21.25 | 0.5 | Hypothetical lipoprotein | Fjoh_4940 | Function unknown | OM lipoprotein | | |
| A5FMI9 | SPI | 34.89 | 0.49 | Phosphate-selective porin O and P | Fjoh_0542 | Inorganic ion transport and metabolism | OM integral protein | | |
| A5FK31 | SPI | 29.4 | 0.48 | SusC-like TonB-dependent receptor | Fjoh_1405 | Carbohydrate transport and metabolism | OM integral protein | SusC | PUL3 & PUL6 |
| A5FA64 | SPI | 26.38 | 0.48 | Outer membrane efflux protein | Fjoh_4902 | Cell wall/membrane/envelope biogenesis | OM integral protein | | |
| A5FJ27 | SPI | 50.04 | 0.48 | Uncharacterized protein | Fjoh_1765 | Coenzyme transport and metabolism | OM integral protein | | |

| | | | | | | | | | | |
|---|---|---|---|---|---|---|---|---|---|---|
| A5FM74 | SPII-LES | 30.32 | 0.48 | RagB/SusD domain protein | Fjoh_0666 | Carbohydrate transport and metabolism | Surface-exposed lipoprotein | SusD | PUL3* | |
| A5FFU8 | SPII | 24.76 | 0.48 | Substrate import-associated zinc metallohydrolase lipoprotein | Fjoh_2892 | Function unknown | OM lipoprotein | | | |
| A5FIS8 | SPII | 30.84 | 0.48 | GldK | Fjoh_1853 | Gliding/T9SS | OM lipoprotein | | | |
| A5FCU8 | SPI | 26.65 | 0.47 | Outer membrane protein-like protein | Fjoh_3959 | Cell wall/membrane/envelope biogenesis | OM integral protein | | | |
| A5FEH0 | SPI | 27.12 | 0.47 | Candidate beta-glycosidase Glycoside hydrolase family 3 | Fjoh_3389 | Carbohydrate transport and metabolism | Periplasmic | GH3 | | |
| A5FJM5 | SPI | 32.5 | 0.47 | Candidate beta-glucosidase Glycoside hydrolase family 3 | Fjoh_1567 | Carbohydrate transport and metabolism | Periplasmic | GH3 | PUL4 & PUL7 | |
| A5FF76 | SPII-LES | 26.62 | 0.47 | RagB/SusD domain protein | Fjoh_3126 | Carbohydrate transport and metabolism | Surface-exposed lipoprotein | SusD | PUL13 & PUL17 | |
| A5FMK2 | SPI | 24.27 | 0.46 | Colicin import membrane protein | Fjoh_0540 | Function unknown | Unknown | | | Yes |
| A5FLZ8 | SPI | 26.72 | 0.46 | SusC-like TonB-dependent receptor | Fjoh_0736 | Carbohydrate transport and metabolism | OM integral protein | SusC | PUL3 & PUL6 | |
| A5FKI3 | SPI | 30.74 | 0.45 | Alpha-ketoglutarate decarboxylase | Fjoh_1257 | Energy production and conversion | OM integral protein | | | |
| A5FGE4 | SPI | 22.76 | 0.45 | SusC-like TonB-dependent receptor | Fjoh_2711 | Carbohydrate transport and metabolism | OM integral protein | SusC | PUL10 & PUL14 | |
| A5FFV0 | SPI | 33.93 | 0.45 | SusC-like TonB-dependent receptor | Fjoh_2894 | Carbohydrate transport and metabolism | OM integral protein | SusC | PUL11 & PUL15 | |
| A5FB12 | SPI | 20.72 | 0.45 | Alpha/beta hydrolase fold protein | Fjoh_4606 | Function unknown | Periplasmic | | | |
| A5FAG6 | SPI | 27.03 | 0.45 | GHL10 domain-containing protein | Fjoh_4809 | Function unknown | Periplasmic | | | |
| A5FE81 | SPII | 35.85 | 0.45 | BamD | Fjoh_3469 | Cell wall/membrane/envelope biogenesis | OM lipoprotein | | | Yes |
| A5FN99 | SPI | 32.45 | 0.44 | Peptidase family M24 | Fjoh_0276 | Amino acid transport and metabolism | Periplasmic | | | |
| A5FNF3 | SPI | 26 | 0.44 | BamB-like | Fjoh_0237 | Cell wall/membrane/envelope biogenesis | Periplasmic | | | |
| A5FA84 | SPII | 22.79 | 0.44 | Lipocalin-like domain-containing protein | Fjoh_4891 | Lipid transport and metabolism | OM lipoprotein | | | |
| A5FLQ6 | SPI | 27.65 | 0.43 | TonB-dependent receptor | Fjoh_0821 | Inorganic ion transport and metabolism | OM integral protein | | | |
| A5FKR4 | SPI | 35.63 | 0.43 | DUF3078 domain-containing protein | Fjoh_1173 | Cell wall/membrane/envelope biogenesis | OM integral protein | | | |
| A5FIL8 | SPI | 33.74 | 0.43 | SusC-like TonB-dependent receptor | Fjoh_1924 | Carbohydrate transport and metabolism | OM integral protein | SusC | PUL8* | |
| A5FJW3 | SPI | 24.6 | 0.42 | Carboxypeptidase-like regulatory domain-containing protein | Fjoh_1476 | Cell wall/membrane/envelope biogenesis | OM integral protein | | | |
| A5FJV5 | SPI | 33.28 | 0.42 | TonB-dependent receptor | Fjoh_1485 | Inorganic ion transport and metabolism | OM integral protein | | | |
| A5FH73 | SPI | 40.76 | 0.42 | FGE-sulfatase domain-containing protein | Fjoh_2417 | Function unknown | Periplasmic | | | |

| | | | | | | | | |
|---|---|---|---|---|---|---|---|---|
| A5FFB9 | SPI | 30.29 | 0.42 | TonB-dependent receptor, plug | Fjoh_3092 | Inorganic ion transport and metabolism | OM integral protein | | |
| A5FAG7 | SPI | 20.26 | 0.42 | Candidate esterase Carbohydrate esterase family 1 | Fjoh_4810 | Function unknown | Periplasmic | | |
| A5FA54 | SPI | 34.14 | 0.42 | TonB-dependent receptor, plug | Fjoh_4916 | Coenzyme transport and metabolism | OM integral protein | | |
| A5FAG5 | SPI | 21.68 | 0.42 | Beta-N-acetylhexosaminidase | Fjoh_4808 | Carbohydrate transport and metabolism | Periplasmic | GH20 | |
| A5FIN2 | SPII | 51.34 | 0.42 | DUF4136 domain-containing protein | Fjoh_1907 | Function unknown | OM lipoprotein | | |
| A5FL78 | SPI | 28.46 | 0.41 | Uncharacterized protein | Fjoh_1011 | Function unknown | Periplasmic | | |
| A5FJX0 | SPI | 29.85 | 0.41 | SprT | Fjoh_1466 | Gliding/T9SS | OM integral protein | | |
| A5FH92 | SPI | 24.79 | 0.41 | Candidate d-4,5 unsaturated beta-glycuronidase Glycoside hydrolase family 88 | Fjoh_2406 | Carbohydrate transport and metabolism | Periplasmic | GH88 | |
| A5FGB5 | SPI | 29.5 | 0.41 | DUF4294 domain-containing protein | Fjoh_2737 | Function unknown | Periplasmic | | |
| A5FE38 | SPI | 28.27 | 0.41 | Outermembrane protein | Fjoh_3514 | Function unknown | OM integral protein | | |
| A5FB56 | SPI | 36.89 | 0.41 | SusC-like TonB-dependent receptor | Fjoh_4562 | Carbohydrate transport and metabolism | OM integral protein | SusC | PUL30/34 & PUL35 |
| A5FK37 | SPI | 31.25 | 0.41 | Candidate alpha-glucosidase Glycoside hydrolase family 97 | Fjoh_1400 | Carbohydrate transport and metabolism | Periplasmic | GH97 | PUL3 & CAZyme cluster 1 |
| A5FC59 | SPII-LES | 30.29 | 0.41 | RagB/SusD domain protein | Fjoh_4195 | Carbohydrate transport and metabolism | Surface-exposed lipoprotein | SusD | PUL24 & PUL27 |
| A5FK70 | SPI | 25.11 | 0.4 | TonB-dependent receptor | Fjoh_1368 | Inorganic ion transport and metabolism | OM integral protein | | |
| A5FIC9 | SPI | 31.5 | 0.4 | Peptidase family M16 domain protein | Fjoh_2012 | Function unknown | Periplasmic | | |
| A5FFP7 | SPI | 34.19 | 0.4 | Hypothetical protein | Fjoh_2958 | Cell wall/membrane/envelope biogenesis | OM integral protein | | |
| A5FHC7 | SPII | 27.8 | 0.4 | Peptidase family M61 domain protein | Fjoh_2360 | Function unknown | OM lipoprotein | | |
| A5FA21 | SPII-LES | 35.63 | 0.4 | RagB/SusD domain protein | Fjoh_4950 | Carbohydrate transport and metabolism | Surface-exposed lipoprotein | SusD | PUL33 & PUL39 |
| A5FBB1 | SPI | 28.06 | 0.39 | Glutathione hydrolase proenzyme | Fjoh_4506 | Amino acid transport and metabolism | Periplasmic | | |
| A5FAK8 | SPI | 20.51 | 0.39 | OmpH family outer membrane protein | Fjoh_4758 | Function unknown | Periplasmic | | |
| A5FAV4 | SPI | 28.9 | 0.39 | SusC-like TonB-dependent receptor | Fjoh_4671 | Carbohydrate transport and metabolism | OM integral protein | SusC | PUL31 & PUL37 |
| A5FMB4 | SPI | 28.64 | 0.39 | GspD-like type II secretion system secretin protein | Fjoh_0618 | Intracellular trafficking, secretion, and vesicular transport | Periplasmic | | |
| A5FJ92 | SPI | 52.11 | 0.38 | DUF6089 domain-containing protein | Fjoh_1692 | Cell wall/membrane/envelope biogenesis | OM integral protein | | |

| | | | | | | | | | | |
|---|---|---|---|---|---|---|---|---|---|---|
| A5FGD1 | SPII-LES | 21.91 | 0.38 | RagB/SusD domain protein | Fjoh_2712 | Carbohydrate transport and metabolism | Surface-exposed lipoprotein | SusD | PUL10 & PUL14 | |
| A5FLA7 | SPI | 28.54 | 0.37 | SprF | Fjoh_0978 | Gliding/T9SS | OM integral protein | | | |
| A5FF77 | SPI | 36.61 | 0.37 | SusC-like TonB-dependent receptor | Fjoh_3127 | Carbohydrate transport and metabolism | OM integral protein | SusC | PUL13 & PUL17 | |
| A5FDH2 | SPI | 27.26 | 0.37 | Alpha/beta hydrolase fold protein | Fjoh_3736 | Lipid transport and metabolism | Periplasmic | | | |
| A5FM73 | SPI | 31.93 | 0.36 | SusC-like TonB-dependent receptor | Fjoh_0665 | Carbohydrate transport and metabolism | OM integral protein | SusC | PUL3 | |
| A5FH56 | SPI | 28.11 | 0.36 | SusC-like TonB-dependent receptor | Fjoh_2431 | Carbohydrate transport and metabolism | OM integral protein | SusC | PUL9 & PUL13 | |
| A5FHX2 | SPII | 27.38 | 0.36 | Peptidase family S33 | Fjoh_2171 | Lipid transport and metabolism | OM lipoprotein | | | |
| A5F9W0 | SPI | 36.62 | 0.35 | Transporter | Fjoh_5008 | Energy production and conversion | OM integral protein | | | Yes |
| A5FD66 | SPI | 63.62 | 0.35 | Carboxypeptidase regulatory-like domain-containing protein | Fjoh_3841 | Inorganic ion transport and metabolism | OM integral protein | | | |
| A5FK32 | SPII-LES | 27.43 | 0.35 | RagB/SusD domain protein | Fjoh_1406 | Carbohydrate transport and metabolism | Surface-exposed lipoprotein | SusD | PUL3 & PUL6 | |
| A5FJ03 | SPI | 50.9 | 0.34 | Porin | Fjoh_1779 | Function unknown | OM integral protein | | | |
| A5FBI5 | SPI | 47.7 | 0.34 | SusC-like TonB-dependent receptor | Fjoh_4434 | Carbohydrate transport and metabolism | OM integral protein | SusC | PUL29 & PUL32 | |
| A5FA22 | SPI | 33.27 | 0.34 | SusC-like TonB-dependent receptor | Fjoh_4951 | Carbohydrate transport and metabolism | OM integral protein | SusC | PUL33 & PUL39 | |
| A5FK33 | SPII | 48.26 | 0.34 | Hypothetical lipoprotein | Fjoh_1407 | Function unknown | OM lipoprotein | | | |
| A5FA20 | SPII-LES | 31.14 | 0.34 | IPT/TIG domain-containing protein | Fjoh_4949 | Function unknown | Surface-exposed lipoprotein | | | |
| A5FBS1 | SPI | 32.06 | 0.33 | DUF6377 domain-containing protein | Fjoh_4332 | Transcription & Signal transduction mechanisms | Periplasmic | | | |
| A5FA04 | SPI | 26.05 | 0.33 | NLP/P60 protein | Fjoh_4966 | Cell wall/membrane/envelope biogenesis | Periplasmic | | | |
| A5FC08 | SPI | 32.03 | 0.33 | SusC-like TonB-dependent receptor | Fjoh_4255 | Carbohydrate transport and metabolism | OM integral protein | SusC | PUL26 & PUL29 | |
| A5FFR0 | SPI | 29.63 | 0.33 | Outer membrane efflux protein | Fjoh_2941 | Intracellular trafficking, secretion, and vesicular transport | OM integral protein | | | |
| A5FMK1 | SPI | 54.79 | 0.32 | Uncharacterized protein | Fjoh_0539 | Function unknown | OM integral protein | | | |
| A5FNW0 | SPI | 34.21 | 0.32 | Endonuclease/exonuclease/phosphatase | Fjoh_0074 | Function unknown | Periplasmic | | | |
| A5FB55 | SPII-LES | 40.73 | 0.32 | RagB/SusD domain protein | Fjoh_4561 | Carbohydrate transport and metabolism | Surface-exposed lipoprotein | SusD | PUL30/PUL34 & PUL35 | |
| A5FAV5 | SPII-LES | 24.4 | 0.32 | RagB/SusD domain protein | Fjoh_4672 | Carbohydrate transport and metabolism | Surface-exposed lipoprotein | SusD | PUL31 & PUL37 | |
| A5FNK1 | SPI | 39.66 | 0.31 | SusC-like TonB-dependent receptor | Fjoh_0185 | Carbohydrate transport and metabolism | OM integral protein | SusC | PUL1* | |

| | | | | | | | | | |
|---|---|---|---|---|---|---|---|---|---|
| A5FKB7 | SPI | 21.35 | 0.31 | L, D-transpeptidase catalytic domain-containing protein | Fjoh_1328 | Cell wall/membrane/envelope biogenesis | Periplasmic | | |
| A5FD40 | SPI | 26.48 | 0.29 | SusC-like TonB-dependent receptor | Fjoh_3871 | Carbohydrate transport and metabolism | OM integral protein | SusC | PUL20 & PUL23 |
| A5FKG3 | SPI | 39.12 | 0.29 | PKD domain-containing protein | Fjoh_1272 | Function unknown | OM integral protein | | |
| A5FM71 | SPII | 33.14 | 0.29 | Ig-like domain-containing protein | Fjoh_0663 | Function unknown | OM lipoprotein | | |
| A5FE85 | SPI | 42.3 | 0.28 | Metallophosphoesterase | Fjoh_3473 | Nucleotide transport and metabolism | Periplasmic | | |
| A5FEI0 | SPI | 41.36 | 0.27 | Outer membrane protein beta-barrel domain-containing protein | Fjoh_3381 | Function unknown | OM integral protein | | |
| A5FH32 | SPI | 39.53 | 0.27 | TonB-dependent receptor | Fjoh_2466 | Inorganic ion transport and metabolism | OM integral protein | | |
| A5FMB7 | SPII-LES | 31.79 | 0.27 | Fibronectin, type III domain protein | Fjoh_0621 | Function unknown | Surface-exposed lipoprotein | | |
| A5FN23 | SPII | 52.97 | 0.27 | Wza | Fjoh_0361 | Cell wall/membrane/envelope biogenesis | OM lipoprotein | | |
| A5FF33 | SPI | 27.23 | 0.26 | TonB-dependent receptor, plug | Fjoh_3179 | Coenzyme transport and metabolism | OM integral protein | | |
| A5FC34 | SPI | 52.24 | 0.26 | TonB-dependent siderophore receptor | Fjoh_4221 | Inorganic ion transport and metabolism | OM integral protein | | |
| A5FLI1 | SPII | 30.9 | 0.26 | Efflux transporter, RND family, MFP subunit | Fjoh_0906 | Cell wall/membrane/envelope biogenesis | IM lipoprotein | | |
| A5FE84 | SPII | 41.66 | 0.26 | 5'-Nucleotidase domain protein | Fjoh_3472 | Nucleotide transport and metabolism | OM lipoprotein | | |
| A5FGA3 | SPI | 50.97 | 0.25 | Putative auto-transporter adhesin head GIN domain-containing protein | Fjoh_2749 | Cell wall/membrane/envelope biogenesis | Cell surface/extracellular | | |
| A5FJM0 | SPI | 30.16 | 0.24 | Candidate beta-glycosidase Glycoside hydrolase family 30 | Fjoh_1562 | Carbohydrate transport and metabolism | Periplasmic | GH30_1 | PUL4 & PUL7 |
| A5FCM0 | SPI | 38.29 | 0.24 | TonB-dependent siderophore receptor | Fjoh_4039 | Inorganic ion transport and metabolism | OM integral protein | | |
| A5FJM2 | SPI | 41.81 | 0.24 | Candidate beta-glycosidase Glycoside hydrolase family 3 | Fjoh_1564 | Carbohydrate transport and metabolism | Periplasmic | GH3 | PUL4 & PUL7 |
| A5FNK0 | SPII-LES | 45.09 | 0.24 | RagB/SusD domain protein | Fjoh_0184 | Carbohydrate transport and metabolism | Surface-exposed lipoprotein | | PUL1 |
| A5FAV9 | SPII | 51.84 | 0.24 | Peptidase family M28 | Fjoh_4661 | Function unknown | OM lipoprotein | | |
| A5FEN2 | SPI | 21.62 | 0.23 | TonB-dependent siderophore receptor | Fjoh_3320 | Inorganic ion transport and metabolism | OM integral protein | | |
| A5FNJ9 | SPII-LES | 51.1 | 0.23 | Fibronectin, type III domain protein | Fjoh_0183 | Function unknown | Surface-exposed lipoprotein | | |
| A5FNC9 | SPII | 62.84 | 0.23 | Lipocalin-like domain-containing protein | Fjoh_0259 | Lipid transport and metabolism | OM lipoprotein | | |
| A5FND5 | SPI | 30.58 | 0.22 | TonB-dependent receptor | Fjoh_0249 | Inorganic ion transport and metabolism | OM integral protein | | |
| A5FEE5 | SPI | 33.44 | 0.22 | Outer membrane protein beta-barrel domain-containing protein | Fjoh_3401 | Function unknown | OM integral protein | | |

| | | | | | | | | | | |
|---|---|---|---|---|---|---|---|---|---|---|
| A5FC74 | SPI | 49.35 | 0.22 | SusC-like TonB-dependent receptor | Fjoh_4194 | Carbohydrate transport and metabolism | OM integral protein | SusC | PUL24 & PUL27 | |
| A5FJN2 | SPI | 55.26 | 0.22 | SusC-like TonB-dependent receptor | Fjoh_1560 | Carbohydrate transport and metabolism | OM integral protein | SusC | PUL4 & PUL7 | |
| A5FKQ1 | SPII | 64.18 | 0.22 | Hypothetical lipoprotein | Fjoh_1177 | Posttranslational modification, protein turnover, chaperones | OM lipoprotein | | | |
| A5FJM4 | SPII | 38.55 | 0.22 | Candidate beta-glycosidase Glycoside hydrolase family 30 | Fjoh_1566 | Carbohydrate transport and metabolism | OM lipoprotein | GH30_3 | PUL4 & PUL7 | |
| A5FB06 | SPII | 41.3 | 0.21 | META domain-containing protein | Fjoh_4623 | Posttranslational modification, protein turnover, chaperones | OM lipoprotein | | | |
| A5FJM1 | SPII | 34.41 | 0.2 | Candidate beta-glycosidase Glycoside hydrolase family 30 | Fjoh_1563 | Carbohydrate transport and metabolism | Periplasmic | GH30_1 | PUL4 & PUL7 | |
| A5FGL7 | SPII | 27.25 | 0.2 | 3-keto-disaccharide hydrolase domain-containing protein | Fjoh_2621 | Function unknown | OM lipoprotein | | | |
| A5FE35 | SPII-LES | 38.24 | 0.19 | RagB/SusD domain protein | Fjoh_3524 | Carbohydrate transport and metabolism | Surface-exposed lipoprotein | SusD | PUL16 & PUL20 | |
| A5FIM2 | SPII | 80.22 | 0.18 | Peptidoglycan hydrolase | Fjoh_1913 | Cell wall/membrane/envelope biogenesis | OM lipoprotein | | | |
| A5FJ15 | SPI | 25.49 | 0.17 | Lipid/polyisoprenoid-binding YceI-like domain-containing protein | Fjoh_1778 | Function unknown | Periplasmic | | | |
| A5FEK0 | SPI | 42.73 | 0.17 | Outer membrane protein beta-barrel domain-containing protein | Fjoh_3350 | Cell wall/membrane/envelope biogenesis | OM integral protein | | | |
| A5FEL3 | SPI | 34.89 | 0.17 | Uncharacterized protein | Fjoh_3349 | Function unknown | OM integral protein | | | |
| A5FCV6 | SPI | 49.23 | 0.16 | SprF-like protein | Fjoh_3951 | Gliding/T9SS | OM integral protein | | | |
| A5FNS6 | SPI | 52.43 | 0.15 | Phosphate-selective porin O and P | Fjoh_0105 | Function unknown | OM integral protein | | | Yes |
| A5FE72 | SPI | 35.59 | 0.15 | SprF-like protein | Fjoh_3477 | Gliding/T9SS | OM integral protein | | | |
| A5FE36 | SPI | 53.19 | 0.13 | SusC-like TonB-dependent receptor | Fjoh_3525 | Carbohydrate transport and metabolism | OM integral protein | SusC | PUL16 & PUL20 | |
| A5FMS1 | SPII | 47.68 | 0.13 | Peptidase family M28 | Fjoh_0454 | Function unknown | OM lipoprotein | | | |
| A5FJZ6 | SPII | 55.44 | 0.13 | Hypothetical lipoprotein | Fjoh_1436 | Function unknown | OM lipoprotein | | | |
| A5FBD6 | SPI | 43.97 | 0.1 | Outer membrane efflux protein | Fjoh_4485 | Intracellular trafficking, secretion, and vesicular transport | OM integral protein | | | |
| A5FI22 | SPI | 76.55 | 0.1 | Possible lipoprotein carrier protein LolA | Fjoh_2111 | Cell wall/membrane/envelope biogenesis | Periplasmic | | | |
| A5FNC8 | SPII | 62.58 | 0.1 | OmpA/MotB domain protein | Fjoh_0258 | Cell wall/membrane/envelope biogenesis | OM lipoprotein | | | |
| A5FAL0 | SPII | 60.41 | 0.1 | META domain-containing protein | Fjoh_4760 | Posttranslational modification, protein turnover, chaperones | OM lipoprotein | | | |
| A5FHI0 | SPI | 54.95 | 0.07 | OmpA/MotB domain protein | Fjoh_2321 | Cell wall/membrane/envelope biogenesis | OM integral protein | | | |
| A1E5U4 | SPI | 28.23 | 0.05 | SprD | Fjoh_0980 | Gliding/T9SS | Unknown | | | |

| | | | | | | | | | |
|---|---|---|---|---|---|---|---|---|---|
| A5FJM9 | SPII | 51.16 | 0.05 | GldJ | Fjoh_1557 | Gliding/T9SS | OM lipoprotein | | |
| A5FFC2 | SPI | 47.4 | 0.02 | DUF2911 domain-containing protein | Fjoh_3076 | Function unknown | Periplasmic | | |
| A5FK70 | SPI | 52.17 | 0 | TonB-dependent receptor | Fjoh_1368 | Inorganic ion transport and metabolism | OM integral protein | | |
| A5FG24 | SPI | 135.31 | 0 | Choloylglycine hydrolase peptidase family C59 | Fjoh_2831 | Cell wall/membrane/envelope biogenesis | Periplasmic | | |
| A5FBC7 | SPII-LES | 117.53 | 0 | RagB/SusD domain protein | Fjoh_4490 | Polysaccharide utilization | Surface-exposed lipoprotein | SusD | PUL33 |

### *ΔlolB1*

| Accession | Signal peptide | Significance | *lolB1*/WT FC | Description | Gene code | COG Class Description | Localization | CAZy family | PUL (literature and CAZy prediction) | Essentiality |
|---|---|---|---|---|---|---|---|---|---|---|
| A5FC16 | SPII-LES | 115.07 | 64 | Peptidase family M57 | Fjoh_4241 | Posttranslational modification, protein turnover, chaperones | OM lipoprotein | | | |
| A5FHF7 | SPII | 54.84 | 48.29 | Hypothetical lipoprotein | Fjoh_2342 | Function unknown | OM lipoprotein | | | |
| A5FNT4 | SPI | 70.41 | 29.07 | DUF4252 domain-containing protein | Fjoh_0096 | Function unknown | Periplasmic | | | |
| A5FNT5 | SPII | 49.59 | 18.55 | Hypothetical lipoprotein | Fjoh_0097 | Function unknown | OM lipoprotein | | | |
| A5FDE8 | SPI | 46.23 | 8.22 | DUF3570 domain-containing protein | Fjoh_3759 | Lipid transport and metabolism | OM integral protein | | | |
| A5FF13 | SPII-LES | 35.45 | 6.36 | Hypothetical lipoprotein | Fjoh_3186 | Function unknown | Surface-exposed lipoprotein | | | |
| A5FAC1 | SPII-LES | 48.44 | 5.83 | Lipocalin-like domain-containing protein | Fjoh_4843 | Lipid transport and metabolism | Surface-exposed lipoprotein | | | |
| A5FHZ5 | SPII | 72.66 | 5.67 | Hypothetical lipoprotein | Fjoh_2151 | Function unknown | OM lipoprotein | | | |
| A5FNR2 | SPI | 55.74 | 5.27 | OMP_b-brl_2 domain-containing protein | Fjoh_0127 | Cell wall/membrane/envelope biogenesis | OM integral protein | | | |
| A5F9V9 | SPII-LES | 47.43 | 3.53 | Cytochrome-c peroxidase | Fjoh_5007 | Energy production and conversion | Surface-exposed lipoprotein | | | |
| A5F9R8 | SPII | 38.34 | 3.2 | Hypothetical lipoprotein | Fjoh_5055 | Function unknown | OM lipoprotein | | | |
| A5FHA6 | SPI | 43.83 | 3.17 | Antitoxin component YwqK of YwqJK toxin-antitoxin module | Fjoh_2385 | Function unknown | Periplasmic | | | |
| A5FNB6 | SPII | 27.56 | 3.02 | META domain-containing protein | Fjoh_0275 | Posttranslational modification, protein turnover, chaperones | OM lipoprotein | | | |
| A5FND7 | SPI | 36.13 | 2.9 | TonB-dependent receptor, plug | Fjoh_0252 | Inorganic ion transport and metabolism | OM integral protein | | | |
| A5FAK9 | SPI | 35.49 | 2.9 | DUF4251 domain-containing protein | Fjoh_4759 | Function unknown | Periplasmic | | | |
| A5FB40 | SPI | 34.29 | 2.74 | Outer membrane protein beta-barrel domain-containing protein | Fjoh_4578 | Cell wall/membrane/envelope biogenesis | OM integral protein | | | |
| A5FJ27 | SPI | 44.08 | 2.47 | Uncharacterized protein | Fjoh_1765 | Coenzyme transport and metabolism | OM integral protein | | | |

| | | | | | | | | | |
|---|---|---|---|---|---|---|---|---|---|
| A5FAG3 | SPI | 32.39 | 2.36 | Uncharacterized conserved protein UCP016719 | Fjoh_4806 | Function unknown | Periplasmic | | |
| A5FNX9 | SPII | 31.59 | 2.26 | Peptidylprolyl isomerase | Fjoh_0050 | Posttranslational modification, protein turnover, chaperones | OM lipoprotein | | |
| A5FLD6 | SPII | 26.42 | 2.2 | DUF4421 domain-containing protein | Fjoh_0952 | Cell motility | OM lipoprotein | SusD | PUL26 & PUL29 |
| A5FC07 | SPII-LES | 24.35 | 2.2 | RagB/SusD domain protein | Fjoh_4254 | Carbohydrate transport and metabolism | Surface-exposed lipoprotein | SusC | PUL26 & PUL29 |
| A5FC08 | SPI | 23.28 | 2.16 | SusC-like TonB-dependent receptor | Fjoh_4255 | Carbohydrate transport and metabolism | OM integral protein | | |
| A5FAF7 | SPI | 32.54 | 2.15 | Uncharacterized conserved protein UCP016719 | Fjoh_4816 | Function unknown | Periplasmic | | |
| A5FC53 | SPII | 27.88 | 2.13 | Copper homeostasis protein | Fjoh_4206 | Cell wall/membrane/envelope biogenesis | OM lipoprotein | | |
| A5FEP9 | SPII-LES | 23.07 | 2.13 | Glucose sorbosone dehydrogenase | Fjoh_3307 | Carbohydrate transport and metabolism | Surface-exposed lipoprotein | | |
| A5FAG0 | SPII-LES | 22.68 | 2.09 | Hypothetical lipoprotein | Fjoh_4803 | Function unknown | Surface-exposed lipoprotein | SusD | PUL22 & PUL25 |
| A5FCG9 | SPII-LES | 36.91 | 2.08 | RagB/SusD domain protein | Fjoh_4094 | Carbohydrate transport and metabolism | Surface-exposed lipoprotein | | |
| A5FMH4 | SPII-LES | 22.25 | 2.06 | Hypothetical lipoprotein | Fjoh_0561 | Function unknown | Surface-exposed lipoprotein | | |
| A5FNS8 | SPI | 34.61 | 2.05 | PepSY_like domain-containing protein | Fjoh_0107 | Function unknown | Periplasmic | | |
| A5FI22 | SPI | 70.99 | 2.04 | Possible lipoprotein carrier protein LolA | Fjoh_2111 | Cell wall/membrane/envelope biogenesis | Periplasmic | | |
| A5FLY5 | SPII-LES | 21.6 | 2.04 | PKD domain containing protein | Fjoh_0757 | Function unknown | Surface-exposed lipoprotein | | |
| A5FAI4 | SPII | 32.55 | 2.03 | DUF3300 domain-containing protein | Fjoh_4779 | RNA processing and modification | OM lipoprotein | | |
| A5FMN5 | SPI | 30.98 | 1.99 | Beta-lactamase peptidase family S12 | Fjoh_0500 | Defense mechanisms | Periplasmic | | |
| A5FGA3 | SPI | 55.98 | 1.98 | Putative auto-transporter adhesin head GIN domain-containing protein | Fjoh_2749 | Cell wall/membrane/envelope biogenesis | Cell surface/extracellular | | |
| A5FEZ9 | SPI | 20.4 | 1.93 | Uncharacterized protein | Fjoh_3206 | Function unknown | Periplasmic | | |
| A5FMK7 | SPII | 20.62 | 1.91 | Hypothetical lipoprotein | Fjoh_0527 | Function unknown | OM lipoprotein | GH105 | PUL26 & PUL29 |
| A5FC14 | SPII | 27.09 | 1.9 | Candidate d-4,5-unsaturated beta-glycuronidase Glycoside hydrolase family 105 | Fjoh_4250 | Carbohydrate transport and metabolism | OM lipoprotein | SusC | PUL6 & PUL10 |
| A5FIC3 | SPI | 29.73 | 1.89 | SusC-like TonB-dependent receptor | Fjoh_2020 | Carbohydrate transport and metabolism | OM integral protein | SusC | PUL20 & PUL23 |
| A5FD25 | SPI | 23.16 | 1.86 | SusC-like TonB-dependent receptor | Fjoh_3882 | Carbohydrate transport and metabolism | OM integral protein | | |

| | | | | | | | | | |
|---|---|---|---|---|---|---|---|---|---|
| A5FHB6 | SPI | 40.82 | 1.86 | Peptidoglycan-binding LysM | Fjoh_2379 | Cell wall/membrane/envelope biogenesis | Periplasmic | | |
| A5FJB8 | SPI | 27.43 | 1.85 | SprF-like protein | Fjoh_1677 | Gliding/T9SS | OM integral protein | | |
| A5FHQ3 | SPI | 29.74 | 1.85 | Tetratricopeptide TPR_2 repeat protein | Fjoh_2246 | Function unknown | Periplasmic | | |
| A5FK94 | SPII-LES | 22 | 1.81 | Hypothetical lipoprotein | Fjoh_1339 | Function unknown | Surface-exposed lipoprotein | | |
| A5FJV3 | SPI | 24.15 | 1.8 | Glutamine cyclotransferase | Fjoh_1483 | Function unknown | Periplasmic | | |
| A5FGG2 | SPII | 23.3 | 1.8 | Hypothetical lipoprotein | Fjoh_2679 | Function unknown | OM lipoprotein | | |
| A5FL98 | SPI | 21.32 | 1.78 | Uncharacterized protein | Fjoh_0984 | Function unknown | Periplasmic | | |
| A5FEL3 | SPI | 30.21 | 1.78 | Uncharacterized protein | Fjoh_3349 | Function unknown | OM integral protein | | |
| A5FMN9 | SPI | 30.21 | 1.77 | TPR repeat-containing protein | Fjoh_0488 | Function unknown | Periplasmic | | |
| A5FLC8 | SPI | 37.81 | 1.76 | PgPepO oligopeptidase peptidase family M13 | Fjoh_0959 | Posttranslational modification, protein turnover, chaperones | Periplasmic | | |
| A5FFP7 | SPI | 23.3 | 1.76 | Hypothetical protein | Fjoh_2958 | Cell wall/membrane/envelope biogenesis | OM integral protein | | |
| A5FD35 | SPII | 22.37 | 1.76 | Lipolytic enzyme, G-D-S-L family | Fjoh_3879 | Amino acid transport and metabolism | OM lipoprotein | | |
| A5FF07 | SPI | 23.81 | 1.66 | Gliding motility protein RemI | Fjoh_3194 | Gliding/T9SS | Cell surface/extracellular | | |
| A5FKG3 | SPI | 25.81 | 1.66 | PKD domain-containing protein | Fjoh_1272 | Function unknown | OM integral protein | | |
| A5FH76 | SPII | 20.43 | 1.66 | Hypothetical lipoprotein | Fjoh_2420 | Function unknown | OM lipoprotein | | |
| A5FNT1 | SPII-LES | 37.49 | 1.65 | Peptidase S41, subfamily S41A unassigned peptidases | Fjoh_0093 | Cell wall/membrane/envelope biogenesis | Surface-exposed lipoprotein | Pept_SE/GH 3 | PUL32 & PUL38 |
| A5FAE4 | SPII | 22.82 | 1.64 | Candidate beta-glycosidase Glycoside hydrolase family 3 | Fjoh_4819 | Carbohydrate transport and metabolism | OM lipoprotein | | |
| A5FH32 | SPI | 24.45 | 1.59 | TonB-dependent receptor | Fjoh_2466 | Inorganic ion transport and metabolism | OM integral protein | | |
| A5FMW8 | SPI | 20.97 | 1.57 | Dipeptidyl-peptidase | Fjoh_0416 | Amino acid transport and metabolism | Periplasmic | | |
| A5FJV4 | SPI | 21.37 | 1.55 | GLPGLI family protein | Fjoh_1484 | Function unknown | Unknown | | |
| A5FF84 | SPI | 28.19 | 1.54 | DUF4861 domain-containing protein | Fjoh_3122 | Function unknown | Periplasmic | | |
| A5FC67 | SPI | 29.12 | 1.52 | DUF4861 domain-containing protein | Fjoh_4187 | Function unknown | Periplasmic | | |
| A5FJM8 | SPI | 33.5 | 1.51 | Por secretion system protein PorU precursor. C-terminal signal peptidase | Fjoh_1556 | Gliding/T9SS | OM protein | | |
| A5FGM2 | SPI | 22.13 | 1.51 | Peptidase family S33-like protein | Fjoh_2626 | Lipid transport and metabolism | Periplasmic | GH20 | |
| A5FAG5 | SPI | 21.13 | 1.5 | Beta-N-acetylhexosaminidase | Fjoh_4808 | Carbohydrate transport and metabolism | Periplasmic | | |

| | | | | | | | | | |
|---|---|---|---|---|---|---|---|---|---|
| A5FIZ8 | SPI | 21.74 | 0.66 | Outer membrane protein beta-barrel domain-containing protein | Fjoh_1789 | Cell wall/membrane/envelope biogenesis | OM integral protein | | |
| A5FIR6 | SPII-LES | 31.11 | 0.66 | PKD domain containing protein | Fjoh_1873 | Function unknown | Surface-exposed lipoprotein | | |
| A5FCU8 | SPI | 28.83 | 0.64 | Outer membrane protein-like protein | Fjoh_3959 | Cell wall/membrane/envelope biogenesis | OM integral protein | | |
| A5FJ10 | SPI | 24.1 | 0.63 | Aminopeptidase N-like protein | Fjoh_1773 | Amino acid transport and metabolism | Periplasmic | | |
| A5FHK9 | SPI | 35.82 | 0.62 | TonB-dependent siderophore receptor | Fjoh_2282 | Inorganic ion transport and metabolism | OM integral protein | | |
| A5FJJ6 | SPI | 23.09 | 0.6 | TonB-dependent receptor | Fjoh_1588 | Inorganic ion transport and metabolism | OM integral protein | | |
| A5FCM6 | SPI | 21.43 | 0.59 | AB hydrolase-1 domain-containing protein | Fjoh_4027 | Function unknown | Periplasmic | | |
| A5FBD6 | SPI | 22.02 | 0.58 | Outer membrane efflux protein | Fjoh_4485 | Intracellular trafficking, secretion, and vesicular transport | OM integral protein | | |
| A5FNF3 | SPI | 26.25 | 0.57 | BamB-like | Fjoh_0237 | Cell wall/membrane/envelope biogenesis | Periplasmic | | |
| A5FL26 | SPI | 36.94 | 0.55 | Peptidase subfamily M23B-like protein | Fjoh_1067 | Cell cycle control, cell division, chromosome partitioning | Periplasmic | | |
| A5FD66 | SPI | 24.61 | 0.55 | Carboxypeptidase regulatory-like domain-containing protein | Fjoh_3841 | Inorganic ion transport and metabolism | OM integral protein | GH3 | PUL4 & PUL7 |
| A5FJM5 | SPI | 37.69 | 0.53 | Candidate beta-glucosidase Glycoside hydrolase family 3 | Fjoh_1567 | Carbohydrate transport and metabolism | Periplasmic | SusD | PUL34 |
| A5FBC2 | SPII-LES | 25.79 | 0.51 | RagB/SusD domain protein | Fjoh_4499 | Carbohydrate transport and metabolism | Surface-exposed lipoprotein | GH97 | PUL3 & CAZyme cluster 1 |
| A5FK37 | SPI | 34.98 | 0.48 | Candidate alpha-glucosidase Glycoside hydrolase family 97 | Fjoh_1400 | Carbohydrate transport and metabolism | Periplasmic | | |
| A5FN31 | SPI | 24.15 | 0.44 | Polysaccharide export protein | Fjoh_0353 | Cell wall/membrane/envelope biogenesis | Periplasmic | SusC | PUL31 & PUL37 |
| A5FAV4 | SPI | 30.31 | 0.44 | SusC-like TonB-dependent receptor | Fjoh_4671 | Carbohydrate transport and metabolism | OM integral protein | SusD | PUL10 & PUL14 |
| A5FGD1 | SPII-LES | 26.22 | 0.41 | RagB/SusD domain protein | Fjoh_2712 | Carbohydrate transport and metabolism | Surface-exposed lipoprotein | | |
| A5FNV5 | SPII-LES | 36.62 | 0.39 | PrcB C-terminal domain-containing protein | Fjoh_0069 | Function unknown | Surface-exposed lipoprotein | GH3 | |
| A5FEH0 | SPI | 34.7 | 0.34 | Candidate beta-glycosidase Glycoside hydrolase family 3 | Fjoh_3389 | Carbohydrate transport and metabolism | Periplasmic | | |
| A5FIT1 | SPI | 52.46 | 0.33 | GldN | Fjoh_1856 | Gliding/T9SS | Periplasmic | | |
| A1E5U5 | SPI | 47.7 | 0.32 | SprB | Fjoh_0979 | Gliding/T9SS | Surface-exposed | | |
| A5FHG9 | SPII | 45.05 | 0.32 | DUF4197 domain-containing protein | Fjoh_2327 | Function unknown | OM lipoprotein | | |
| A5FJZ6 | SPII | 54.55 | 0.31 | Hypothetical lipoprotein | Fjoh_1436 | Function unknown | OM lipoprotein | | |

| | | | | | | | | | |
|---|---|---|---|---|---|---|---|---|---|
| A5FMB7 | SPII-LES | 45.03 | 0.29 | Fibronectin, type III domain protein | Fjoh_0621 | Function unknown | Surface-exposed lipoprotein | | |
| A5FMC4 | SPI | 64.4 | 0.28 | Uncharacterized protein | Fjoh_0610 | Function unknown | Periplasmic | | |
| A5FIM2 | SPII | 58.81 | 0.28 | Peptidoglycan hydrolase | Fjoh_1913 | Cell wall/membrane/envelope biogenesis | OM lipoprotein | | |
| A5FJZ7 | SPII | 32.54 | 0.27 | DUF4249 domain-containing protein | Fjoh_1437 | Function unknown | OM lipoprotein | GH30_1 | PUL4 & PUL7 |
| A5FJM1 | SPII | 32.72 | 0.26 | Candidate beta-glycosidase Glycoside hydrolase family 30 | Fjoh_1563 | Carbohydrate transport and metabolism | Periplasmic | | |
| A5FMB4 | SPI | 57.53 | 0.24 | GspD-like type II secretion system secretin protein | Fjoh_0618 | Intracellular trafficking, secretion, and vesicular transport | Periplasmic | | |
| A5FFR0 | SPI | 50.84 | 0.24 | Outer membrane efflux protein | Fjoh_2941 | Intracellular trafficking, secretion, and vesicular transport | OM integral protein | | |
| A5FN23 | SPII | 62.52 | 0.23 | Wza | Fjoh_0361 | Cell wall/membrane/envelope biogenesis | OM lipoprotein | | |
| A5FJM9 | SPII | 51.32 | 0.22 | GldJ | Fjoh_1557 | Gliding/T9SS | OM lipoprotein | SusD | PUL31 & PUL37 |
| A5FAV5 | SPII-LES | 42.6 | 0.21 | RagB/SusD domain protein | Fjoh_4672 | Carbohydrate transport and metabolism | Surface-exposed lipoprotein | SusC | PUL4 & PUL7 |
| A5FJN2 | SPI | 64.36 | 0.2 | SusC-like TonB-dependent receptor | Fjoh_1560 | Carbohydrate transport and metabolism | OM integral protein | | |
| A5FHI9 | SPI | 40.57 | 0.19 | DUF5723 domain-containing protein | Fjoh_2310 | Cell wall/membrane/envelope biogenesis | OM integral protein | SusD | PUL4 & PUL7 |
| A5FJL9 | SPII-LES | 66.23 | 0.18 | RagB/SusD domain protein | Fjoh_1561 | Carbohydrate transport and metabolism | Surface-exposed lipoprotein | | |
| A5FJ38 | SPI | 44.7 | 0.17 | Outer membrane protein beta-barrel domain-containing protein | Fjoh_1745 | Cell wall/membrane/envelope biogenesis | OM integral protein | | |
| A5FCV6 | SPI | 47.24 | 0.16 | SprF-like protein | Fjoh_3951 | Gliding/T9SS | OM integral protein | | |
| A5FNW0 | SPI | 61.25 | 0.15 | Endonuclease/exonuclease/phosphatase | Fjoh_0074 | Function unknown | Periplasmic | GH3 | PUL4 & PUL7 |
| A5FJM2 | SPI | 47.09 | 0.13 | Candidate beta-glycosidase Glycoside hydrolase family 3 | Fjoh_1564 | Carbohydrate transport and metabolism | Periplasmic | | |
| A5FHI0 | SPI | 58.23 | 0.08 | OmpA/MotB domain protein | Fjoh_2321 | Cell wall/membrane/envelope biogenesis | OM integral protein | | |
| A5FIS8 | SPII | 59.6 | 0.08 | GldK | Fjoh_1853 | Gliding/T9SS | OM lipoprotein | | |
| A5FL25 | SPII | 81.18 | 0.02 | Hypothetical lipoprotein | Fjoh_1066 | Function unknown | OM lipoprotein | | |

**Supplementary Table S3. Gliding and T9S-related proteins detected in the OM of the *lolA1* and *lolB1* mutants.**
Fold change and significance values between brackets are respectively < 1.5 and < 20.

| Description | Signal peptide | Accession number | Gene code | Δ*lolA1*/WT FC | Significance | Δ*lolB1*/WT FC | Significance |
|---|---|---|---|---|---|---|---|
| SprD | SPI | A1E5U4 | *Fjoh_0980* | 0.05 | 28.23 | 0.47 | (17.00) |
| GldJ | SPII | A5FJM9 | *Fjoh_1557* | 0.05 | 51.16 | 0.22 | 51.32 |
| SprE | SPII | A1E5T9 | *Fjoh_1051* | 0.05 | (7.03) | 0.06 | (7.90) |
| SprF-like protein | SPI | A5FE72 | *Fjoh_3477* | 0.15 | 35.59 | 0.48 | (16.91) |
| SprF-like protein | SPI | A5FCV6 | *Fjoh_3951* | 0.16 | 49.23 | 0.16 | 47.24 |
| SprF | SPI | A5FLA7 | *Fjoh_0978* | 0.37 | 28.54 | (0.84) | (1.22) |
| SprT | SPI | A5FJX0 | *Fjoh_1466* | 0.41 | 29.85 | 1.67 | (18.94) |
| GldK | SPII | A5FIS8 | *Fjoh_1853* | 0.48 | 30.84 | 0.08 | 59.60 |
| GldN | SPI | A5FIT1 | *Fjoh_1856* | 0.53 | 25.61 | 0.33 | 52.46 |
| SprB | SPI | A1E5U5 | *Fjoh_0979* | (0.78) | (12.22) | 0.32 | 47.70 |
| PorV | SPI | A5FJM7 | *Fjoh_1555* | (0.80) | (10.08) | (1.35) | 22.91 |
| SprA | SPI | Q5I6C7 | *Fjoh_1653* | (0.81) | (7.74) | (0.80) | (8.12) |
| GldM | SPI | A5FIT0 | *Fjoh_1855* | (0.92) | (3.92) | (0.78) | 26.20 |
| SprF-like protein | SPI | A5FJB8 | *Fjoh_1677* | (0.95) | (0.74) | 1.85 | 27.43 |
| RemH | SPI | A5FL98 | *Fjoh_0984* | (1.01) | (0.71) | 1.78 | 21.32 |
| GldB | SPII | A5FJ02 | *Fjoh_1793* | (1.08) | (1.08) | (0.90) | (9.54) |
| RemF | SPI | A5FEZ9 | *Fjoh_3206* | (1.12) | (0.65) | 1.93 | 20.40 |
| RemG | SPI | A5FL97 | *Fjoh_0983* | (1.22) | (5.03) | (1.23) | (10.18) |
| GldI | SPII | A5FHC1 | *Fjoh_2369* | (1.43) | 30.45 | (1.02) | (0.99) |
| GldH | SPII | Q8KRP0 | *Fjoh_0890* | 2.31 | 39.28 | (1.23) | (12.76) |
| PorU | SPI | A5FJM8 | *Fjoh_1556* | nd | - | 1.51 | 33.50 |
| RemI | SPI | A5FF07 | *Fjoh_3194* | 2.72 | 23.89 | 1.66 | 23.81 |
| GldD | SPII | A5FJP0 | *Fjoh_1540* | nd | - | nd | - |

**Supplementary Table S4**. **LolA and LolB homologs in several Bacteroidetes species**.
Homologs identified by DELTA Blast (1) search using LolA1 (WP_081432686.1), LolA2 (WP_012023170.1), LolA3 (WP_012022695.1), LolB1 (WP_012023151.1) and LolB2 (WP_012023169.1) sequences of *F. johnsoniae* as queries (E value ≤ 0.001).

| Query | Species | RefSeq assembly | Protein accession | % ID | E value |
|---|---|---|---|---|---|
| LolA1 | *Bacteroides fragilis* | GCF_000025985.1 | WP_005795926.1 | 20.4 | 1.11E-27 |
| LolB1 | *Bacteroides fragilis* | GCF_000025985.1 | WP_005784040.1 | 17.2 | 7.24E-17 |
| LolA1 | *Bacteroides ovatus* | GCF_001314995.1 | WP_004301819.1 | 22 | 1.22E-26 |
| LolA2 | *Bacteroides ovatus* | GCF_001314995.1 | WP_004323602.1 | 23 | 2.17E-23 |
| LolB1 | *Bacteroides ovatus* | GCF_001314995.1 | WP_004298193.1 | 16.5 | 6.00E-13 |
| LolB2 | *Bacteroides ovatus* | GCF_001314995.1 | WP_004324728.1 | 23.5 | 9.80E_8 |
| LolA1 | *Bacteroides thetaioataomicron* | GCF_014131755.1 | WP_011109203.1 | 22.5 | 2.05E-26 |
| LolB1 | *Bacteroides thetaioataomicron* | GCF_014131755.1 | WP_011108844.1 | 17.3 | 1.11E-13 |
| LolA1 | *Bergeyella zoohelcum* | GCF_000301075.1 | WP_002663348.1 | 28.5 | 2.13E_17 |
| LolB1 | *Bergeyella zoohelcum* | GCF_000301075.1 | WP_002664277.1 | 21.5 | 1.18E-29 |
| LolA1 | *Capnocytophaga canimorsus* | GCF_000220625.1 | WP_013997750.1 | 61.5 | 7.31E-30 |
| LolB1 | *Capnocytophaga canimorsus* | GCF_000220625.1 | WP_126321423.1 | 25.6 | 1.65E-42 |
| LolA1 | *Capnocytophaga canis* | GCF_000827555.1 | WP_042010073.1 | 57.5 | 1.3E-30 |
| LolB1 | *Capnocytophaga canis* | GCF_000827555.1 | WP_042009446.1 | 27.2 | 1.75E-42 |
| LolA1 | *Capnocytophaga cynodegmi* | GCF_000379185.1 | WP_018278573.1 | 58.4 | 1.17E-38 |
| LolB1 | *Capnocytophaga cynodegmi* | GCF_000379185.1 | WP_018278328.1 | 29.5 | 2.79E-55 |

| | | | | | |
|---|---|---|---|---|---|
| LolA1 | *Capnocytophaga gingivalis* | GCF_000174755.1 | EEK14233.1 and EEK14289.1 (fragments) | 62.5 (EEK14233.1) and 48.9 (EEK14289.1) | 4E-15 (EEK14233.1) and 3E-08 (EEK14289.1) |
| LolA2 | *Capnocytophaga gingivalis* | GCF_000174755.1 | WP_002666178.1 | 45.6 | 7.35E-29 |
| LolB1 | *Capnocytophaga gingivalis* | GCF_000174755.1 | WP_040359568.1 | 25.8 | 1.68E-34 |
| LolB2 | *Capnocytophaga gingivalis* | GCF_000174755.1 | WP_002666377.1 | 28.2 | 9.89E-16 |
| LolA1 | *Capnocytophaga ochracea* | GCF_000023285.1 | WP_015782582.1 | 58.8 | 1.94E-33 |
| LolA2 | *Capnocytophaga ochracea* | GCF_000023285.1 | WP_015782811.1 | 44.9 | 9.41E-28 |
| LolB1 | *Capnocytophaga ochracea* | GCF_000023285.1 | WP_015782004.1 | 31.5 | 1.25E-56 |
| LolB2 | *Capnocytophaga ochracea* | GCF_000023285.1 | WP_015782812.1 | 33.3 | 1.63E-22 |
| LolA1 | *Chitinophaga filiformis* | GCF_900102545.1 | WP_089836732.1 | 24.8 | 5.14E-36 |
| LolA2 | *Chitinophaga filiformis* | GCF_900102545.1 | WP_089838851.1 | 27.4 | 1.88E-33 |
| LolB1 | *Chitinophaga filiformis* | GCF_900102545.1 | WP_089834398.1 | 22 | 3.14E-43 |
| LolB2 | *Chitinophaga filiformis* | GCF_900102545.1 | WP_245705585.1 | 27.4 | 1.88E-33 |
| LolA1 | *Chitinophaga pinensis* | GCF_000024005.1 | WP_012793835.1 | 27.3 | 1.25E-36 |
| LolA2 | *Chitinophaga pinensis* | GCF_000024005.1 | WP_012789541.1 | 27.6 | 5.39E-36 |
| LolB1 | *Chitinophaga pinensis* | GCF_000024005.1 | WP_044217960.1 | 23.4 | 4.00E-44 |
| LolB2 | *Chitinophaga pinensis* | GCF_000024005.1 | WP_012789542.1 | 28.5 | 1.08E-17 |
| LolA1 | *Gramella forsetii* | GCF_000060345.1 | WP_011710052.1 | 54.9 | 6.98E-35 |
| LolA2 | *Gramella forsetii* | GCF_000060345.1 | WP_011709538.1 | 38 | 5.53E-33 |
| LolB1 | *Gramella forsetii* | GCF_000060345.1 | WP_011711102.1) | 41.5 | 4.58E-77 |
| LolA1 | *Croceibacter atlanticus* | GCF_000196315.1 | WP_041241137.1 | 59.6 | 8.93E-34 |

| | | | | | |
|---|---|---|---|---|---|
| LolA2 | *Croceibacter atlanticus* | GCF_000196315.1 | WP_013186535.1 | 48.8 | 2.09E-32 |
| LolB1 | *Croceibacter atlanticus* | GCF_000196315.1 | WP_041240893.1 | 37.6 | 1.77E-75 |
| LolB2 | *Croceibacter atlanticus* | GCF_000196315.1 | WP_013186534.1 | 51.2 | 8.71E-49 |
| LolA1 | *Cytophaga hutchinsonii* | GCF_000014145.1 | WP_011586060.1 | 30 | 3.17E-36 |
| LolA2 | *Cytophaga hutchinsonii* | GCF_000014145.1 | WP_011585492.1 | 30 | 1.3E-24 |
| LolB1 | *Cytophaga hutchinsonii* | GCF_000014145.1 | WP_041932091.1 | 26.4 | 2.04E-50 |
| LolB2 | *Cytophaga hutchinsonii* | GCF_000014145.1 | WP_011585491.1 | 31.4 | 9.43E-15 |
| LolA1 | *Flavobacterium columnare* | GCF_007990835.1 | WP_077225215.1 | 59.6 | 8.85E-34 |
| LolA2 | *Flavobacterium columnare* | GCF_007990835.1 | WP_014166421.1 | 59.6 | 8.31E-33 |
| LolA3 | *Flavobacterium columnare* | GCF_007990835.1 | WP_097609575.1 | 37.9 | 4.4E-34 |
| LolB1 | *Flavobacterium columnare* | GCF_007990835.1 | WP_097609630.1 | 41.3 | 2.78E-58 |
| LolB2 | *Flavobacterium columnare* | GCF_007990835.1 | WP_041253277.1 | 39.6 | 1.12E-36 |
| LolA1 | *Flavobacterium johnsoniae* | GCF_034479105.1 | WP_081432686.1 | 100 | 2.37E-41 |
| LolA2 | *Flavobacterium johnsoniae* | GCF_034479105.1 | WP_012023170.1 | 100 | 4.76E-135 |
| LolA3 | *Flavobacterium johnsoniae* | GCF_034479105.1 | WP_012022695.1 | 100 | 1.78E-138 |
| LolB1 | *Flavobacterium johnsoniae* | GCF_034479105.1 | WP_012023151.1 | 100 | 1.78E-104 |
| LolB2 | *Flavobacterium johnsoniae* | GCF_034479105.1 | WP_012023169.1 | 100 | 4.76E-135 |

| | | | | | |
|---|---|---|---|---|---|
| LolA1 | *Elizabethkingia meningoseptica* | GCF_000367325.1 | WP_019051346.1 | 27.1 | 8.2E-21 |
| LolB1 | *Elizabethkingia meningoseptica* | GCF_000367325.1 | WP_026149339.1 | 26.8 | 1.43E-39 |
| LolA1 | *Flavobacterium psychrophilum* | GCF_900101925.1 | WP_011963703.1 | 64.8 | 5.64E-36 |
| LolA2 | *Flavobacterium psychrophilum* | GCF_900101925.1 | WP_011964352.1 | 75.2 | 4.85E-35 |
| LolB1 | *Flavobacterium psychrophilum* | GCF_900101925.1 | WP_011964528.1 | 43.3 | 1.2E-67 |
| LolB2 | *Flavobacterium psychrophilum* | GCF_900101925.1 | WP_034099413.1 | 64.6 | 2.73E-78 |
| LolA1 | *Flavobacterium succinicans* | GCF_000611675.1 | WP_024980104.1 | 82.7 | 3.01E-38 |
| LolB1 | *Flavobacterium succinicans* | GCF_000611675.1 | WP_024980583.1 | 56.8 | 1.28E-70 |
| LolA1 | *Flexibacter flexilis* | GCF_900112255.1 | WP_091512919.1 | 17.8 | 1.89E-28 |
| LolB1 | *Flexibacter flexilis* | GCF_900112255.1 | WP_221405328.1 | 22.2 | 4.81E-27 |
| LolA1 | *Kordia algicida* | GCF_000154725.1 | WP_040559877.1 | 54.2 | 1.98E-35 |
| LolA3 | *Kordia algicida* | GCF_000154725.1 | WP_007093428.1 | 21.3 | 1.58E-8 |
| LolB1 | *Kordia algicida* | GCF_000154725.1 | WP_238528710.1 | 35.1 | 5.24E-63 |
| LolA1 | *Polaribacter irgensii* | GCF_000153225.1 | WP_004570735.1 | 31 | 3.78E-24 |
| LolB1 | *Polaribacter irgensii* | GCF_000153225.1 | WP_004569654.1 | 29.6 | 5.84E-53 |
| LolA1 | *Porphyromonas gingivalis* | GCF_000010505.1 | WP_012457548.1 | 20.7 | 2.13E-14 |
| LolA3 | *Porphyromonas gingivalis* | GCF_000010505.1 | WP_004584472.1 | 25 | 5.45E-12 |
| LolB1 | *Porphyromonas gingivalis* | GCF_000010505.1 | WP_012457941.1 | 17.9 | 1.05E-35 |
| LolA1 | *Prevotella intermedia* | GCF_000439065.1 | WP_028905298.1 | 16.3 | 1.48E-20 |
| LolA3 | *Prevotella intermedia* | GCF_000439065.1 | WP_004367589.1 | 18.6 | 4.73E-11 |

| | | | | | |
|---|---|---|---|---|---|
| LolB1 | *Prevotella intermedia* | GCF_000439065.1 | WP_028905940.1 | 12.6 | 1.16E-28 |
| LolA1 | *Prevotella melaninogenica* | GCF_000144405.1 | WP_013265419.1 | 17.8 | 9.11E-22 |
| LolB1 | *Prevotella melaninogenica* | GCF_000144405.1 | WP_013264029.1 | 15.2 | 6.31E-29 |
| LolA1 | *Riemerella anatipestifer* | GCF_000183155.1 | WP_004919104.1 | 31 | 2.35E-18 |
| LolB1 | *Riemerella anatipestifer* | GCF_000183155.1 | WP_004916613.1 | 25.5 | 4.92E-31 |
| LolA1 | *Sphingobacterium mizutaii* | GCF_007990895.1 | WP_093100220.1 | 27.9 | 1.91E-27 |
| LolB1 | *Sphingobacterium mizutaii* | GCF_007990895.1 | WP_236736514.1 | 24.7 | 1.15E-42 |
| LolA1 | *Sporocytophaga myxococcoides* | GCF_000426725.1 | WP_028982018.1 | 26.6 | 1.72E-41 |
| LolA2 | *Sporocytophaga myxococcoides* | GCF_000426725.1 | WP_051312992.1 | 32.4 | 7.73E-36 |
| LolA3 | *Sporocytophaga myxococcoides* | GCF_000426725.1 | WP_028981309.1 | 21.3 | 1.16E-9 |
| LolB1 | *Sporocytophaga myxococcoides* | GCF_000426725.1 | WP_028981195.1 | 25.4 | 3.04E-34 |
| LolB2 | *Sporocytophaga myxococcoides* | GCF_000426725.1 | WP_028979568.1 | 34.6 | 9.67E-24 |
| LolA1 | *Xanthomarina gelatinilytica* | GCF_000348685.1 | WP_007646658.1 | 59.1 | 4.35E-39 |
| LolA2 | *Xanthomarina gelatinilytica* | GCF_000348685.1 | WP_007647468.1 | 48.2 | 2.63E-29 |
| LolB1 | *Xanthomarina gelatinilytica* | GCF_000348685.1 | WP_007647364.1 | 30.4 | 8.9E-53 |

| | | | | | |
|---|---|---|---|---|---|
| LolB2 | *Xanthomarina gelatinilytica* | GCF_000348685.1 | WP_007647467.1 | 40.6 | 1.66E-35 |
| LolA1 | *Zobellia galactanivorans* | GCF_000973105.1 | WP_013995938.1 | 55.7 | 1.05E-33 |
| LolA2 | *Zobellia galactanivorans* | GCF_000973105.1 | WP_013993418.1 | 42.4 | 1.58E-29 |
| LolB1 | *Zobellia galactanivorans* | GCF_000973105.1 | WP_013993148.1 | 35.1 | 7.35E-66 |
| LolB2 | *Zobellia galactanivorans* | GCF_000973105.1 | WP_013993417.1 | 36.5 | 4.1E-38 |

**Table S5 Bacterial strains used in this study.**

| Strain | Genotype and/or description | Reference |
|---|---|---|
| ***Flavobacterium johnsoniae*** | | |
| WT | *F. johnsoniae* UW101 | (2) |
| *ΔgldJ* | Deletion of *Fjoh_1557* | This study and (3) |
| *ΔgldJ-548* | Deletion of *Fjoh_1557* residues 549-561 | This study and (4) |
| *ΔlolA1* | Deletion of *Fjoh_2111* | This study |
| *ΔlolA2* | Deletion of *Fjoh_1085* | This study |
| *ΔlolB1* | Deletion of *Fjoh_1066* | This study |
| *ΔlolB2* | Deletion of *Fjoh_1084* | This study |
| *ΔlolA3* | Deletion of *Fjoh_0605* | This study |
| *ΔlolA1ΔlolB1* | Deletion of *Fjoh_2111* and *Fjoh_1066* | This study |
| *ΔlolA2ΔlolB2* | Deletion of *Fjoh_1084-1085* | This study |
| *ΔlolA1ΔlolA2 ΔlolA3ΔlolB1ΔlolB2* | Deletion of *Fjoh_2111, Fjoh_1066*, *Fjoh_1084-1085*, and *Fjoh_0605* | This study |
| ***Escherichia coli*** | | |
| Top10 | F-mcrAΔ(mrr-hsdRMS-mcrBC) ϕ80lacZΔM15ΔlacX74 recA1araD139Δ(araleu)7697 galU galK rpsL endA1 nupG; $Sm^R$ | Invitrogen |

| | | |
|---|---|---|
| MT607 | *pro-82 thi-I hsdR17 (r-m+) supE44 recA56* | Received from R. Hallez lab and (5) |
| MG1655 mini-λ-Tet | MG1655 *mini-λ-Tet* | (6) |
| ***Capnocytophaga canimorsus*** | | |
| Cc5 | Wild type (BCCM-LMG 28512) | (7) |

**Table S6 Plasmids used in this study.**

| **Description** | **Reference** | |
|---|---|---|
| **Vectors** | | |
| pYT354 | Suicide vector carrying *sacB*; MCS of pBC SK+ cloned into pYT313; $Amp^R$ ($Ery^R$) | (8) |
| pCP23 | ColE1 ori; (pCP1 ori); $Amp^R$ ($Tet^R$);*E. coli-F. johnsoniae* shuttle plasmid | (9) |
| pCP23-P*ermF* | pCP23 with P*ermF* promoter and MCS from pMM47.A cloned into BamHI and PstI restriction sites | This study |
| pBAD33 | pACYC *ori*; $Cm^R$. Low copy *E. coli* expression plasmid with arabinose inducible promoter | (10) |
| pMM47.A | ColE1 *ori*; (pCC7 *ori*); $Amp^R$; ($Cfx^R$). *E. coli-C. canimorsus* expression shuttle plasmid with *ermF* promoter | (7) |
| pKD4 | Template plasmid for gene disruption in *E. coli* with the $Kan^R$ gene flanked by FRT sites | Received from R. Hallez lab and (11) |
| **Suicide plasmids** | | |
| pYT313-*gldJ*-KO | Deletion of *Fjoh_1557.* | Received form Ben Berks' lab |
| pYT313-*gldJ-548* | Deletion of amino acids 548-561 of *Fjoh_1557.* | Received form Ben Berks' lab |
| pYT354-*lolA1*-KO | Deletion of *Fjoh_2111.* Upstream and downstream regions of *Fjoh_2111* amplified with oligonucleotides 8577 and 8503 and 8504 and 8578 respectively from gDNA and cloned sequentially into pYT354 using ApaI, XhoI and SpeI restriction sites. | This study |

| | | |
|---|---|---|
| pYT354-*lolA2*-KO | Deletion of *Fjoh_1085.* Upstream and downstream regions of *Fjoh_1085* amplified with oligonucleotides 8593 and 8594 and 8595 and 8596 respectively from gDNA and cloned sequentially into pYT354 using SphI, XhoI, and SpeI restriction sites. | This study |
| pYT354-*lolA3*-KO | Deletion of *Fjoh_0605.* Upstream and downstream regions of *Fjoh_0605* amplified with oligonucleotides TD3 and TD4 and TD5 and TD6 respectively from gDNA and cloned into pYT354 (amplified with oligonucleotides TD1 and TD2) by Gibson assembly. | This study |
| pYT354-*lolB1*-KO | Deletion of *Fjoh_1066.* Upstream and downstream regions of *Fjoh_1066* amplified with oligonucleotides 8619 and 8620 and 8621 and 8622 respectively from gDNA and cloned sequentially into pYT354 using SphI, XhoI and SpeI restriction sites. | This study |
| pYT354-*lolB2*-KO | Deletion of *Fjoh_1084.* Upstream and downstream regions of *Fjoh_1084* amplified with oligonucleotides 8625 and 8626 and 8627 and 8628 respectively from gDNA and cloned sequentially into pYT354 using SphI, XhoI and BamHI restriction sites. | This study |
| pYT354-*lolA2*-*lolB2*-KO | Deletion of *Fjoh_1084* and *Fjoh_1085.* Upstream region of *Fjoh_1085* and downstream region of *Fjoh_1084* amplified with oligonucleotides 8593 and 8594 and 8627 and 8628 respectively from gDNA and cloned sequentially into pYT354 using SphI, XhoI and BamHI restriction sites. | This study |
| pYT354-*Ccan_17050*-KO | Deletion of *C. canimorsus Ccan_17050* (*lolB*). Upstream and downstream regions of *Ccan_17050* amplified with oligonucleotides 8631 and 8632 and 8633 and 8634 respectively from gDNA. The two fragments were fused by PCR using oligonucleotides 8631 and 8634 and cloned into pYT354 using SphI and XhoI restriction sites. | This study |
| **Expression plasmids** | | |
| pCP23-P*ermF*-*siaC* | WT sialidase cloned into pCP23-P*ermF*. | This study |
| pCP23-P*ermF*-LES-*siaC* | Sialidase harboring the *F. johnsoniae* LES sequence (SDDFE) cloned into pCP23-P*ermF*. | This study |

| | | |
|---|---|---|
| pCP23-P*ermF-lolA1* | Full length *Fjoh_2111* amplified with oligonucleotides 8499 and 8500 and cloned into pCP23-P*ermF* using NcoI and XhoI restriction sites. | This study |
| pCP23-P*ermF-lolB1* | Full length *Fjoh_1066* amplified with oligonucleotides 8635 and 8636 and cloned into pCP23-P*ermF* using NcoI and XhoI restriction sites. | This study |
| pCP23-P*ermF-lolA1*-P*ermF-lolB1* | Full length *Fjoh_1066* amplified with *ermF* promoter from pCP23-P*ermF-lolB1* with oligonucleotides 8644 and 8645 and cloned into pCP23-P*ermF-lolA1* using XhoI and SpeI restriction sites. | This study |
| pCP23-P*ermF-lolA* *(E. coli)* | Full length *lolA* amplified from *E. coli* gDNA with oligonucleotides 8591 and 8592 and cloned into pCP23-P*ermF* using NcoI and XbaI restriction sites. | This study |
| pCP23-P*ermF-lolB* (*E. coli*) | Full length *lolB* amplified *from E. coli* gDNA with oligonucleotides 5434 and 5435 and cloned into pCP23-P*ermF* using NcoI and XbaI restriction sites. | This study |
| pCP23-P*ermF-lolA-lolB* (*E. coli*) | Full length *lolB* amplified from *E. coli* gDNA with oligonucleotides 8638 and 8639 and cloned into pCP23-P*ermF-lolA* (*E. coli*) using XbaI and SpeI restriction sites. | This study |
| pCP23-P*ermF-lolA* (*C. canimorsus*) | Full length *lolA* (*Ccan_16490*) amplified from *C. canimorsus* Cc5 gDNA with primers 7203 and 7204 and cloned into pCP23-P*ermF* using NcoI and XbaI restriction sites. | This study |
| pCP23-P*ermF-lolB* (*C. canimorsus*) | Full length *lolB* (*Ccan_17050*) amplified from *C. canimorsus* Cc5 gDNA with primers 8687 and 8688 and cloned into pCP23-P*ermF* using NcoI and XhoI restriction sites. | This study |
| pCP23-P*ermF-lolB1L74E-His* | First fragment of *Fjoh_1066* was amplified with primers 8635 and TD12 while second fragment was amplified with TD11 and 8850 (no stop codon). The two fragments were fused by PCR using primers 8635 and 8850 and cloned into pCP23-P*ermF* using NcoI and XhoI restriction sites. | This study |

| | | |
|---|---|---|
| pCP23-*lolB1Δ73-76-His* | First fragment of *Fjoh_1066* was amplified with primers 8635 and TD18 while second fragment was amplified with TD17 and 8850 (no stop codon). The two fragments were fused using primers 8635 and 8850 and cloned into pCP23-P*ermF* using NcoI and XhoI restriction sites. | This study |
| pCP23-*lolB1Δ223-243-His* | *Fjoh_1066* was amplified with primers 8635 and TD25 (no stop codon) and cloned into pCP23-P*ermF* using NcoI and XhoI restriction sites. | This study |
| pCP23-P*ermF-mlolB1* | Full length *Fjoh_1066* with C17G mutation amplified with primers 8643 and 8688 and cloned into pCP23-P*ermF* using NcoI and XhoI restriction sites. | This study |
| pBAD33-*lolA* | Full length *lolA* with its RBS amplified from *E. coli* gDNA with primers 8660 and 8592 and cloned into pBAD33 using KpnI and XbaI restriction sites. | This study |
| pBAD33-*lolB* | Full length *lolB* with RBS amplified from *E. coli* gDNA with primers 8661 and 5435 and cloned into pBAD33 using KpnI and XbaI restriction sites. | This study |
| pBAD33-*lolA1* | Full length *Fjoh_2111* amplified with primers 8663 and 8651 and cloned into pBAD33 using KpnI and XbaI restriction sites. | This study |
| pBAD33-*lolA2* | Full length *Fjoh_1085* amplified with primers 8665 and 8655 and cloned into pBAD33 KpnI and XbaI restriction sites. | This study |
| pBAD33-*lolB1* | Full length *Fjoh_1066* amplified with oligonucleotides 8664 and 8653 and cloned into pBAD33 using KpnI and XbaI restriction sites. | This study |
| pBAD33-*lolB2* | Full length *Fjoh_1084* amplified with primers 8666 and 8657 and cloned into pBAD33 using KpnI and XbaI restriction sites. | This study |

| | | |
|---|---|---|
| pBAD33-*lolA1lolB1* | Full length *Fjoh_1066* amplified with primers 8667 and 8659 and cloned into pBAD33-*lolA1* using XbaI and SphI restriction sites. | This study |
| pBAD33-*lolA2lolB2* | Full length *Fjoh_1084-85* amplified with primers 8666 and 8655 and cloned into pBAD33 using KpnI and XbaI restriction sites. | This study |
| pMM47-*lolA1* | Full length *Fjoh_2111* amplified with oligonucleotides 8499 and 8500 and cloned into pMM47.A using NcoI and XhoI restriction sites. | This study |
| pMM47-*lolB1* | Full length *Fjoh_1066* amplified with oligonucleotides 8635 and 8636 and cloned into pMM47.A using NcoI and XhoI restriction sites. | This study |
| pFL63 | pFL63 expressing *C. canimorsus lolA Ccan_16490*; $Tet^R$ | |

**Table S7 Oligonucleotides used in this study.**

| Name | Sequence 5'-3' |
|---|---|
| 5434 | CATGCCATGGGACCCCTGCCCGATTTT |
| 5435 | GCTCTAGATTATTTCACTATCCAGTTATCC |
| 7203 | CCCCATGGGGAAAAAGATACTATTGTTAATATC |
| 7204 | GGTCTAGATTATAGTTCTGAAATATAGTATCC |
| 8499 | CATACCATGGGAAACAAAATTAATCCAATCATG |
| 8500 | CCGCTCGAGTTAATCTAATTTATTGATGTAG |
| 8504 | CCGCTCGAGGTGCCGACTACAGGAAAAGAC |
| 8578 | TGCACTACTAGTAGCAGCAGAAATACCAGTTG |
| 8591 | CATGCCATGGGAAAAAAAAATTGCCATCACCTGT |
| 8592 | GCTCTAGACTACTTACGTTGATCATCTACCG |
| 8593 | CATGCATGCTATCTAATTCTTTCGGATTTGGAGG |
| 8594 | CCGCTCGAGGTGCTTAATGTTTTTCCCAC |
| 8595 | CCGCTCGAGCGACAAAGAATTTACGATTTCG |
| 8596 | GCACTACTAGTATAGTAATCGCTGTCCAGAATATC |
| 8619 | CATGCATGCGAACAGCCTGAAGCTTTTGG |
| 8620 | CCGCTCGAGCTTAATTTCTCCTTACTTCTTTTG |
| 8621 | CCGCTCGAGTACCAAGCGGTTATAAAAAAG |
| 8622 | GCACTACTAGTCTCACTTCACCTTGCAGAATATC |
| 8625 | CATGCATGCGCGGTGCCGTTTATGGTGAAG |
| 8626 | CCGCTCGAGCCAAAACGATTGCCAGAAAGC |
| 8627 | CCGCTCGAGCAAGTCTGAGTAGAATATTAAAC |
| 8628 | CGGGATCCGTAACCCGTAATAGCCGGAAC |
| 8631 | CATGCATGCAATGTTGATGCTCGTGATGG |
| 8632 | ACGGAATACGGTACGGTGCGGTGTATTTAACTGATGGTAA |
| 8633 | TTAAATACACCGCACCGTACCGTATTCCGTCGGG |
| 8634 | CCGCTCGAGATTGCTTTATGTAAAGCATACG |

| | |
|---|---|
| 8635 | CATACCATGGGAAAAAAATATATTATAATAG |
| 8636 | CCGCTCGAGTTACTTAATTAAAACTTTTTTATAAC |
| 8638 | GCTCTAGAATGCCCCTGCCCGATTTTCGTC |
| 8639 | GCACTACTAGTTTATTTCACTATCCAGTTATCC |
| 8643 | CATACCATGGGAAAAAAATATATTATAATAGTATTAATATCGGTTTTTGTGGTTTCAGGTAAATC |
| 8651 | GGGGTACCATGAAAAAATATATTATAATAG |
| 8653 | GCTCTAGATTACTTAATTAAAACTTTTTTATAAC |
| 8655 | GCTCTAGATTAATTAGTAAAAACTGAATC |
| 8657 | GCTCTAGACTACTCAGACTTGAAATAATTC |
| 8659 | ACATGCATGCTTACTTAATTAAAACTTTTTTATAAC |
| 8660 | GGGTACCCGGGAGTGACGTAATTTGAG |
| 8661 | GGGGTACCAGGGTTATAACTGCAACGTATC |
| 8663 | GGGGTACCAGGAGGACAGCTATGAACAAAATTAATCCAATC |
| 8664 | GGGGTACCAGGAGGACAGCTATGAAAAAATATATTATAATAG |
| 8665 | GGGGTACCAGGAGGACAGCTATGAAAACTAAAATAGCTCTAC |
| 8666 | GGGGTACCAGGAGGACAGCTATGCAAAAATCGACGATTCAG |
| 8667 | GCTCTAGAAGGAGGACAGCTATGAAAAAATATATTATAATAG |
| 8675 | ATTATTAGCCTGGAATAGAGAGTAGAGGGAACTCCCGATGTGTAGGCTGGAGCTGCTTC |
| 8677 | CTTGAACATAGACGATAGCGGACGGTAACGCTAGCATTAGTGTAGGCTGGAGCTGCTTC |
| 8679 | TCCGAAAAATCGAGCGACAGATTGCTCACTCAGGTGCCTCATATGAATATCCTCCTTA |
| 8680 | CAGATTAAGTTTTGCCGGAGAGGGCCACTGTGTCCGCATCATATGAATATCCTCCTTA |
| 8687 | CATACCATGGGAAAATACCTTTTTCTGAAAATAC |
| 8688 | CCGCTCGAGTTAATTATTGATTGCTTTCG |
| 8850 | CCGCTCGAGCTTAATTAAAACTTTTTTATAACCGC |
| TD1 | GAATTGAAGAAGACGGTATTCGGTATCGATAAGCTTGATATCGAATTCCTGCAGCCC |
| TD2 | TATCTTTCTGTTACGGTTATTTCTTTTGTAAATGTCGACCTCGAGGGGGGGCC |
| TD3 | GGCCCCCCCTCGAGGTCGACATTTACAAAAGAAATAACCGTAACAGAAAGATATT |
| TD4 | TCTCGGCCGGAACAATTTGTTTTTTAGCCGATGCTTTTATGGTGTTGTATTTGTC |
| TD5 | GACAAATACAACACCATAAAAGCATCGGCTAAAAAACAAATTGTTCCGGCCGAGA |

| | |
|---|---|
| TD6 | GGGCTGCAGGAATTCGATATCAAGCTTATCGATACCGAATACCGTCTTCTTCAATTC |
| TD11 | ACAGATTTTAATAAGCGTTAGATTCGAGGGAATTACAATGGCAAAAGCTTTAA |
| TD12 | TTAAAGCTTTTGCCATTGTAATTCCCTCGAATCTAACGCTTATTAAAATCTGT |
| TD17 | CAAACAGATTTTAATAAGCGTTAGAACAATGGCAAAAGCTTTAATAACAC |
| TD18 | GTGTTATTAAAGCTTTTGCCATTGTTCTAACGCTTATTAAAATCTGTTTG |
| TD25 | CCGCTCGAGTGAAATGTTATTGTAATTCAGATTA |

**References**

1. Boratyn GM, Schaffer AA, Agarwala R, Altschul SF, Lipman DJ, Madden TL. Domain enhanced lookup time accelerated BLAST. Biol Direct. 2012;7:12.
2. McBride MJ, Xie G, Martens EC, Lapidus A, Henrissat B, Rhodes RG, et al. Novel features of the polysaccharide-digesting gliding bacterium Flavobacterium johnsoniae as revealed by genome sequence analysis. Appl Environ Microbiol. 2009;75(21):6864-75.
3. Braun TF, McBride MJ. Flavobacterium johnsoniae GldJ is a lipoprotein that is required for gliding motility. J Bacteriol. 2005;187(8):2628-37.
4. Johnston JJ, Shrivastava A, McBride MJ. Untangling Flavobacterium johnsoniae Gliding Motility and Protein Secretion. J Bacteriol. 2018;200(2).
5. Aneja P, Charles TC. Poly-3-hydroxybutyrate degradation in Rhizobium (Sinorhizobium) meliloti: isolation and characterization of a gene encoding 3-hydroxybutyrate dehydrogenase. J Bacteriol. 1999;181(3):849-57.
6. Court DL, Swaminathan S, Yu D, Wilson H, Baker T, Bubunenko M, et al. Mini-lambda: a tractable system for chromosome and BAC engineering. Gene. 2003;315:63-9.
7. Mally M, Cornelis GR. Genetic tools for studying Capnocytophaga canimorsus. Appl Environ Microbiol. 2008;74(20):6369-77.
8. Zhu Y, Thomas F, Larocque R, Li N, Duffieux D, Cladiere L, et al. Genetic analyses unravel the crucial role of a horizontally acquired alginate lyase for brown algal biomass degradation by Zobellia galactanivorans. Environ Microbiol. 2017;19(6):2164-81.
9. Agarwal S, Hunnicutt DW, McBride MJ. Cloning and characterization of the Flavobacterium johnsoniae (Cytophaga johnsonae) gliding motility gene, gldA. Proc Natl Acad Sci U S A. 1997;94(22):12139-44.
10. Guzman LM, Weiss DS, Beckwith J. Domain-swapping analysis of FtsI, FtsL, and FtsQ, bitopic membrane proteins essential for cell division in Escherichia coli. J Bacteriol. 1997;179(16):5094-103.
11. Datsenko KA, Wanner BL. One-step inactivation of chromosomal genes in Escherichia coli K-12 using PCR products. Proc Natl Acad Sci U S A. 2000;97(12):6640-5.
